# The structure of the native cardiac crossbridge in the rigor state

**DOI:** 10.64898/2026.01.27.702119

**Authors:** Cristina M. Risi, Tyler Nguyen, Betty Belknap, Howard D. White, Jose R. Pinto, P. Bryant Chase, Vitold E. Galkin

## Abstract

Cardiac contraction is driven by double-headed myosin cycling on cardiac thin filaments, where troponin-tropomyosin regulates myosin access to actin. Prior research used a single-headed myosin bound to bare actin, thereby limiting insight into coordination between myosin heads and the influence of troponin-tropomyosin on actomyosin interactions. Here, we report a high-resolution structure of the native cardiac rigor cross-bridge formed by heavy meromyosin bound to the thin filament. We show that direct communication between the two bound heads, uneven interactions between the heads and tropomyosin, and spatial constraints imposed by troponin govern myosin placement along the thin filament. Additionally, the two heads display non-equivalent motor-light chain interactions, yielding distinct lever-arm conformations indicative of asymmetric intramolecular strain. Together, these findings provide a structural framework for how the two myosin heads coordinate and how the components of the thin filament are integrated into force generation by active cross-bridges.

## INTRODUCTION

Cross-bridges between actin and myosin form ubiquitously in eukaryotic cells and mediate muscle contraction, intracellular cargo transport, and cytoskeletal remodeling via a mechanochemical cycle that couples ATP hydrolysis and filamentous actin (F-actin) binding to generate force (reviewed in^1,2^). While the steps involved in the ATPase cycle are the same for evolutionarily distant myosins, the rates and equilibrium constants of those steps are tailored for the functional role of each myosin^3,4^. Class II myosins found in skeletal and cardiac striated muscles exist as dimers composed of two heads connected by a long α-helical coiled-coil tail^5^ that allows them to assemble into filaments (thick filaments)^6^. Each head comprises two heavy chains and a pair of essential (ELC) and regulatory light chains (RLC)^1,7^ (Fig. 1A). Controlled proteolysis of cardiac myosin allows for isolation of its three main regions, widely used in physiological and structural studies – a so-called subfragment 1 (S1-head) comprised of 847 N-terminal residues and the two light chains, subfragment 2 (S2) stretching from 848 to 1216, and light meromyosin (LMM) that comprises the C-terminal residues 1218-1936^8^ (Fig. 1A). LMM is responsible for thick filament formation, while S2, together with the two S1 heads, forms a two-headed heavy meromyosin molecule (HMM) that does not polymerize into filaments (Fig. 1A) and can be used to study the interaction of individual double-headed myosin II fragments with F-actin.

**Figure 1.**
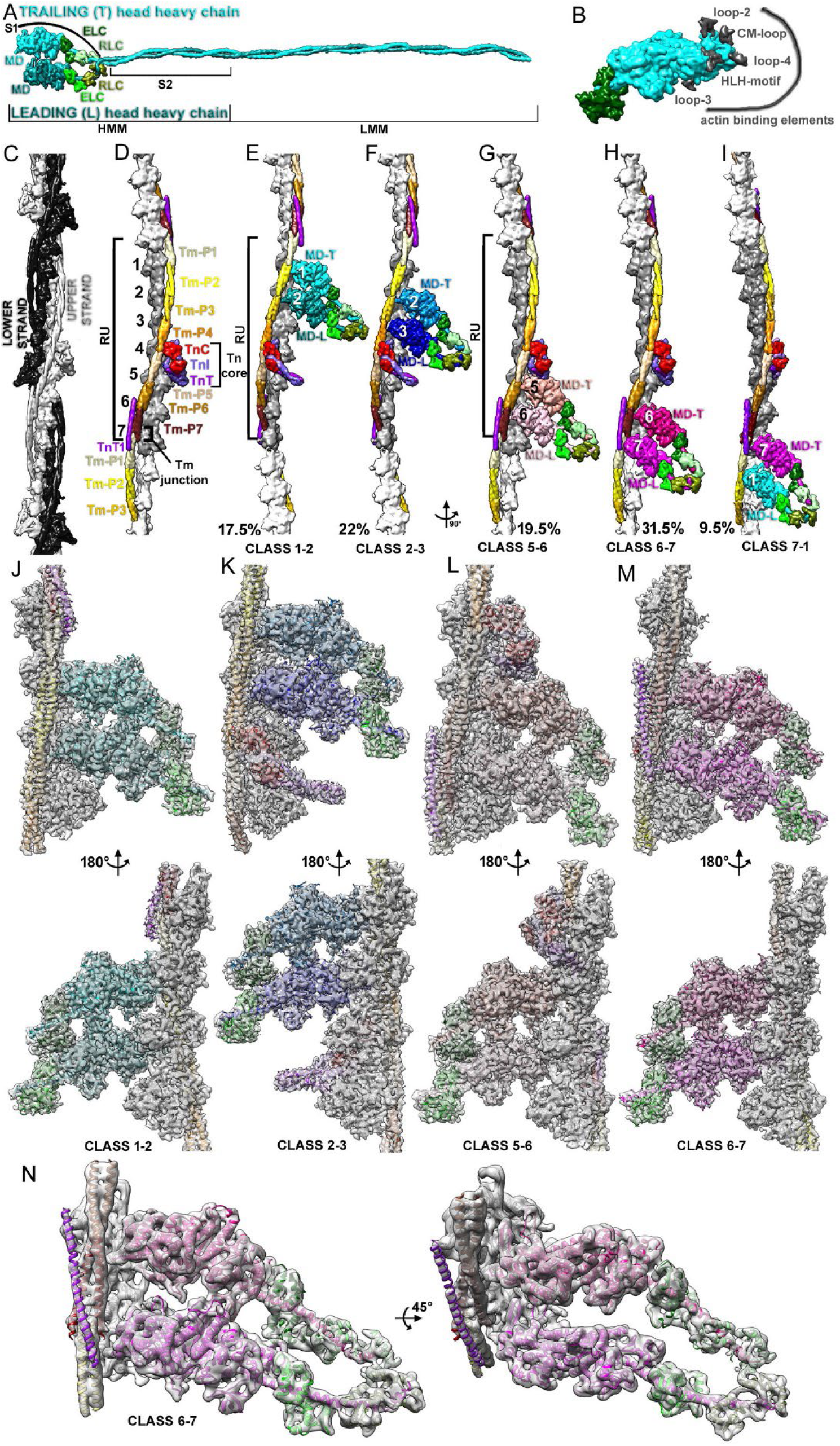
Overview of the native cardiac cross-bridge structure. (A) Two myosin molecules that form a double-headed molecule are shown in cyan and dark cyan, while associated light chains are shown in different shades of green. Different proteolytic fragments of myosin are marked with black brackets. (B) Positions of actin and Tm binding elements (dark gray) in the myosin motor domain. (C) TF is comprised of the upper and lower strands. (D) Structural elements of the TF regulatory unit (RU). The sequential numbers of actin protomers in the RU are shown in black. TnC is red, TnT is purple, and TnI is medium purple. Tm periods are colored in khaki, yellow, gold, orange, tan, sienna, and brown starting from period 1. (E-I) Cartoons of five structural classes of TF RUs with bound HMM and their associated frequencies (black numbers). Trailing and leading myosin heads are shown as cyan and dark cyan (Class 1-2), dodger blue and blue (Class 2-3), salmon and pink (Class 5-6), deep pink and magenta (Class 6-7), and magenta and cyan (Class 7-1). The ordinal numbers of actin molecules interacting with the two myosin heads are indicated. Myosin light chains are shown in four shades of green. (J-M) Actual 3D reconstructions of structural classes 1-2, 2-3, 5-6, and 6-7 (gray transparent surfaces) generated with RLC masked out and filtered to 3.9 Å, 4.2 Å, 4.0 Å, and 3.8 Å, respectively, are shown with their associated atomic models (colored atoms). The atomic models’ color codes are as in (E-H). (N) 3D reconstruction of Class 6-7 that includes RLCs and the myosin neck region, filtered to 6 Å resolution.

The actomyosin cross-bridge cycle relies on allosteric coupling between myosin’s actin-binding elements, ATP-binding active site, and the lever arm (reviewed in ^1^), which ensures that the biochemical state of the active site (i.e., nucleotide-free, ATP, ADP.Pi, or ADP-bound) determines the structural conformation of the myosin head and the lever arm (reviewed in^9^). The completion of myosin’s force-generating power stroke results in the formation of its high-affinity rigor interface with actin, involving all five actin-binding elements (Fig. 1B) interacting with actin. A subsequent power stroke occurs only after ATP binds to myosin, weakening the actin-myosin interaction and leading to a post-rigor state. Detached from cTF, the myosin then performs a recovery stroke by hydrolyzing ATP into ADP and Pi, forming a pre-power stroke (primed) state that traps ADP and P_i_ within the myosin active site until formation of the cross-bridge in the next round of the myosin cross-bridge cycle ^10,11^.

Our structural knowledge of the actomyosin complex primarily comes from helically averaged cryo-EM structures. Several rigor actomyosin complexes have been visualized by cryo-EM at near-atomic resolution, including cardiac myosin-S1 (3.8 Å; ^12,13^), myosin-V-S1 (3.0 Å; ^14^), and motor domains of skeletal myosin (5.2 Å; ^15^), myosin IIC (3.9 Å; ^16^), myosin VI (4.6 Å; ^17^), and myosin Ib (3.9 Å; ^18^). Comparison of these structures^1^ shows that, in the rigor state, the actomyosin interface is formed by interactions among specific structural regions on myosin referred to as the cardiomyopathy (CM) loop, the helix-loop-helix (HLH) motif, and loop-2 (Fig. 1B). Isoform-specific contributions from loop-3 and loop-4, which vary among myosin isoforms, can optimize actomyosin interaction. Recently, a 4.4 Å structure of primed myosin-V S1 bound to F-actin revealed that the HLH motif, loop-3, and part of loop-2 are involved in initial actomyosin interactions^19^. The main reason to use single-headed constructs for double-headed myosins (e.g., cardiac and skeletal myosins, myosin IIc, myosin V, and myosin VI) has been to obtain an actomyosin complex suitable for helical averaging using F-actin helical symmetry.

Dimerization of myosin II heads is not necessary for their motor function^20^, but it is essential for thick filament regulation and mechanical performance. The interaction between the two heads, folded back onto the S2 region (the interacting-heads motif, or IHM), is needed for the formation of the thick filament’s relaxed state^6,21^. On the other hand, striated muscle myosins require both heads to generate maximal force^22^. The binding of the two heads to F-actin is cooperative^23^. Time-resolved X-ray and mechanical measurements on isolated muscle cells show that the combined steric and mechanical coupling between the two myosin heads promotes the attachment of both heads to actin-based thin filaments during mechanical stretch^24^. Importantly, cryo-electron tomography (cryo-ET) of native cardiac murine sarcomeres reveals simultaneous interaction between both myosin heads and actin-based thin filaments^25,26^. Even though the interactions of the two striated myosin heads with actin are coupled, most of the structural work on muscle myosin’s interaction with F-actin has been done using single-headed fragments (e.g., S1 or motor domain)^12,15,27-29^. Indeed, this approach eliminated all the structural information relevant to communication between the two heads, leaving structural aspects of such communication unrevealed. As defined in previous structural work^26,30^, we will refer to the two non-equivalent myosin heads as the leading and trailing heads throughout the manuscript.

In cardiac muscle, the interaction between myosin heads and F-actin is tightly regulated to ensure coordinated cycles of contraction and relaxation. Therefore, myosin heads do not interact with bare F-actin but instead bind to the actin-based multiprotein complex—the cardiac thin filament (cTF) (Fig. 1C and D). The TF backbone is the actin filament; thus, the TF has two strands, which are not identical due to the presence of regulatory proteins and are called the upper and lower strands (Fig. 1C, grey and black surfaces). The TF regulatory apparatus is made up of the calcium-sensing troponin (Tn) complex (Fig. 1D, red, medium purple, and purple surfaces) and tropomyosin, which blocks actomyosin interactions in the absence of Ca^2+ 31-35^. Tm is comprised of a seven-quasi-equivalent-period repeat (Fig. 1D, Tm P1-P7) containing amino acid residues involved in binding to actin and formation of a continuous cable structure^36-39^. Recent cryo-EM studies of cardiac TF (cTF) have revealed the molecular details of its Ca^2+^-mediated allosterically coupled regulation^39-43^.

Ca^2+^ binding to the TnC subunit causes conformational changes in the TnC and TnI subunits (Fig. 1D red and medium purple surface, respectively), which release the Tm cable (Fig. 1D, Tm P1 to P7) from its “closed” position on actin subunits that sterically block myosin from binding to its sites on actin. Tn and Tm are spaced at intervals of every seven actin subunits to form repeating regulatory units (RUs) on each strand (Fig. 1D, large black bracket) connected via junction regions stabilized by the Tn-T region called TnT1 (Fig. 1D, small black bracket and purple surface). Since Tm covers seven actin subunits, its interactions with actin protomers are mediated through partially equivalent Tm repeats (Fig. 1D, marked P1 through P7). All the above suggest that the seven actins that make up the RU would not be identical in their interactions with myosin due to the regulatory elements of the TF (e.g., Tn-C, Tn-I, and Tn-T) and the unequal Tm repeats. This adds complexity to understanding actomyosin interactions between the two non-equivalent heads and the seven non-equivalent actin subunits.

To address these challenging questions, we obtained single-particle 3-dimensional (3D) reconstructions of cardiac myosin HMM molecules bound to all available actin protomers that comprise a native cardiac TF RU. We show that the interaction of the two myosin heads with the RU of the native cTF is multifaceted. It is affected by the cTF regulatory elements (e.g., TnC, TnT, TnI, and Tm), variable interactions between myosin loop-4 and Tm periods, and direct interactions between the leading and trailing heads via their loops 2 and 3. Further, we demonstrate that TnC and TnI together stabilize the myosin (e.g., fully activated) state of cTF. Our results have direct implications for understanding cardiac contraction and provide a structural framework for in silico molecular modeling.

## RESULTS

### Overview of the HMM interactions with the native cardiac TF

To elucidate the structure of a rigor cardiac cross-bridge, we used native porcine cardiac TFs and a porcine left ventricular cardiac HMM. Amino acid sequence alignment shows that the human and porcine TF proteins are highly identical (Fig. S1 and S2) – from 91.2% in TnT to 100% in actin. Cardiac myosin isoforms are also highly conserved between the two species (Fig. S3), sharing 97.7% identity in the heavy chain and over 95% identity in the light chains. Taken together, these findings make the porcine model an excellent model for the human heart.

Since we sought to understand how double-headed myosin molecules interact with non-equivalent (due to the presence of Tn subunits) actin protomers of the cardiac RU, we obtained cTFs sparsely decorated with HMM (Fig. S4A) using a substoichiometric concentration of HMM at high Ca^2+^ (pCa=4) to reduce the previously reported^44,45^ cooperative binding of myosin to TFs. Importantly, high Ca^2+^ is physiologically appropriate for cross-bridge formation during early systole. High-quality cTF fragments were manually extracted from 41,405 micrographs (Fig. S4B) and classified into 40 classes based on the positioning of HMM relative to the Tn complex (Fig. S5). Due to the TF’s two-stranded geometry and the length of cTF segments, multiple classes were redundant in terms of the HMM molecule’s position within the RU. Therefore, the obtained class frequencies were used to calculate the odds of HMM interacting with two adjacent actins within the RU, based on five possible HMM binding sites within the RU (Fig. 1E-I). These five class names (e.g., 1-2, 2-3, 5-6, 6-7, and 7-1) are used throughout to indicate the position of the HMM molecule within the RU, with actin subunits numbered from top to bottom of the RU (Fig. 1E-I). Classes 3-4 and 4-5 do not exist because of the Tn core located on actin subunit 4 (Fig. 1D, small bracket) that prevents myosin binding to actin 4. The comparison of the frequencies of the five classes (Fig. 1E-I, black numbers and Fig. S6A) revealed that HMM preferentially binds to actins 6-7 (31.5%), followed by about 20% frequency of binding to actins 2-3 and 5-6, while having the lowest odds of interaction with actins 7-1 (9.5%). Our data suggest that HMM does not differentiate between the two strands of the TF (Fig. S6A) because the frequencies of the five classes mentioned above are very similar across both strands. We hypothesize that the N-terminus of TnT interferes with the leading myosin head approaching actin 1 (see Fig. S6B-G and discussion).

The actual 3D reconstructions are shown for four classes (Fig. 1J-M), as class 7-1 was too small to yield a sufficiently detailed structure. The experimental details are provided in the supplementary information (Figs. S7–S15, Tables S1 and S2). Due to the high flexibility of the HMM tail, reconstructions included only ELCs (Fig. 1J-M), except for class 6-7, where we were able to determine the high-resolution structure that included ELCs (Fig. 1J), along with a slightly lower 6 Å structure of the two heads with both light chains visualized (Fig. 1N, and S18). The golden-standard Fourier Shell Correlation (FSC) 0.143 criterion yielded modest resolutions ranging from 4.3 Å for the largest class 6-7 to 4.8 Å for the smallest class 2-3 (Figs. S8A, S10A, S12A, and S14A) due to the noisy tails above and below the tightly masked region (Fig. S8B, red arrows). The FSC approach yields the most pessimistic resolution estimate because it requires splitting the set into two independent subsets, which naturally decreases the signal-to-noise ratio in the resulting two maps. RELION^46^ local resolution maps and local resolution estimation by a deep learning algorithm^47^ (Figs. S8B and C, S10B and C, S12B and C, and S14B and C) estimated the resolution at the actomyosin interface to be better (classes 1-2, 5-6, and 6-7) or close (class 2-3) to 4 Å, which is consistent with the molecular details seen in the density maps (Figs. S8D, S10D, S12D, and S14D). We also resolved the interface between the myosin motor domains and ELCs, as well as the structure of the Tn complex core at 4.2 Å resolution (Figs. S8B and S10B). At the reported resolution, we were able to unambiguously dock large and intermediate side chains of amino acids, yielding reliable models of the active rigor cross-bridge. Of note, the reported resolution here is similar to that of the previously published helically averaged cardiac actomyosin complex^12^ and the single particle reconstruction of the actomyosin V complex^19^.

### CM-loop and HLH-motif actin interactions are uniform across the two heads

The interface between the TF and cardiac myosin is made up of five myosin elements – the CM-loop, HLH-motif, loop-2, loop-3, and loop-4 (Fig. 1B). Our data show that the interactions among actin, the CM-loop, and the HLH-motif are conserved for both the trailing and leading heads across all four classes (Fig. 2). Specifically, the upper part of the CM-loop (residues V404, V406, and V411) creates a hydrophobic patch with actin residues A26, V30, and Y337 (Fig. 2B and C, black atoms). At the same time, K413 of the CM-loop forms ionic interactions with D25 and E334 of actin (Fig. 2B and C, dark grey atoms and grey arrows). Myosin V404L/M, V406M, and V411I, along with actin A26V, are linked to HCM in humans (Table S3).

**Figure 2.**
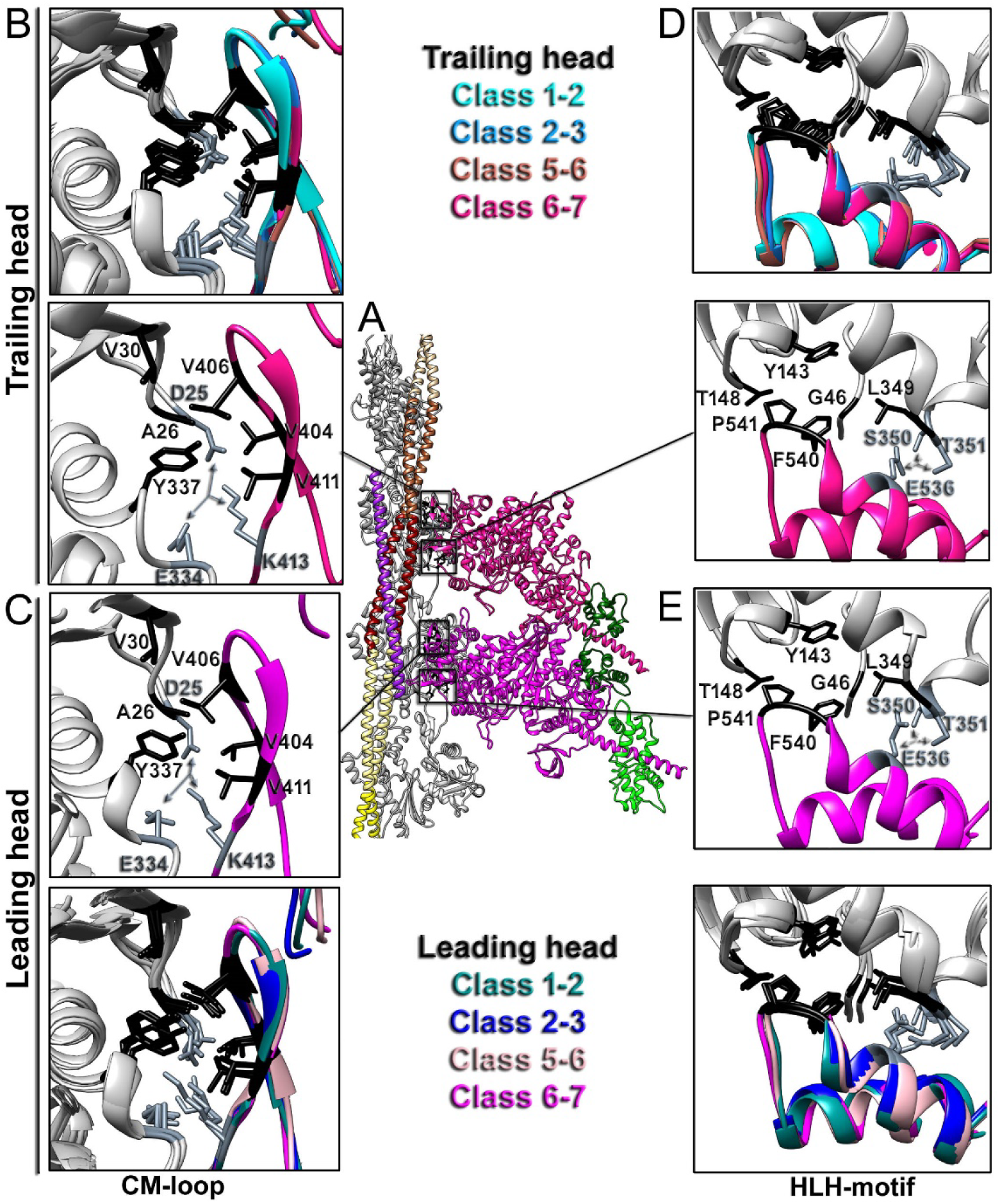
CM-loop and HLH-motif interactions with actin are uniform across the four structural classes. (A) Regions harboring the CM-loop (B-C) and HLH-motif (D-E) in the trailing (B, D) and leading (C, E) heads are marked with black rectangles and detailed in panels B-E. (B-E) Residues of actin and myosin that form hydrophobic interactions are shown in black, while those forming ionic bonds (gray arrows) are shown in slate gray. Superimposition of CM-loops or HLH-motifs from the four classes demonstrates uniform interactions. The color codes for the atomic models are shown in Fig. 1E-H.

HLH-motif residues F540 and P541, located at the tip of the loop, penetrate the hydrophobic cleft of F-actin to form a hydrophobic patch with residues G46, Y143, T148, and L349 of actin (Fig. 2D and E, black atoms). Additionally, an H-bond between HLH-motif E536 and S350 and T351 of actin (Fig. 2D and E, dark grey atoms and grey arrows) stabilizes the actomyosin interface. Cardiac myosin E536D and F540L missense mutations cause cardiomyopathies (Table S3).

Our data indicate that CM-loop and HLH-motif interactions with actin are uniform across all actins within the cTF RU, underscoring their universal role in establishing interactions between the upper (CM-loop) and lower (HLH-motif) domains of myosin and actin.

### Loop-2 facilitates communication between the two heads

The single-particle approach allowed us to visualize the structure and interactions of loop-2 (Fig. 3). Notably, the region G626-T629 of loop-2 was not clearly resolved in all density maps across the structural classes, suggesting weak interactions with the surrounding residues. We found that loop-2 interactions differ between the trailing (upper) and leading (lower) heads, stemming from the local environment of loop-2. While the trailing head’s loop-2 is exposed only to the actin surface (Fig. 3A, C, D, and G), the leading head’s loop-2 is near both actin and loop-3 of the trailing head above it (Fig. 3B, D, F, and H). The following interactions are identical between the two heads. The base of loop-2 is stabilized by the ionic interactions of K639/K640 with the negative tip of actin’s N-terminus and myosin E535 (Fig. 3A-H, straight black arrows). Consequently, myosin K639E leads to left ventricular non-compaction cardiomyopathy (Table S3). Loop-2 residues E632 and K635, located at its upper part, form salt bridges with the actin residues R95 and E100 (Fig. 3A-H, black curved arrows). On the other hand, loop-2 of the leading head in each of the four classes forms an interaction that is not observed in the trailing head – its K633 forms a salt bridge with loop-3 E574 of the trailing head (Fig. 3B, D, F, and H, red straight arrow). Additionally, interactions between the Class 5-6 trailing head loop-2 and its counterparts in Classes 1-2, 2-3, and 6-7 vary because the Tn core complex lies above loop-2 in Class 5-6 (Fig. 3E, purple ribbons). TnT pushes loop-2 away from the actin interface, establishing a new set of contacts—loop-2 E632 forms a salt bridge with K405 of the CM-loop, while loop-2 K635 forms an H-bond with TnT N266 (Fig. 3E, green curved arrows). The base of loop-2 interactions remains unaffected by TnT and involves ionic interactions between K639/K640 and the negative tip of actin’s N-terminus, as well as with myosin E535 (Fig. 3E, straight black arrows).

**Figure 3.**
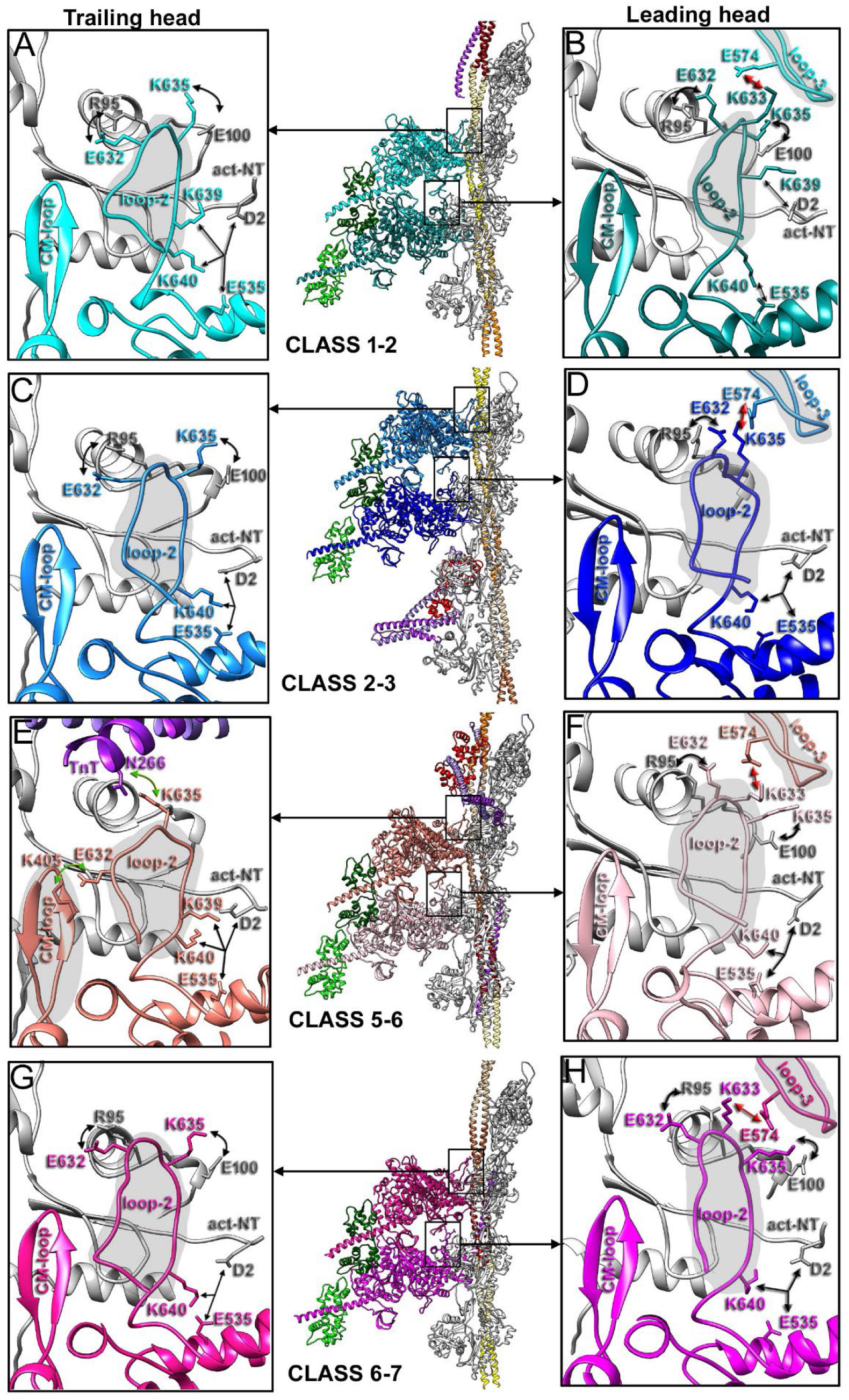
Interactions of loop-2 with actin and loop-3. Loop-2 interactions are compared between the trailing (A, C, E, and G) and leading (B, D, F, and H) heads for Classes 1-2 (A and B), 2-3 (C and D), 5-6 (E and F), and 6-7 (G and H). Salt bridges between loop-2 and actin, present in both heads, are marked with black curved arrows. Canonical ionic interactions between myosin E535, loop-2 K639/K640, and the N-terminus of actin (act-NT) are marked with straight black arrows. Ionic bonds between loop-2 and loop-3 observed only in the leading head are marked with straight red arrows. Finally, unique interactions between loop-2, TnT, and CM-loop are identified in the trailing head of Class 5-6 and are depicted with green curved arrows. The atomic models’ color codes are as shown in Fig. 1E-H.

To summarize, the leading head’s loop-2 interactions with loop-3 of the trailing head above contribute to cardiac myosin binding to cTF due to the formation of an additional energetically favorable ionic bond. In addition, the communication between the two heads via the leading head’s loop-2 and the trailing head’s loop-3 synchronizes their binding to cTF.

### The interaction between loop-4 and each Tm period is distinct

Tm consists of seven pseudo-repeats or periods (TmP) (Fig. 1D), each interacting with two neighboring actin molecules—TmP1 with actins 7 and 1, TmP2 with actins 1 and 2, TmP3 with actins 2 and 3, and so on (detailed in Fig. S16A). In this way, the last repeat (TmP7) contacts the first repeat (TmP1) of the adjacent Tm molecule on actin 7, forming a junction region (Fig. 1D, small black bracket). The amino acid sequences of these repeats are not 100% identical (Fig. S2B), which explains differences in their interactions with actin protomers reported here for the myosin state of Tm (M-state) on the actin surface (Fig. S16F). Specifically, TmP2 has the fewest contacts with actin (one) (Fig. S16B and F), while TmP4 forms four ionic bonds with actin (Fig. S16C and F). Notably, the actin residues involved in interactions with Tm periods are not unique since E226, K238, D311, and K326 interact with multiple Tm periods, consistent with the amino-acid repetitiveness in the Tm periods.

Because of the periodic structure of Tm discussed above, the interaction of myosin loop-4 with Tm is unique for each actin along the length of the Tm molecule; therefore, it was completely obscured in previous structures due to helical averaging^12,29,48^. Figure 4 shows the unique interactions of myosin loop-4 with six of the seven actins that make up TF RU (one actin interacts with Tn core and is unavailable for actomyosin interactions). Since Classes 1-2 and 2-3, as well as Classes 5-6 and 6-7, had myosin heads attached to the same actin protomers, we confirmed that their loop-4 regions possessed the same interactions between loop-4, Tm, and actin (Fig. S17A and B). Therefore, we omitted Class 2-3 trailing and Class 5-6 trailing data in Figure 4. In all six compared myosin heads, loop-4 was stapled to the body of the actin filament via a salt bridge between its E371 and R147 of actin (Fig. 4B-G, red straight arrows).

**Figure 4.**
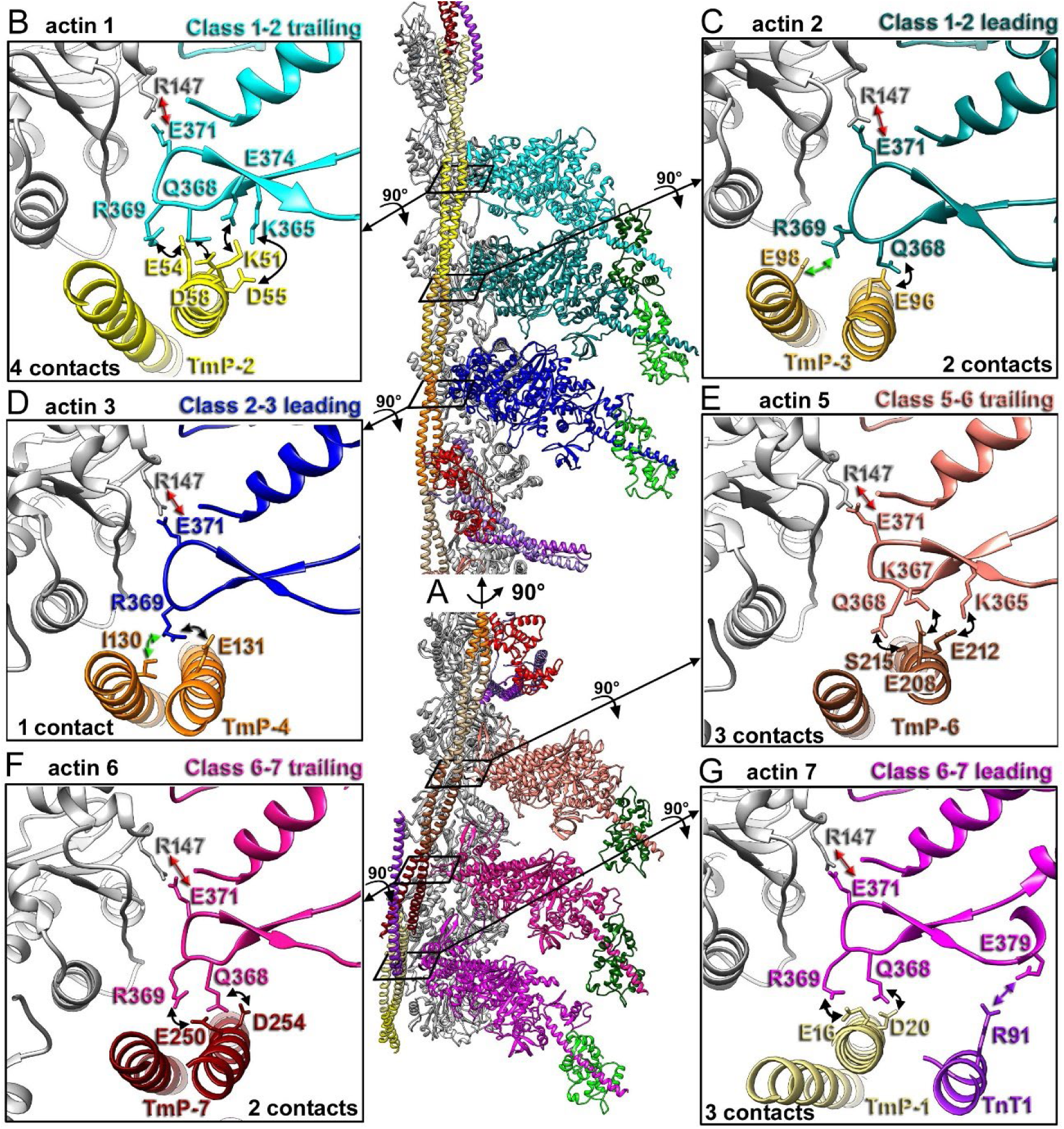
Interactions of myosin loop-4 with actin and Tm periods along RU. (A) Regions harboring loop-4 in the trailing or leading head are marked with black rectangles and detailed in B-G. (B-G) The color codes for the myosin heads and Tm periods are the same as in Fig. 1E-H. The mandatory interaction of E371 of loop-4 with R147 of actin is indicated with red straight arrows. Variable ionic interactions between loop-4 and the nearest Tm strand are marked with black curved arrows. Interactions of loop-4 with the distal Tm strand observed for Tm periods 3 (C) and 4 (D) are depicted with curved green arrows. Finally, the salt bridge between loop-4 and TnT1 is shown with a purple straight arrow in (G). The sequential numbers of actin protomers and the total number of contacts between loop-4 and Tm are indicated in black.

The positioning of Tm cable on the surface of actin near loop-4 (e.g., its activation state) was consistent across all six myosin heads (Figure S17C), but the interactions of loop-4 with Tm varied due to differences in the amino acid composition of Tm regions. The myosin head bound to actin 1 (Class 1-2 trailing head; Fig. 4B) had the highest number of interactions with Tm, forming four ionic bonds with Tm-P2 (Fig. 4B, black curved arrows). Heads attached to actins 2 and 3 interacting with Tm periods 3 and 4 were unique among the six because their loop-4 interacted with both Tm strands (Fig. 4C and D, green and black curved arrows), leading to an extended conformation of their loop-4 (Fig. S17C). Notably, these two heads formed fewer contacts with Tm than the other four heads – the head attached to actin 2 formed two ionic bonds (Fig. 4C), while the one attached to actin 3 formed one ionic and one hydrophobic bond via its R369, which we counted as one contact because the energy of a single hydrophobic bond is low (Fig. 4D). The head attached to actin 5 and 6 possessed three and two interactions with Tm, respectively (Fig. 4E and F, black curved arrows). Remarkably, the head bound to actin 7, in addition to its two interactions with Tm (Fig. 4G, black curved arrows), formed a salt bridge with R91 of TnT1 (Fig. 4G, purple straight arrow), highlighting the role of TnT in actomyosin interactions. Consistent with earlier predictions from the helical reconstructions of the cardiac actomyosin complex^12,29^, all six classes show that R369, located at the tip of loop-4, forms salt bridges with Tm residues. However, the other interactions reported here were previously unknown due to the helical averaging technique. Overall, the interactions of myosin heads with the Tm cable attached to actins 1, 5, and 7 were more prominent than those involving heads with actins 2 and 3 (Fig. B-G, contact numbers), which may have implications for the overall myosin interaction landscape on the surface of TF as suggested in the discussion. Not surprisingly, a large cohort of missense mutations in Tm is linked to various forms of human cardiomyopathy (summarized in Table S3). Notably, TnT R91, shown here to interact with loop-4 of myosin, is a hot spot for HCM mutations (human R92L/Q/W; see Table S3).

Our data indicate that the loop-4 contribution to myosin head interaction with Tm varies along the cTF RU, underscoring loop-4’s role in positioning myosin heads along the cTF RU.

### Interactions between myosin, ELC, and RLC

In line with recent cryo-ET data^26^, we demonstrate that a two-headed cardiac myosin molecule can have both heads attached to the TF in a strongly bound state (Fig. 1). This requires the two lever arms to adopt quite different conformations (Fig. 1N, 5A and B, and Movie S1). Comparison of the cardiac motor domain structures in the pre-power stroke, ADP, and rigor states (Fig. 5A) unambiguously shows that despite a dramatic swing in the rod domain between the trailing and leading heads, both myosin heads are in the rigor state – i.e., the swing of the lever arm occurs in the plane (Fig. 5A, red arrow) that is perpendicular to the plane of the power stroke lever arm movement (Fig. 5A, black arrows and transparent gray surface). Because the myosin molecule is flexible, we could resolve the neck region of the HMM (where the two RLCs interact) at 6 Å resolution (Fig. S19); therefore, we filtered the entire map to 6 Å resolution, as shown in Figure 1N. This is a dramatic improvement from the previously reported 30 Å cryo-EM structure^30^ and 12 Å cryo-ET data^26^. At this resolution, the secondary structure is clearly defined, showing that the rod domain of myosin (Fig. 5B, blue bracket), which contains both light chains (Fig. 5B, light and dark green ribbons), maintains its α-helical structure up to the kink region (Fig. 5B, blue arrows). The rod domains of the trailing and leading heads begin to diverge at the converter domain (Fig. 5B, converter), with an angular divergence of approximately 25° (Fig. 5B, yellow straight arrow). The trailing head has its rod domain relatively straight. In contrast, the leading head has its rod domain tilted even further after the ELC (Fig. 5B, blue arrowhead), which increases the angular divergence between the two rod domains to approximately 45° (Fig. 5B, curved yellow arrow).

**Figure 5.**
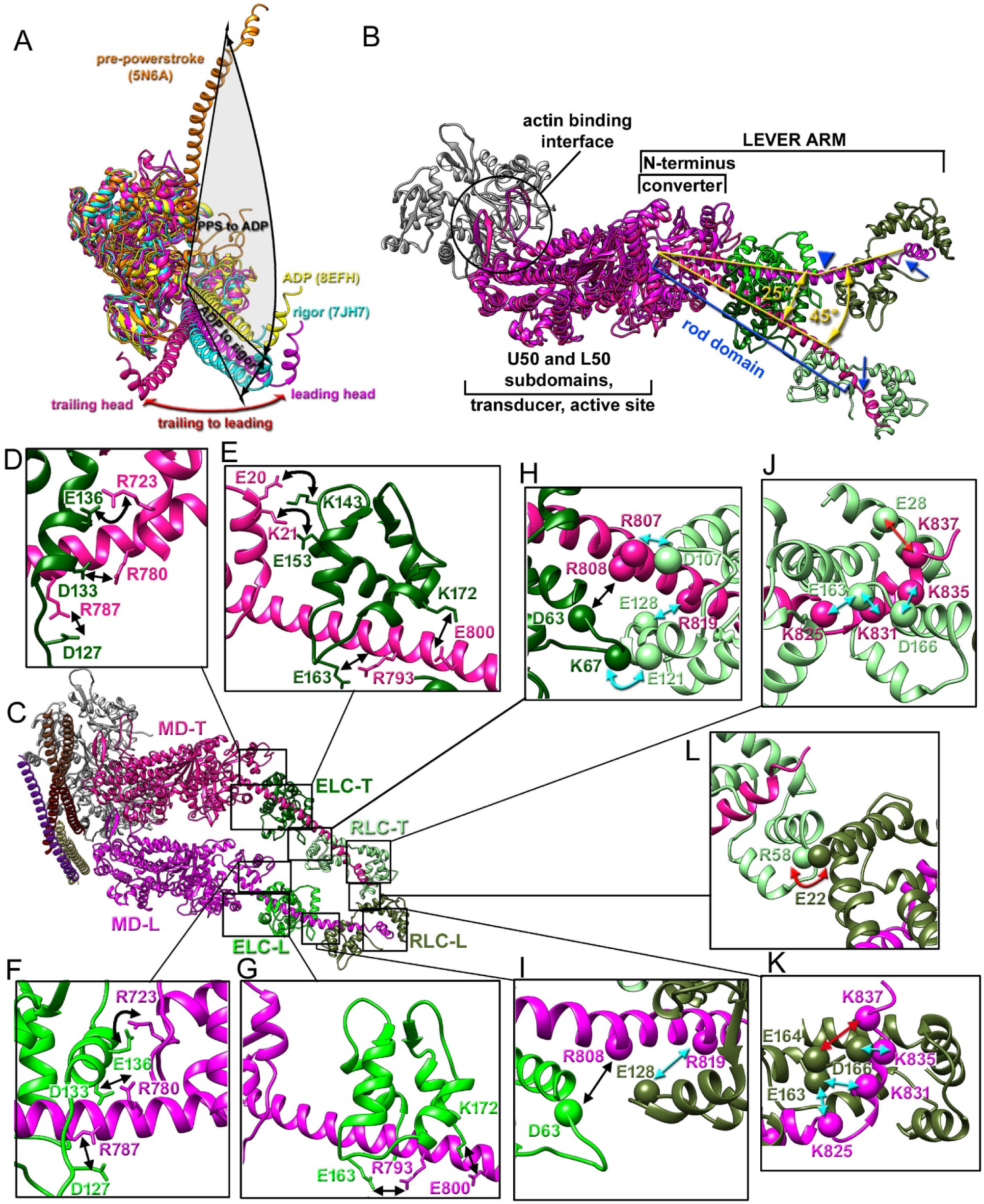
Myosin light chain interactions. (A) The structures of the pre-power stroke (PDB 5N6A), ADP (8EFH), and rigor (7JH7) of the myosin motor with modeled rod domains are shown in orange, yellow, and cyan, respectively, to illustrate the lever-arm swing during the power stroke. The plane of lever-arm movement is outlined in transparent gray. The trajectory of the rod domain upon transition from the trailing to the leading head is shown with a red arrow to demonstrate that it lies in the same plane as the rigor structure. (B) Alignment of trailing (deep pink) and leading (magenta) head motor domains from Classes 6-7 with respect to their positions on actin (gray) shows minimal divergence in the U50/L50 subdomains, 25° angular divergence in the region between the converter domain and the ELC-binding site on the rod domain (straight yellow arrow), followed by 45° angular divergence in the RLC-binding region (curved yellow arrow) due to a kink in the leading head’s rod domain (blue arrowhead). Hook regions of the myosin rod domain are marked with blue arrows. (C-K) Interactions between ELC and the myosin motor domain are marked with curved black arrows, while ELC interactions with the rod domain are marked with black straight arrows. A predicted interaction between trailing head ELC and RLC is depicted with a curved cyan arrow in (H). Predicted interactions between RLC and the myosin rod domain are marked with straight cyan arrows in (H-K). Unique interactions between the two RLCs and the myosin hook region are outlined with straight red arrows in (J) and (K). Finally, a predicted salt bridge between the two RLCs is denoted in (L) with a curved red arrow. Amino acids whose side-chain positions were derived from the 4.2 Å map are shown as atoms, while amino acids whose positions were predicted from the 6 Å map are denoted as spheres.

To build the model of the complete dual-headed myosin (Fig. 1N), we used the ELC structure from the high-resolution map (Fig. 1M), which yielded a perfect fit within the 6 Å map without any perturbations. The folding of the trailing (Fig. S20B) and leading (Fig. S20C) ELCs was consistent with the crystal structure of cardiac myosin^49^, with corresponding RMSDs of 1.65 Å^2^ and 1.35 Å^2^, respectively. Comparison of the RLCs with their counterparts in the cryo-EM structure of the cardiac thick filament^6^ yielded RMSDs of 4.3 Å^2^ (Fig. S20D) and 3.0 Å^2^ (Fig. S20E), which were expected given the different structural states of the extended and folded myosin.

The resolution at the interface between the myosin motor domains and adjacent ELCs, obtained for the map with RLCs masked (Fig. S8; 4.2 Å), enables us to directly compare this interface in the trailing and leading myosin heads (Fig. 5D, E, F, and G). Both ELCs form a salt bridge between E136 and R723 of the myosin motor domain (Fig. 5D and F, curved black arrows), a contact previously observed in the crystal structure of cardiac myosin^49^. However, the trailing ELC is positioned closer to the myosin motor domain than its leading counterpart. Hence, it forms two additional bonds with the myosin motor domain (Fig. 5E, black curved arrows), which are absent in the leading head (Fig. 5G). Our data also indicate that both ELC C-lobes form four salt bridges with their rod domains (Fig. 5D, E, F, and G, black straight arrows) that secure their binding to myosin. These include ELC residues D127, D133, E163, and K172 interacting with R787, R780, R793, and E800 of the rod domain, respectively. Due to resolution limitations at the edges of the map, we can only propose that D63 of the N-lobe of ELCs forms a salt bridge with R808 of the rod domain (Fig. 5H and I, straight black arrows), a finding previously reported in the cryo-EM structure of the folded back state of cardiac myosin^50^. Overall, our data suggest that the C-lobe mediates most of the ELC interactions with myosin via ionic bonds with both the motor and rod domains. It is worth noting that R723, R787, and R793 are linked to cardiomyopathies in humans (Table S3).

Finally, our 6 Å map of the neck region provides insights into RLC interactions with the ELCs, myosin rod domains, and each other (Fig. 5H-L), which are not equivalent between the trailing and leading heads due to differences in the geometries of their lever arms. Our data predict that E128, E163, and D166 of both RLCs form ionic bonds with R819, K825/K831, and K837 of their myosin rod domains (Fig. 5H-K, cyan straight arrows). In contrast, due to the proximity of the two light chains in the trailing head, ELC K67 may form a salt bridge with E121 of the trailing RLC (Fig. 5H, cyan curved arrow), which does not exist in the leading head (Fig. 5I). Also, RLC D107 likely interacts with R807 of the myosin rod domain only in the trailing head (Fig. 5H, cyan straight arrow). Due to differences in the positioning of the rod domain hook region, we predict that in the trailing head, RLC E28 makes an ionic interaction with myosin K837 (Fig. 5J, red straight arrow). In contrast, in the leading head, RLC E164 interacts with myosin K837 (Fig. 5K, red straight arrow). Finally, the proximity of the two RLCs in the myosin neck region points to a salt bridge between R58 and E22 (Fig. 5L, red curved arrow). Clinical data on mutations in myosin at R807, R819, K835, and K837, as well as missense mutations in ELC E153 and RLC E22, R58, E163, and D166 linked to human cardiomyopathies (see Table S3), verify these predicted interactions.

To summarize, the interactions between motor domains and ELCs differ between the leading and trailing heads when HMM is bound to cTF in the rigor state – the trailing ELC forms additional ionic interactions with the myosin motor domain, making its tail region structurally stiffer. Our data also indicate that, in addition to the rod domains’ interactions, the neck region is stabilized by an ionic bond between the RLCs.

### The structure of the Ca^2+^-bound Tn core in the fully activated myosin (M) state

Class 5-6 provided a unique opportunity to examine the structure of the Tn core of the Ca^2+^-bound TF in the myosin-bound state (M-state) at near-atomic resolution (Fig. S10B and C; ∼4.2 Å). Earlier, we reported two states of the Ca^2+^-bound Tn core – partially and fully activated^43^. Our data indicate that in the presence of Ca^2+^ and rigor myosin, the Tn core adopts only one conformation, a fully activated one.

In the previous structure of the Ca^2+^-bound TF^42,43^ the density of the 41 N-terminal residues of TnI was not observed. In contrast, in the presence of HMM, we observed an additional 10 amino acids extending to T32, which are proximal to the regulatory low-affinity Ca^2+^-binding site of TnC (Fig. 6B). Our data indicate that the three lysines (Fig. 6B; K37, K39, K41) in this region of TnI play an essential role in stabilizing the TnC structure. From NMR experiments^51^, TnC residues D65, D67, S69, T71, and E76 (Fig. 6B, green atoms) coordinate the Ca^2+^ ion (Fig. 6B, green sphere). Notably, TnI K37 and K39 form salt bridges with TnC D73 and E66 (Fig. 6B, cyan curved arrows) - residues in the Ca^2+^-binding loop of TnC that are not involved in coordinating the Ca^2+^ ion. Hence, K37 and K39 stabilize the low-affinity regulatory Ca^2+^-binding site of TnC. The charge-reducing mutation K37Q (human K36Q) is linked to DCM (Table S3), consistent with the proposed role for K37. TnI K41 stabilizes the positioning of the TnC N-helix by forming an ion bond with its E10 (Fig. 6B, straight cyan arrow). The N-helix is important for TnC Ca^2+^-sensitivity, since mutations in this helix are linked to cardiomyopathies^52^. Overall, our data provide a mechanistic role for the N-terminal region of TnI in TF regulation.

**Figure 6.**
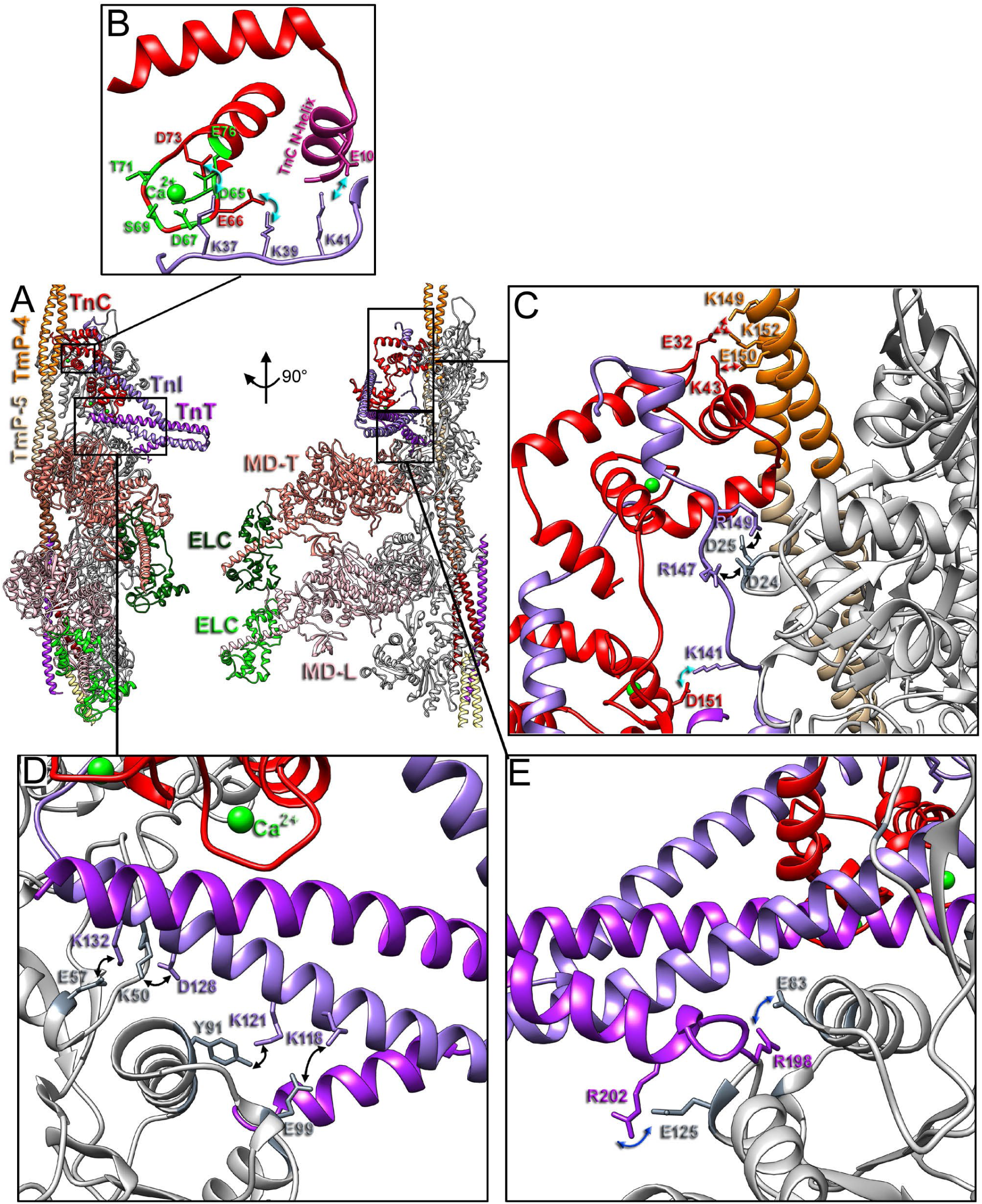
Interactions of the Tn core with Tm and actin. (A) Overview of Class 5-6, with regions of interest marked as black boxes. (B) Residues in the TnI N-terminal region (medium purple atoms) stabilize the TnC low-affinity regulatory Ca^2+^-binding site via ionic interactions (cyan arrows) with TnC residues (red atoms). The Ca^2+^ atom is shown as a green sphere, and the TnC residues that coordinate it are shown as green atoms. (C-E) TnC N-lobe interactions with Tm residues are marked with red arrows in (C). Ionic bonds between TnI and actin are marked with black curved arrows in (C-E), and the TnI salt bridge with TnC is depicted with a cyan arrow in (C). Ionic interactions between TnT and actin are marked with blue curved arrows in (E).

Our data indicate that TnC E32 forms ionic interactions with Tm K149 and K152 in Tm period 4, whereas TnC K43 forms a salt bridge with Tm E150 in the same period (Fig. 6C, red arrows). Hence, Tm stabilizes the open conformation of the TnC N-lobe. This is consistent with the activation of TF by cMyBP-C in the absence of Ca^2+ 43^.

The extended region of TnI, sandwiched between TnC and actin (porcine residues 137-151), plays an essential role in providing structural mobility to the Tn core^42,43^. Its upper part (R147 and R149) forms a prominent ionic interaction with the negatively charged actin loop (residues D24 and D25) (Fig. 6C, black curved arrows). At the same time, its K141, located below, forms a salt bridge with TnC D151 (Fig. 6C, cyan arrow). Therefore, this part of TnI serves as a “door hinge” that allows the whole Tn core to rock on the surface of actin. Actin D24N, K50T/Q, Y91C/H, E99K, along with TnI K147 (human K146S), are linked to cardiomyopathies in humans (Table S3).

The Tn core IT arm is formed by the two long and one short helices in TnT (Fig. 6D, purple) and TnI (Fig. 6D, medium purple) that provide structural support to secure the Tn complex on the surface of the actin backbone. Specifically, TnI K118, K121, D128, and K132 form ionic interactions with actin E99, Y91, K50, and E57, respectively (Fig. 6D, black curved arrows). A mutation in D128 (human D127Y) is associated with RCM/HCM (Table S3). The short TnT helix makes two salt bridges with actin E83 and E125 via its R198 and R202, respectively (Fig. 6E, blue curved arrows). Mutations in R202 (human R205L/W) are linked to DCM (Table S3).

To summarize, we show that the N-terminal residues of TnI stabilize the low-affinity regulatory Ca2+-binding site of TnC, whereas TnC stabilizes the fully activated state of Tm (M-state). In addition, TnI serves as a hinge between TnC and actin, providing rotational freedom to the Tn core while maintaining its attachment to actin and Tm.

## DISCUSSION

### The overall significance of the native cardiac cross-bridge structure

High-resolution structural studies of cardiac cross-bridges are essential for linking the molecular mechanisms of actomyosin interactions to whole-heart mechanics, since force and power output ultimately depend on how myosin heads are organized, recruited, and cycled between sequestered and activated states within cardiac sarcomeres. During the cardiac contraction–relaxation cycle, the two heads of β-cardiac myosin are mechanically and allosterically coupled. In relaxation, the heads adopt the IHM in which asymmetric head–head and head–tail interactions sequester myosin by sterically blocking key actin-binding elements ^6,21,50,53^. Upon muscle activation, disruption of the IHM increases the probability that one of the myosin heads will bind cTF and undergo a mechanochemical cycle, allowing inter-head strain to propagate through the shared myosin neck region (i.e., S2/lever-arm junction) so that the biochemical transitions of one head can bias those of its partner ^22,23,54,55^. In this context, the rigor cross-bridge structure with both myosin heads bound to cTF is particularly important because it provides a physiologically relevant^24^ “end-point” geometry for a true two-headed attachment that defines differential axial positions of the paired lever arms within the active cross-bridge and establishes essential boundary conditions for *in silico* modeling of cardiac force production and cooperative recruitment that cannot be obtained from single-headed (i.e., S1-decorated) actomyosin alone.

Using time-resolved cryo-EM, as done for the actomyosin-V S1 mutant with reduced ATP turnover^19^, to capture all possible structural states of the HMM bound to the native cTF in the presence of ATP would provide a complete mechanistic view of the cardiac actomyosin cycle. However, the rapid rate of ATP hydrolysis by the cardiac HMM, the large number of structural intermediates arising from the presence of two heads, and the heterogeneous positioning of the heads along the cTF make this task currently unachievable.

### The interactions of leading and trailing myosin heads with cTF RU are not identical

Our structural understanding of the interactions between the myosin motor domain and actin in the rigor state is based on helically averaged cryo-EM structures. While this approach is suitable for single-headed myosins (e.g., myosin-Ib^18^), its application to two-headed myosins (e.g., class II myosins^12,13,15^) results in a loss of important information about interactions between the two myosin heads (Fig. 3). Importantly, muscle myosins do not interact with bare F-actin, but rather with an actin-based multiprotein complex – the TF, which, because of a lack of helical symmetry, makes actomyosin interactions along the TF non-equivalent (Figs. 3 and 4). Here, we provide detailed information on how the two heads of cardiac myosin interact with one another and with the actin protomers that comprise the cTF regulatory unit.

The interactions of myosin with cTF are mediated by five myosin elements (Fig. 1B) – CM-loop, HLH-motif, loop-2, loop-3, and loop-4. We show here that the interactions of the CM-loop or HLH-motif with actin are identical for the two heads and redundant along the cTF RU (Fig. 2). The HLH-motif is the most conserved part of the actomyosin interface involved in the initial weak binding of myosin^19^. Biochemical mutagenesis studies have confirmed its importance for myosin’s ATPase activity and motility^56^. The CM loop is crucial for stabilizing the strongly bound actomyosin upon Pi release and closure of the myosin cleft^19^. Not surprisingly, class II myosins exhibit very similar interfaces between their CM loops and actin^12,13,15^. Along with the HLH-motif and CM-loop, the loop-4 interaction with actin is conserved across all RU actins involved in actomyosin interactions (Fig. 4, straight red arrows). In a diverse array of previously published rigor actomyosin structures^12,13,15,17^, an acidic residue in loop-4 forms a charged interaction with the same basic residue on actin. Overall, our data for the CM-loop, HLH-motif, and loop-4 interactions with actin are consistent with previously published helically averaged structures, as these myosin elements show uniform interactions across the RU.

In contrast to its binding to actin, loop-4 interactions with Tm differ because of Tm sequence variability along the RU (Fig. 4), resulting in fewer interactions of myosin heads with Tm periods 3, 4, and 7 (Fig. 4C, D, and F), while providing more extensive interactions of myosin heads with Tm periods 1, 2, and 6 (Fig. 4B, E, and G). Importantly, we show that myosin heads interact with TnT1 via loop-4 (Fig. 4G, purple straight arrow), thereby demonstrating a direct role of TnT in regulating actomyosin interactions. This has implications for myosin targeting on the cTF RU discissed further below.

Loop 2 exhibits substantial variations in length and flexibility among myosin isoforms, consistent with its importance^56,57^ in tuning actomyosin interactions for different cellular needs^58^. The degree of visualization varied across helical reconstructions of different myosins^12-18^, but its density has never been fully resolved. Regrettably, Loop-2 density was largely absent in the helically-based reconstructions of cardiac actomyosin complexes ^12,13,29^. The single-particle approach enabled us to resolve most of loop-2 and show that some of its interactions are uniform across the RU (Fig. 3, straight and curved black arrows), whereas others form only when loop-2 resides in the leading head (Fig. 3, red arrows) or is proximal to the Tn core (Fig. 3, green arrows). Of note, we observe a cohort of ionic interactions between loop-2 and actin across all classes, involving both leading and trailing heads (Fig. 3, straight and curved black arrows). However, the positioning of the N-terminus of actin and the trajectory of loop-2 are not identical across the classes, demonstrating flexibility in these two mobile regions. The uniform interactions include ionic bonds between K639/K640, located at the base of loop-2, with myosin E535 and D2 of the N-terminus of actin (Fig. 3, black straight arrows), along with salt bridges between loop-2 E632/K635 and R95/E100 of actin (Fig. 3, black curved arrows). Notably, when loop-2 is pushed down by the bottom part of the Tn core (Fig. 3E, purple), the aforementioned interactions are disrupted, and K635 forms an H-bond with TnT N266, while E632 forms a salt bridge with K405 of the CM-loop (Fig. 3E, green curved arrows), which presumably reduces the affinity of the myosin head for actin below the Tn core (discussed below).

The communication between the two myosin heads has been recognized using different techniques ^22,23,54,55^. Here, we show that, in addition to the self-evident interaction between the heads via the myosin neck region (e.g., RLCs and S2), the two heads communicate via the interaction between the leading head’s loop-2 K635 and the trailing head’s loop-3 E574 above it (Fig. 3, red arrows). This interaction between loops 2 and 3 creates an additional ionic bond, thereby stabilizing the actomyosin complex. Given that loop-2 is involved in both weak and strong actomyosin interactions^19,56^, its contact with loop-3 of the trailing head may either promote trailing head binding to the cTF or, in the case of a weakly bound trailing head, enhance strong binding of the leading head. In either case, our data are consistent with the early proposed idea^22^ that “even if only one head actually performs the work, the other serves to guide the operative head to its maximally effective orientation”. Direct communication between the two myosin heads via interaction of loops 2 and 3, reported here, reveals a novel mechanism of allosteric coupling between the two heads of cardiac myosin. The summary of interactions of the two myosin heads with actin, Tm, and each other is shown in Fig 7A.

**Figure 7.**
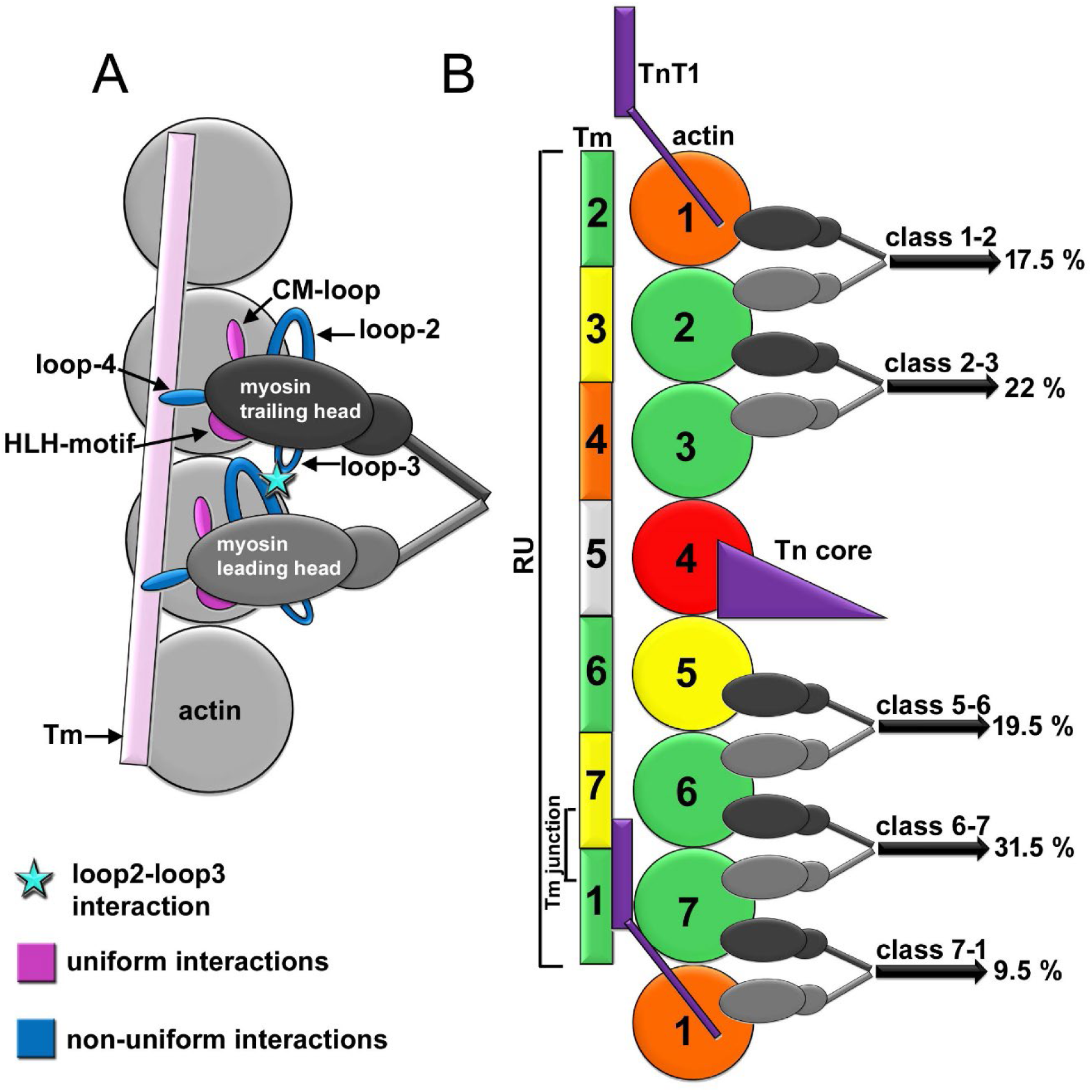
Summary of cardiac myosin interactions with cTF. (A) The two myosin heads have uniform interactions between their CM-loop and HLH-motif (marked in magenta) with actin, whereas loop-2, loop-3, and loop-4 (marked in blue) of the trailing (dark gray) and leading (light gray) heads interact differently with actin/Tm. Loop-2 of the leading head interacts with loop-3 of the trailing head (cyan star). (B) Actin molecules that comprise cTF RU are shown as circles, Tm periods bound to these actins are shown as rectangles, the actin-4 blocking Tn core is depicted as a purple triangle, and TnT1 is shown as a purple rectangle with its N-terminus as a purple line. Binding of the double-headed cardiac myosin to cTF RU depends on the availability of actin (restricted by the Tn core, actin is red; partially restricted by the TnT1 N-terminus, actin is orange; somewhat restricted by Tn core, actin is yellow; fully available, actin is green) and the strength of interactions with Tm (green ≥3 contacts, yellow=2 contacts, orange=1 contact, gray=no interactions). Interaction of the leading head (light gray) with partially restricted actin (e.g., Class 7-1) is less favorable than if the trailing head binds to partially restricted actin (e.g., Class 1-2). The measured frequency for each position is shown as a percentage, and the actin protomer and Tm period numbers within the cTF RU are shown in black.

Of note, we confirm our previous observation^12^ that, in contrast to other myosin II isoforms ^15,16^, loop-3 of cardiac myosin does not form a “Milligan contact” with actin.

### Myosin loops 2, 3, and 4, along with TnT, guide HMM positioning on the surface of the cTF RU

Cryo-ET of the murine skeletal sarcomere in the rigor state^25^ showed that myosin binding is a stochastic process regulated by physical constraints, such as actin subunit orientation, resulting in a pseudo-regular arrangement of cross-bridges spaced by 37 nm (i.e., the actin crossover length). However, due to the high cross-bridge frequency, cryo-ET failed to visualize Tn complexes, leaving the positioning of myosin on the TF RU undefined. Here, we show that, except for actin 4, which holds the Tn core and is marked in red in Fig. 7B, almost all actins within the RU can interact with myosin. Their preference for myosin binding depends on the spatial constraints imposed by TnT and the specific Tm repeat, as well as on cross-talk between the two heads arising from the interaction between loops 2 and 3, as discussed above. These factors are outlined in Fig. 7B along with the experimentally measured frequencies of HMM binding to adjacent actin protomers within the RU. The lowest frequency (9.5%) of Class 7-1 may be explained by a clash between the TnT1 and the myosin head (Fig. S6C, large black arrow). Hence, actin 1 is marked in orange in Fig. 7B. However, a much higher frequency of Class 1-2 (17.5%) suggests that the type of the head that interacts with TnT1-constrained actin matters – the leading head interaction with actin 1 is more prohibitive than that of the trailing head. We assume that the stabilization of the trailing head via the loop 2 and 3 interaction alleviates the TnT1 inhibitory effect. A slight clash between myosin and the TnT C-terminus (Fig. S6E, small black arrow), along with decreased interactions of loop-2 with actin caused by its being pushed down by the Tn core’s bottom (Fig. 3E), renders actin 5 less suitable for myosin attachment (Fig. 7B, shown in yellow), consistent with the 19.5% frequency of Class 5-6. The interaction of the leading head of Class 5-6 with the unrestricted actin 6 presumably alleviates the negative effects of Tn on the trailing head’s binding to actin 5 via the loop-2 and loop-3 interaction. The interaction of HMM with actins 2-3 and 6-7 is unrestricted by the Tn subunits, but Class 6-7 still has a measurably higher frequency (31.5%) than Class 2-3 (22%). This coincides with more extensive interactions of both loop-4s in Class 6-7 with Tm repeats 7 and 1 (Fig. 4F and G, 5 contacts in total) compared with the fewer contacts of Class 2-3 loop-4s with Tm repeats 3 and 4 (Fig. 4C and D, 3 contacts in total). Notably, we found that Tm repeat 2 forms the most extensive interactions with loop-4 of myosin attached to actin 1 (Fig. 4B, 4 contacts), which presumably reduces the constraining effect of TnT1 on myosin binding to actin 1. In fact, our data suggest that the loop-4 interaction with Tm is an important adaptation that mitigates the effects of the Tn subunits on actomyosin interactions within the cTF RU (hence, very comparable frequencies), which is in general agreement with the overall stochastic distribution of cross-bridges along TFs observed by cryo-ET^25^. On the other hand, our data indicate that actins 6 and 7, located at the Tm junction region, comprise a somewhat more favorable site for myosin binding (spaced by 37 nm), which may have a direct physiological consequence – the univocal distance from the junction region to the two Tn cores allows myosin to cooperatively activate two RUs simultaneously. To summarize, our data indicate that a complete description of actomyosin interactions must account for communication between the two myosin heads via loops 2 and 3, variable interactions of loop 4 with Tm repeats, and the effects of TnT.

### TnI stabilizes Ca^2+^-bound TnC and enables rotational flexibility of the Tn core

Because of the inherent flexibility of the cTF elements^43^, only the junction region of the cTF had previously been resolved at near-atomic resolution^39^. Now, with HMM and similar to what was found in the presence of cMyBP-C^43^, the cTF Tn core was effectively locked in a single fully activated structural state, greatly reducing cTF’s intrinsic structural heterogeneity and enabling the acquisition of near-atomic details of the Tn core’s interactions with actin and Tm, which were previously missing.

First, we demonstrate that extensive interactions between the TnC N-lobe and the third period of Tm (Fig. 6C, red arrows) stabilize Tm in the M-state, uncovering a new role for the Ca^2+^-binding TnC in regulating cTF. Second, we reveal novel, important roles for TnI. The well-known function of TnI is to stabilize the inhibited state of TF by interacting with the actin/Tm via its C-terminus^40-42^. Here, we show that the three lysines at the N-terminus of TnI stabilize the regulatory low-affinity Ca^2+^-binding site of TnC (Fig. 6B, cyan arrows) when TF is in the M-state. This provides a mechanistic explanation for the increased affinity of cTFs for Ca^2+^ in the presence of active cross-bridges^59,60^, since such an interaction was not previously detected in the Ca^2+^-free or Ca^2+^-bound cTF in the absence of myosin^40,41,43^. Furthermore, we show that TnI plays a crucial role in positioning the inherently mobile^40,41,43^ Tn core region on the actin surface, functioning as a hinge between the actin surface and the TnC subunit (Fig. 6C and D). Notably, the helical region of TnT adjacent to the intrinsically disordered TnT-linker provides an additional anchor point for the Tn core on actin’s surface (Fig. 6E) and thus plays a role in the Tn core’s swing upon cTF activation^40,41,43^.

### The trailing and leading head lever arms differ structurally

Recent cryo-EM of the double-headed actomyosin complex in the ADP state^30^ showed that the leading lever arm was more biased toward the pre-power-stroke state, whereas the trailing lever arm was pulled toward the rigor state. Consistent with these findings, comparing our rigor structure with the ADP structure reveals that in both, the trailing lever arm is in the rigor position (Fig. S20F, blue arrow). However, our leading lever arm (rigor) differs from the ADP structure, where the lever arm is racked up (Fig. S20F, red arrow), as expected given the bound ADP.

Cardiac myosin light chains provide a key regulatory layer that tunes both cross-bridge mechanics and thick-filament activation in the heart. The ELC, through its cardiac N-terminal extension (NTE), interacts with actin and regulates the actomyosin cycle^61-63^. Of note, we have not observed any defined additional density that could be attributed to the NTE helix predicted to interact with actin, suggesting that NTE interaction with cTF is stochastic (i.e., involves multiple sites with comparable affinities). ELC plays a role in maintaining compliance at the head–neck junction^49,64^. Notably, we found that the interactions between the ELC and the myosin head differ between the leading and trailing heads – the trailing head has additional salt bridges (Fig. 5E, curved black arrows) that are absent in the leading head (Fig. 5G), indicating that the trailing lever arm is more mechanically rigid. Complementarily, experiments involving cardiac RLC removal^65^ demonstrated that disrupting RLC support of the myosin neck region alters lever-arm mechanical behavior, suggesting a role in providing lever-arm rigidity. Again, our data indicate that the trailing head has an ion bond between the ELC and RLC (Fig. 5H, cyan curved arrow) that is absent in the leading lever arm (Fig. 5I), suggesting enhanced mechanical compliance of the trailing lever arm relative to the leading lever arm.

Because key transitions in the actomyosin mechanochemical cycle, especially ADP release, are strain-sensitive^66,67^, the difference in mechanical compliance between the two lever arms when both are bound may result in different kinetics for the two heads. One would expect that the enhanced rigidity of the trailing lever arm relative to the leading lever arm should redistribute strain and thereby alter contractility in two possible ways: (1) a stiffer trailing lever arm should increase instantaneous cross-bridge stiffness, potentially raising force per attached head; (2) greater rigidity would also be expected to increase the effective trailing lever arm strain during the two-head–bound intermediate that may speed trailing-head ADP release and detachment, potentially increasing shortening velocity and power. These predictions are consistent with the requirements of both heads for maximal force output^22^.

## Materials and Methods

### Proteins and Buffers

Porcine native cardiac TFs were prepared as described in^41^. HMM purification from Triton-washed myofibrils was done using the previously published protocol^68^ modified as described in^69,70^. A-buffer was used for cryo-EM experiments: 50 mM potassium acetate, 10 mM 3-(N-morpholino) propane sulfonic acid (MOPS), 3 mM MgCl_2,_ and 0.1 mM CaCl_2_ (pCa=4). The pH was adjusted to 7.2 at 7°C after all components were mixed.

### Cryo EM

A total of 1.5 μL of 1 μM cTFs in A-buffer was applied to the center of the glow-discharged lacey carbon grid at 7°C in the Vitrobot Mark IV (FEI, Inc) chamber, mixed with 1.5 μL of 0.1 μM HMM in A-buffer on the surface of the grid, and immediately blotted with Whatman #1 filter paper for 3-5 sec and vitrified. Summaries of imaging conditions and image reconstruction are provided in SI Appendix Table S1. Image analysis was performed using RELION^46^, while modeling was done using UCSF Chimera^71^, Namdinator^72^, ISOLDE^73^, and PHENIX^74^. Validation statistics for models are provided in SI Appendix Table S2. Experimental details are provided in SI Materials and Methods.

## Acknowledgments

This work was supported by NIH grant R01 HL160966 (to V.E.G., J.R.P and P.B.C), and R01GM120137 (to V.E.G.), J.R.P. provides consulting to Kate Therapeutics.

## Author contributions

C.R. and T.N. performed image analysis and 3D-reconstructions;

C.R. did flexible fitting and model refinement; B.B. purified proteins in H.D.W.’s lab; H.D.W., P.B.C., and J.R.P. read the manuscript; C.R. and V.E.G wrote the SI; V.E.G. conducted cryo-EM, performed image analysis, 3D-reconstruction, model refinement, and wrote the manuscript.

## Data availability

Accession numbers:

PDB: 9YA8, 9YAQ, 9YJP, 9YK9, 9YKN

EMD: 72724, 72734, 73030, 73042, 73055

Details provided in Tables S1 and S2.

## Declaration of Interests

The authors declare no competing interests.

## Notes

### Competing Interest Statement

The authors have declared no competing interest.

