## Supplementary material for "The structure of the native cardiac crossbridge in the rigor state": Supplimental Information

**This PDF file includes:**

Supplementary text
Figures S1 to S20
Movie S1
Tables S1 to S3
SI References

**Supplementary Information Text**

**SI Materials and Methods.**

**Imaging and image analysis.** Details of imaging conditions and image reconstruction are provided in Table S1. Micrographs were collected on a 300-kV Titan Krios electron microscope equipped with a K3 direct electron detector and an energy filter operating in super-resolution mode. Micrographs were recorded in 40 subframes at a dose rate of ~0.85 e-/Å2 per frame over a defocus range of 0.5–3.5 µm, with a pixel size of 0.678 Å. All processing was performed in RELION [1]. Images were imported into RELION for motion correction and dose weighting using internal RELION algorithms. The images were binned to a pixel size of 1.356 Å, and contrast transfer function (CTF) parameters were determined using CTFFIND4 [2]. Particle selection was performed manually from 41,405 micrographs to extract 5,235,506 cTF segments (537X537 Å with 27.4 Å overlap) having HMM sparsely bound (Figure S4A, red arrows). Particles were decimated to 4.068 Å/px (Figure S4B, S5A-B). To evaluate HMM interactions along the length of the entire regulatory unit (RU), which comprises seven actin molecules, we generated low-resolution models of cTF with an attached two myosin heads (Figure 5C, short surfaces) to cover all possible positions of HMM within the TF RU. 40 classes based on the relative position of the Tn complex (Figure S5C, red arrows) and HMM (Figure S5C, green arrows) were generated. The resulting class averages differed from the initial models due to the presence of RLCs in the myosin neck region (Figure S5C, long surfaces). Because cTF is comprised of two strands, some classes were redundant in terms of HMM binding to cTF RU. We used segments from the following five classes (Figure S5C): (1) Class39 from the 40-class sorting, representing HMM bound to actins 6 and 7 of the RU (referred to as Class 6-7); (2) Class31 with HMM bound to actins 5 and 6 (referred to as Class 5-6); (3) Class11 with HMM bound to actins 2 and 3 (referred to as Class 2-3); (4) Class10 with HMM bound to actins 1 and 2 (referred to as Class 2-3); (5) Class37 representing HMM bound to actins 7 and 1 (referred to as Class 2-3) for 3D focused refinement [3-5]. Details for the refinement of each class are provided below.

Class 6-7: The overview of the Class 6-7 reconstruction is shown in Figure S7. The class output from the initial position sorting (Figure S7A, hot pink surface), in addition to the expected position of HMM relative to the Tn core (Figure S7A, blue arrow), contained traces of additional HMM molecules bound elsewhere (Figure S7A, magenta and brown arrows), suggesting heterogeneity within the class. To overcome this, three individual templates were built using the initial class average (Figure S7A, hot-pink surface) as a guide. Templates 1 (Figure S7B, orange surface) and 2 (Figure S7B, purple surface) contained additional HMM molecules above or below the expected position (Figure S7B, brown and blue arrows or magenta and blue arrows, respectively), while the third template had only one HMM molecule (Figure S7B, green surface and blue arrow). Templates were low-pass filtered to 15 Å and used for supervised 3D classification in RELION [1] (Figure S7B, solid surfaces). As expected, double-decorated classes represented a small fraction of the class: double HMM occupancy mode 1 (n=11,675) (Figure S7C, orange surface), double HMM occupancy mode 2 (n=10,791) (Figure S7C, purple surface), and single HMM occupancy (n=86,644) (Figure S7C, green surface). The class representing the single HMM occupancy (n=86,644) (Figure S7C, red box) was re-extracted at the full scale of 1.356 Å/px, 3D-refined starting from the class average low-pass filtered to 15 Å (Figure S7C, green surface), and post-processed (Figure S7D). To obtain a high-resolution map, the particle polishing algorithm in RELION was used for reference-based beam-induced motion correction. The regulatory light chains (RLCs) were not well resolved in the initial refinement (Figure S7D, black arrow), suggesting that, due to disorder, RLCs required further classification (discussed in detail later). Finally, to obtain a high-resolution structure of the HMM bound to actin molecules 6 and 7, a focused refinement was done using a 10 Å mask that excluded the opposite TF strand, distal actin molecules, and RLCs (Figure S7E, red surface). The resulting high-resolution density map was segmented (Figure S7F, yellow surface) and used to model (Figure S7G, cyan ribbons).

The global resolution for the map (4.3 Å) (Figure S8A) was calculated in RELION using the gold-standard FSC criterion of 0.143, which measures the normalized cross-correlation between the two half-maps in Fourier space. To do this, the entire particle subset is randomly split into two halves, and each half is independently reconstructed into a density map. This method of resolution determination is problematic when a cryo-EM density map exhibits resolution variability across local regions due to heterogeneity or intrinsic disorder, and it also yields the most pessimistic resolution estimate by reducing the signal-to-noise ratio (SNR) in the two halves [6, 7]. We used RELION to calculate a local resolution map, which showed a range of 3.8-9.2 Å (Figure S8B). The region resolved at the highest resolution of 3.8 Å included actin and the actomyosin interface (Figure S8B, black arrows), whereas the ELC interface with the myosin motor domain was resolved at 4.2 Å (Figure S8B, black arrow). As expected, the distal portions of the map were the lowest resolution (Figure S8B, red arrows). Given the drawbacks of the RELION resolution-determination algorithm discussed above, we also used CryoRes, a deep-learning-based algorithm [6], to determine local resolution from the cryo-EM density map (Figure S8C). The resolution estimate ranged from 3.79 Å to 4.27 Å within the map (Figure S8C), consistent with the RELION local resolution determination. That resolution range was supported by the map's quality, enabling visualization of side chains in the cryo-EM map (Figure S8D). Detailed views of the map components (actin, myosin motor domain, myosin CM-loop, loop-2, loop-3, loop-4, HLH-motif, and the ELC interface) at 3.8-4.2 Å resolution that are discussed in the paper are shown along with the atomic model to illustrate the quality of the data (Figure S8D).

<u>Class 5-6</u>: The overview of the Class 5-6 reconstruction is shown in Figure S9. The class output from the initial position sorting (Figure S9A, hot pink surface) contained traces of HMM near the lower region of the filament (Figure S9A, purple arrow). Two templates were generated using the initial class average (Figure S9A, hot-pink surface) as a reference. Template 1 (Figure S9B, yellow surface) contained an HMM molecule at its expected position (Figure S9B, orange arrow) while the second template (Figure S9B, cyan surface) contained an additional HMM molecule where the traces were observed (Figure S9B, purple and orange arrows). The templates were low-pass filtered to 15 Å and used for supervised 3D classification in RELION (Figure S9B, solid surfaces). Consistent with the initial class average, the majority of segments were classified as having one HMM molecule bound to actins 5 and 6 (n=55,229) (Figure S9C, yellow surface), with a few segments possessing two HMM molecules bound (n=5,451) (Figure S9C, cyan surface). The primary class (Figure S9C, red box) was re-extracted at 1.356 Å/px and reconstructed (Figure S9D) starting from the class average (Figure S9C, yellow surface), low-pass filtered to 15 Å. To obtain a high-resolution map, the particle polishing

algorithm was applied before focused refinement. To get the high-resolution structure, we used a 10 Å mask that excluded the opposing TF stand and distal actin molecules, as well as the RLCs (Figure S9E, green surface). The high-resolution density map was segmented (Figure S9F, lavender surface) and used for modeling (Figure S9G, blue ribbons).

The global (4.6 Å) (Figure S10A) and local resolutions (4.0 – 8.6 Å) (Figure S10B) were calculated using RELION's built-in algorithms. A local resolution map generated using RELION yielded the highest resolution of 4.0 Å (Figure S10B, black arrows) within actin, the myosin motor domain, and at the actomyosin interface. The resolution estimation at the ELC interface was 4.4 Å (Figure S10B, black arrow). The resolution estimation determined by CryoRes [6] ranged from 3.85 Å to 4.3 Å (Figure S10C), which was in line with the RELION local resolution map. Resolution estimates were consistent with the details in the cryo-EM map (Figure S10D). Detailed views of the components of the map, along with associated atomic models discussed in the manuscript, are depicted in Figure S10D.

<u>Class 2-3</u>: Details of the processing are shown in Figure S11. The class average from the initial position sorting (Figure S11A, hot pink surface) contained traces of additional HMM molecules bound to actins rather than 2 and 3 (Figure S11A, dark red, chartreuse, purple, brown, teal, and orange arrows). To sort out those minor modes, four templates were made using the initial class average (Figure S11, hot pink surface) as a reference. Template 1 (Figure S11B, yellow surface) contained one HMM molecule bound to actins 2 and 3 (Figure S11B, orange arrow), template 2 contained three HMM molecules (Figure S11B, green surface, orange, brown, and teal arrows), template 3 (Figure S11B, lavender

surface) also contained three HMM molecules (Figure S11B, orange, dark red, and chartreuse arrows), and template 4 (Figure S11B, cyan surface) contained two HMM molecules (Figure S11B, orange and purple arrows). The templates were low-pass filtered to 15 Å and used for supervised 3D classification in RELION (Figure S11B, solid surfaces). The frequencies of resultant modes are as follows: single HMM occupancy (n=63,599) (Figure S11C, yellow surface), triple HMM occupancy mode 1 (n=2,783) (Figure S11C, green surface), triple HMM occupancy mode 2 (n=2,916) (Figure S11C, lavender surface), and double HMM occupancy (n=10,691) (Figure S11C, cyan surface). The main class with HMM bound to actins 2 and 3 (n=63,599) (Figure S11C, yellow surface and brown arrow) had weaker-than-expected density for the Tn core (Figure S11C, yellow surface and grey arrow), suggesting that HMM bound next to the Tn core might cause partial dissociation of the Tn complex. To obtain the structure of the intact cross-bridge, segments selected in (C) were sorted by the integrity of the Tn tandem (Figure S11D, solid surfaces). The frequencies of resultant modes are as follows: intact Tn tandem (n=31,180) (Figure S11E, magenta surface) and upper Tn occupancy (n=32,419) (Figure S11E, blue surface). The class with the intact Tn tandem (Figure S11E, magenta surface, red box) was re-extracted, refined, and post-processed at 1.356 Å/px (Figure S11F), starting from the class average, low-pass filtered to 15 Å (Figure S11E, magenta surface). To obtain a high-resolution map, the particle polishing algorithm was applied before focused refinement. Next, we used a focused refinement approach (Figure S11) by utilizing a 10 Å mask that excluded the opposite strand of TF, along with distal actins and RLCs, from the refinement process (Figure S11E, red surface). The high-resolution density map was segmented (Figure S11F, green surface) and used for modeling (Figure S11I, purple ribbons).

The global (4.8 Å) (Figure S12A) and local resolutions (4.2 – 8.7 Å) (Figure S12B) were calculated using RELION. The local resolution map yielded the highest resolution of 4.2 Å (Figure S12B, black arrows) within actin, myosin motor domain, and at the actomyosin interface (Figure S12B). CryoRes [6] resolution estimate ranged from 4.27 to 4.68 Å (Figure S12C). This local resolution determination was consistent with visualization of large amino acid side chains in the cryo-EM map (Figure S12D).

<u>Class 1-2</u>: Details are shown in Figure S13. The class average from the initial position sorting (Figure S13A, hot pink surface) contained traces of extra density at the top and bottom of the map (Figure S13A, light grey, cornflower, light green, gold, dark red, dark grey, teal, light purple, sienna, deep pink, and khaki arrows), suggesting heterogeneity. Twelve templates were made based on the initial class average (Figure S13A, hot pink surface). The first template (Figure S13B, cerulean surface) contained an HMM molecule bound to actins 1 and 2 (Figure S13B, sienna arrow). In addition we generated ten templates possessing two or three HMM molecules bound to the TF: template 1 (Figure S13B, chartreuse surface, sienna and dark red arrows), 2 (Figure S13B, lavender surface, sienna and dark grey arrows), 3 (Figure S13B, magenta surface, sienna and khaki arrows), 4 (Figure S13B, orange surface, sienna and deep pink arrows), 5 (Figure S13, green surface, sienna and teal arrows), 6 (Figure S13B, red surface, sienna and light green arrows), 7 (Figure S13B, purple surface, sienna and cornflower arrows), 8 (Figure S13B, yellow surface, sienna and light grey arrows), 9 (Figure S13B, blue surface, sienna and light purple arrows), and template 10 (Figure S13B, dark green surface, sienna and

gold arrows). There was one triple HMM occupancy mode (Figure S13B, cyan surface, teal, sienna, and dark green arrows). The templates were low-pass filtered to 15 Å and used for supervised 3D sorting in RELION (Figure S13B, solid surfaces). The frequencies of resultant modes were as follows: single HMM occupancy (n=54,451) (Figure S13C, cerulean surface), double HMM occupancy mode 1 (n=508) (Figure S13C, chartreuse surface), double HMM occupancy mode 2 (n=267) (Figure S13C, lavender surface), double HMM occupancy mode 3 (n=438) (Figure S13C, magenta surface), double HMM occupancy mode 4 (n=289) (Figure S13C, orange surface), double HMM occupancy mode 5 (n=4,391) (Figure S13C, green surface), double HMM occupancy mode 6 (n=5,249) (Figure S13C, red surface), double HMM occupancy mode 7 (n=1,670) (Figure S13C, purple surface), double HMM occupancy mode 8 (n=151) (Figure S13C, yellow surface), double HMM occupancy mode 9 (n=5,447) (Figure S13C, blue surface), double HMM occupancy mode 10 (n=2,210) (Figure S13C, dark green surface), and triple HMM occupancy (n=120) (Figure S13C, cyan surface). The primary class representing one HMM molecule bound to actins 1 and 2 (n=54,451) (Figure S13C, cerulean surface, red box) was re-extracted, refined, and post-processed at a raster of 1.356 Å/px (Figure S13D) starting from the class average filtered to 15 Å (Figure S13C, cerulean surface). To achieve a high-resolution map, the particle polishing algorithm was used for reference-based beam-induced motion correction (Figure S13D). Next, we used focused refinement to obtain the high-resolution 3D reconstruction (Figure S13E-G). This was done by utilizing a 10 Å mask that excluded the TF opposite strand, distal actins, and RLCs (Figure S13E, light yellow surface). The resultant density map was segmented (Figure S13F, light pink surface) and used for modeling (Figure S13G, dark brown ribbons).

The global (4.5 Å) (Figure S14A) and local resolutions (3.9 – 8.5 Å) (Figure S14B) were calculated using RELION built in algorithms. The local resolution map calculated by RELION was consistent with resolution estimation from CryoRes [6] and ranged from 3.78 to 4.2 Å within actin, myosin motor domain, and ELCs interface with myosin (Figure S14B and C). Close-up views of the actual map along with the atomic model in the regions of interest discussed in the manuscript are shown to illustrate the quality of the data (Figure S14D).

Class 7-1: Details are shown in Figure S15. The class average from the initial position sorting (Figure S15A, hot pink surface) contained traces of extra density along the entire density map (Figure S15A, dark grey, cornflower, light green, deep pink, peach, light grey, sienna, dark brown, teal, and khaki arrows), indicating pronounced heterogeneity within the class. To characterize it, 11 templates were generated using the initial class average (Figure S15A, hot-pink surface) as a guide. Template 1 (Figure S15B, cyan surface) contained one HMM molecule bound to actins 7 and 1 (Figure S15B, sienna arrow). In addition we generated nine templates for multiple HMMs bound to TF: template 1 (Figure S15B, blue surface, sienna and light green arrows), template 2 (Figure S15B, purple surface, sienna and deep pink arrows), 3 (Figure S15B, orange surface, sienna and khaki arrows), 4 (Figure S15B, dark green surface, sienna and dark brown arrows), template 5 (Figure S15B, yellow surface, sienna and teal arrows), 6 (Figure S15B, red surface, sienna and dark grey arrows), 7 (Figure S15B, green surface, sienna and peach arrows), 8 (Figure S15B, lavender surface, sienna and light grey arrows), and template 9 (Figure S15B, magenta surface, sienna and cornflower arrows). There was one triple HMM occupancy mode (Figure S15B, gold surface, light green, sienna, and dark grey arrows).

The templates were low-pass filtered to 15 Å and used for supervised 3D sorting in RELION (Figure S15B, solid surfaces). The frequencies of resultant modes were as follows: single HMM occupancy (n=66,037) (Figure S15C, cyan surface), double HMM occupancy mode 1 (n=988) (Figure S15C, blue surface), double HMM occupancy mode 2 (n=1,248) (Figure S15C, purple surface), double HMM occupancy mode 3 (n=237) (Figure S15C, orange surface), double HMM occupancy mode 4 (n=1,460) (Figure S15C, dark green surface), double HMM occupancy mode 5 (n=509) (Figure S15C, yellow surface), double HMM occupancy mode 6 (n=1,537) (Figure S15C, red surface), double HMM occupancy mode 7 (n=507) (Figure S15C, green surface), double HMM occupancy mode 8 (n=1,347) (Figure S15C, lavender surface), double HMM occupancy mode 9 (n=1,089) (Figure S15C, magenta surface), and the triple HMM occupancy (n=56) (Figure S15C, gold surface). The class with a single HMM molecule (Figure S15C, cyan surface, red box) had two junction regions (Figure S15C, black asterisks), suggesting the set was a mixture of particles having HMM bound to either actin molecules 6-7 or actin molecules 7-1. Segments were sorted by the position of HMM (Figure S15D) to select particles with the HMM bound to actin molecules 7-1 (Figure S15D, cerulean surfaces). The sorting indicated that the class predominantly was comprised of segments having HMM bound to actins 6 and 7: HMM bound to actin 7-1 (n=19,798) (Figure S15E, cerulean surface) and HMM bound to actin molecules 6-7 (n=46,239) (Figure S15E, chartreuse surface). Due to the small size of the Class 7-1 set, we decided not to pursue the refinement of this class.

**Modeling:** Actin, Tm, HMM, and TnT1 were modeled separately before the refinement of the overall model. Modeling was done similarly for every class (e.g., Classes 6-7, 5-6, 2-3, and 1-2).

Actin modeling: To generate an atomic model of actin, four actin subunits were independently segmented from the focused refined map using the UCSF Chimera [8] segmenting tool [9]. The segmented maps were auto-sharpened using the built-in algorithm in PHENIX [10]. Actin subunits from the porcine cardiac actomyosin complex (PDB 7JH7) [11] were rigid-body docked into the sharpened maps using UCSF Chimera [8]. The four N-terminal residues were added to the models in UCSF Chimera [8] using the amino acid sequence (UniProt ID: B6VNT8). The atomic models were placed in ChimeraX [12] [13] and flexibly fit with the ISOLDE algorithm [14]. The models were then refined in PHENIX [10] using the real-space refinement algorithm to restore stereochemistry.

Tropomyosin and TnT1 modeling: To generate an initial structure of the cardiac porcine α-tropomyosin, we used its amino acid sequence (UniProt: P42639) and human α-tropomyosin from PDB 7KO5 [5] as a template. Human and porcine sequences are 98.59% identical (Figure S2B). The Tm junction region structure in the M-state was modeled using its structure in the $Ca^{2+}$-free state (PDB 8DD0 [15]) as a starting point. The initial model of Tm was rigid-body docked into the segmented Tm portion of the map using UCSF Chimera [8] and subsequently refined in Namdinator [16].

TnT1 helix was modeled using the Alphafold database (AF-A0A5G2Q8N0-F1-v6) [17, 18]. The TnT1 helix portion of the map was segmented, and the initial TnT1 model was docked into the map using UCSF Chimera [8] followed by the refinement in PHENIX [10].

HMM modeling: To generate an initial model of the cardiac myosin using homology modeling in SWISS MODEL [19], we used the porcine cardiac amino acid sequence (UniProt: P79293) and the structure of skeletal myosin (PDB 5H53) [20] as a template. The two sequences shared 77.65 % identity. The structure 5H53 was selected as a template because it included a larger portion of the rod as well as loop 2, which was missing in the cardiac actomyosin structure 7JH7 [11]. The ELC from the crystal structure PDB 6FSA [21] was used as a template for the porcine ventricular amino acid sequence (UniProt: F1SNW4) in SWISS MODEL [19]. The two amino acid sequences were 94.90% identical. The porcine cardiac myosin homology model was aligned to the motor domain in PDB 5H53 [20]. The porcine cardiac ventricular isoform homology models for the ELCs and RLCs were also aligned to their structural counterparts in 5H53 [20]. The components were combined into a single-headed myosin molecule, and rigid-body docked into either the trailing or the leading heads, which were independently segmented from the overall map using UCSF Chimera [8] segmenting tool [9]. The initial refinement was performed using Namdinator [16] to correct for the significant movement of the tail region into the experimental density maps. Next, the models were iteratively refined in ChimeraX using ISOLDE plugin [14] followed by real-space refinement in PHENIX [10].

Tn core modeling: We used the previously published model of the $Ca^{2+}$-bound fully activated state from the upper strand (PDB 8UZX) [3] as a template for the Tn core. The cardiac TnI homology model, with 10 additional N-terminal residues (i.e., residues 32-42), was generated using SWISS MODEL [19] and structurally aligned to the overall Tn core structure (PDB 8UZX [3]). Next, the initial Tn core structure was rigid-body docked into the segmented map using UCSF Chimera [8], followed by the real-space refinement in PHENIX [10].

All the components (i.e., actin, Tm junction, TnT1 helix, myosin, Tn core, and ELC) were combined into a single model for each class. The parameters for the resultant models for each class are shown in Table S2.

<u>Visualization of the RLCs of the HMM molecule bound to actin molecules 6 and 7</u>: The polished consensus map was generated before the focused classification of the HMM RLC region (Figure S7D and S18A, grey surfaces). The HMM bound to actins 6 and 7, along with the portion of the Tm junction and TnT1 helix (Figure S18A, cerulean surface), was segmented from the consensus map using UCSF Chimera [8] segmenting tool [9] and used in the RELION particle subtraction routine to obtain 86,644 particles representing that region. The segmented map was filtered to 15 Å and used to generate a reference and a mask (Figure S18B, cerulean surface and grey mesh, respectively) for an overall 3D refinement and post-processing (Figure S18B, grey surfaces) using 86,644 subtracted particles. The densities that corresponded to RLCs were not sufficient to accommodate the RLCs' molecules (Figure S18C, inserts, light green ribbons), presumably due to RLCs' disordering, since the RLCs were unresolved at both the N and C-lobes (Figure S18D, black arrows). Therefore, we used the overall 3D map, filtered to 15 Å, as a reference and a mask (Figure S18E, dark green surface and grey mesh, respectively) for the unsupervised sorting of the subtracted particles. The resultant class averages and corresponding frequencies for the three classes are shown in Figure S18F: class 1 (n=42,634) (Figure S19F, green surface), class 2 (n=17,133) (Figure S18F,

chartreuse surface), and class 3 (n=26,877) (Figure S18F, teal surface). Classes 2 and 3 were discarded because they lacked a large portion of the RLC density. Class 1 was used for 3D refinement, using the 3D class-average low-pass filtered to 15 Å as the starting reference and mask (Figure S18G, green surface and grey mesh, respectively). The 3D reconstruction of Class1 RLCs showed that RLC densities were still incomplete (Figure S18H, inserts, light green ribbons) since the N and C lobes remained partially unresolved (Figure S18I, black arrows). The particles from class 1 (n=42,634) were used as the input for the second iteration of unsupervised classification. The class average was filtered to 15 Å and used as the reference and mask (Figure S18J, dark red surface and grey mesh, respectively). The resulting class averages and corresponding frequencies from the 3-class sorting are shown in Figure S18K: class 1 (n=10,625) (red surface), class 2 (n=29,660) (dark orange surface), and class 3 (n=2,349) (orange surface). Classes 1 and 3 were discarded due to partially missing RLC densities, while class 2 was low-pass filtered to 15 Å and used as the starting reference and mask (Figure S18L, dark orange surface and grey mesh, respectively) for the 3D refinement. In the resultant map, the trailing RLC was semi-disordered (Figure S18M, inserts, light green ribbons), since its N and C lobes were not fully resolved (Figure S18N, black arrows) while the leading RLC was resolved. The particles from class 2 (n=29,660) were used as the input for the third iteration of unsupervised sorting. The class average was again filtered to 15 Å and used as the reference and mask (Figure S18O, navy blue surface and grey mesh, respectively). The resulting class averages from the 3-class sorting are shown in Figure S18P: class 1 (n=2,192) (blue surface), class 2 (n=1,265) (cyan surface), and class 3 (n=26,203) (light blue surface). The class 3 class average was low-pass filtered to 12 Å and used as the starting reference and mask (Figure S18Q, light blue surface and grey mesh) for 3D refinement. The analysis of the RLC densities (Figure S18R, insert, light green ribbons) showed that the C-lobe of the trailing head was fully resolved, but the N-lobe was partially missing (Figure S18S, black arrow). The last unsupervised sorting was started from the low-pass-filtered class average as the reference and the mask (Figure S18T, deep pink surface and grey mesh, respectively). The class output maps from the final 3-class sorting are shown in Figure S18U: class 1 (n=8,153) (magenta surface), class 2 (n=175) (hot pink surface), and class 3 (n=17,875) (light pink surface). The 3D refinement was performed using the same approach as described previously, with the class-average filtered to 15 Å as the model and mask (Figure S18V, light pink surface and grey mesh). The final map showed the secondary structures of both trailing and leading RLCs (Figure S18W, inserts) and was used to generate an atomic model of the HMM that included the neck region (Figure S18X, green ribbons). Given the high heterogeneity of the RLCs and the limited initial class sizes for myosin bound to actin molecules 5–6, 2–3, and 1–2, we chose not to pursue further classification of the RLCs in these subsets.

Regulatory light chain (RLC) modeling: The RLC from the thick filament PDB 8G4L [22] was used as a template for the porcine cardiac ventricular amino acid sequence (UniProt: Q8MHY0) in SWISS MODEL [19], as they share 96.39% identity. To build the HMM model with RLCs, the final map from unsupervised sorting (Figure S18W) was used as input. The trailing and leading heads were independently segmented from the map filtered to 6 Å using the UCSF Chimera [8] segmenting tool [9], and auto-sharpened using PHENIX [10]. The two models were rigid-body docked into the two sharpened maps and refined in Namdinator [16]. The high-resolution model of myosin motor domains and ELCs from

Class 6-7, which yielded a perfect fit to the 6 Å map, was annealed with the rod domain, and RLC coordinates obtained from the 6 Å map using the PHENIX [10] rebuild algorithm were used to reconnect residues 808 and 809. The parameters for the resultant model are shown in Table S2.

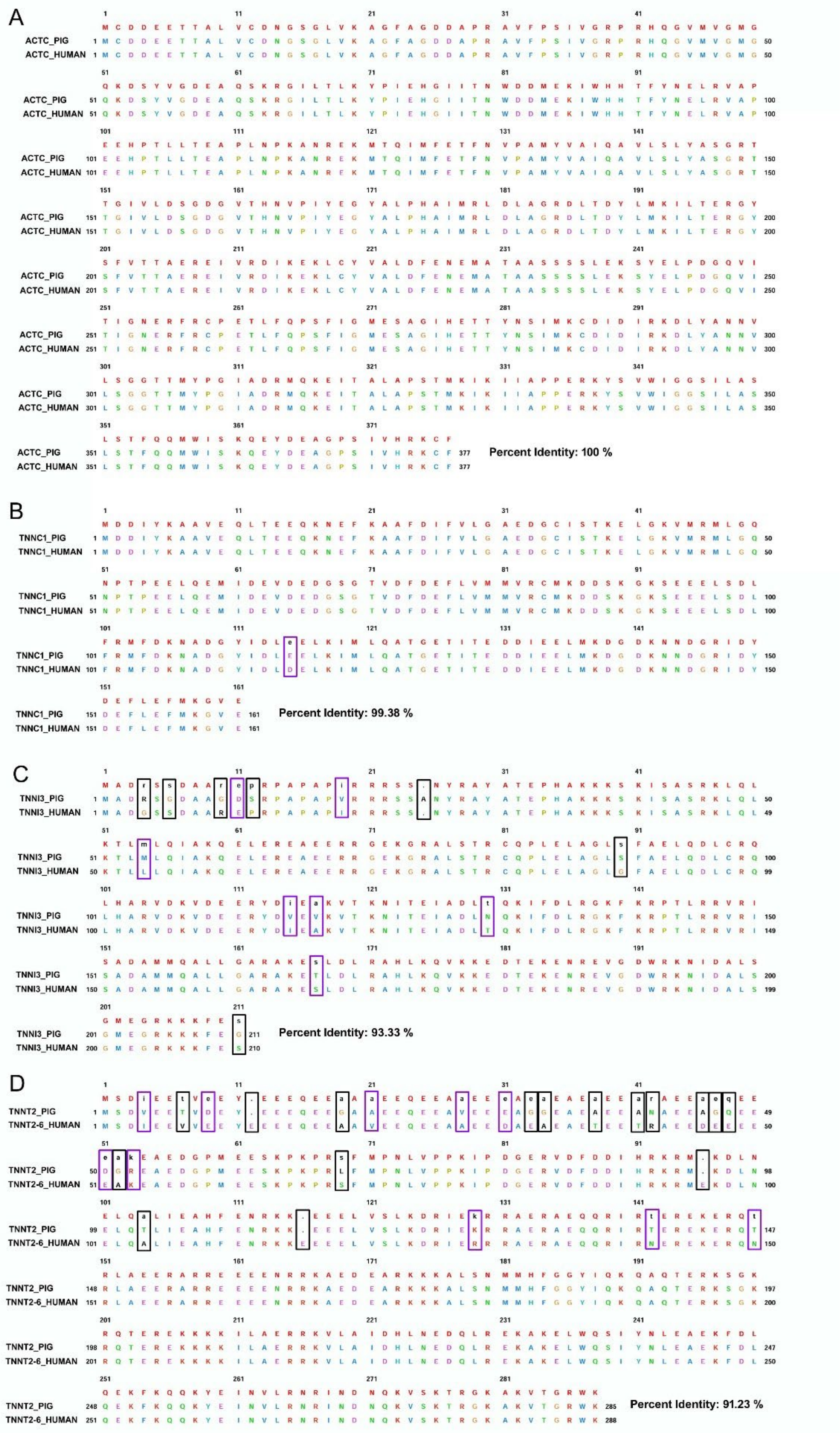
A
ACTC_PIG
ACTC_HUMAN
Percent Identity: 100 %
B
TNNC1_PIG
TNNC1_HUMAN
Percent Identity: 99.38 %
C
TNNI3_PIG
TNNI3_HUMAN
Percent Identity: 93.33 %
D
TNNT2_PIG
TNNT2-6_HUMAN
Percent Identity: 91.23 %

**Figure S1. Amino acid alignment of porcine and human cardiac actin, TnC, TnI, and TnT.** (A) Sequence alignment of porcine and human cardiac actin, which share 100% identity. (B) Sequence alignment of porcine and human cardiac TnC. There is one conserved substitution between the porcine and human isoforms, indicated with a purple rectangle. (C) Amino acid alignment of porcine and human cardiac TnI. Differences between the porcine and human sequences are marked with black and purple rectangles, respectively, for non-conserved and conserved substitutions. (D) Amino acid alignment of porcine and human cardiac TnT. Differences between the porcine and human sequences are marked the same as in (C).

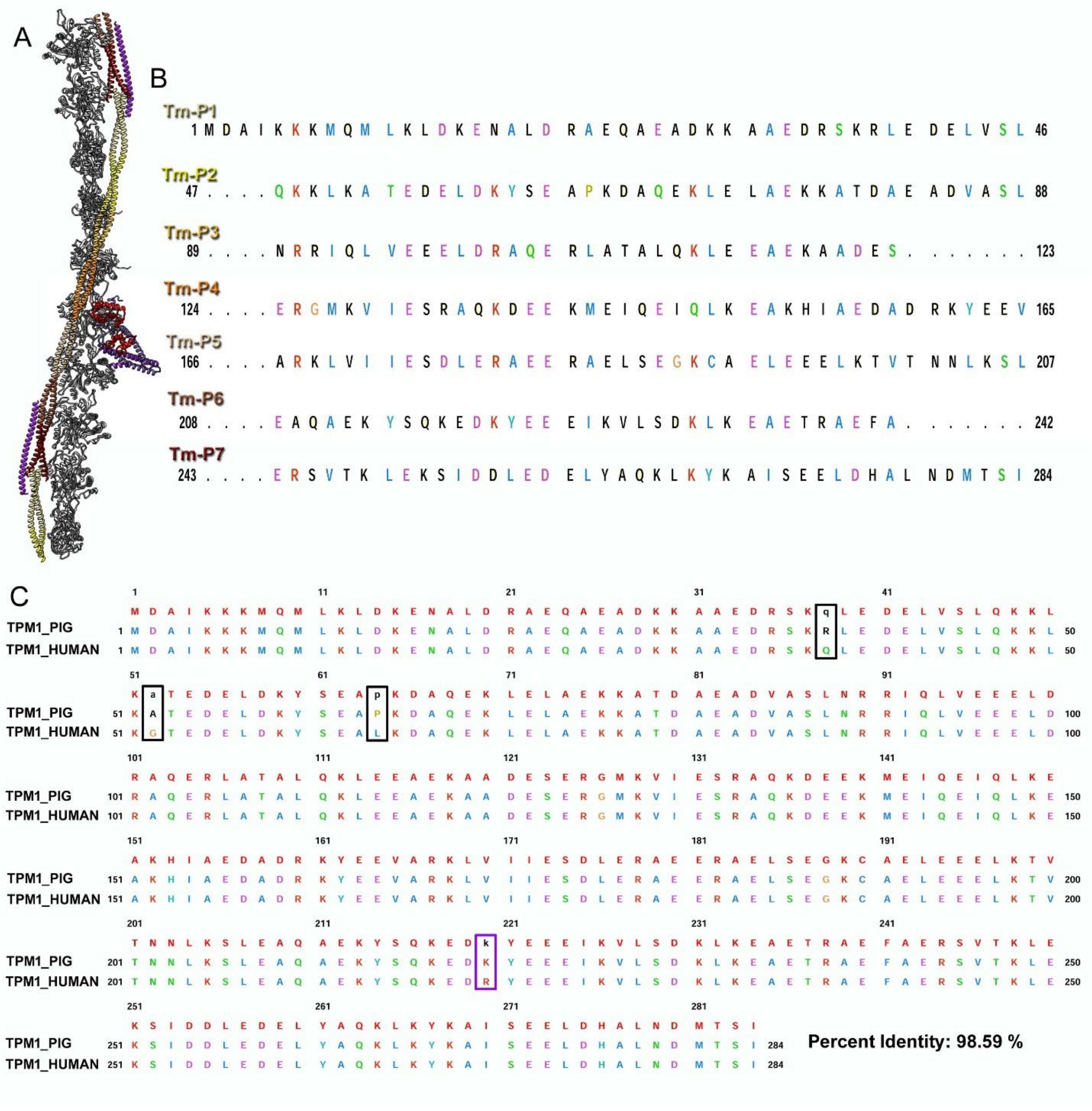


**Figure S2. Amino acid alignment of porcine and human cardiac α-Tm.** (A) Structure of the cTF RU, using the same color codes as main Figure 1. The components of the RU are actin (gray ribbons), TnT (purple ribbons), TnI (medium purple ribbons), TnC (red ribbons), and Tm (multicolored ribbon). The seven periods of Tm, starting with period 1, are shown as ribbons in the following color order: khaki, yellow, gold, orange, tan, sienna, and dark red. (B) Canonical sequence alignment of the seven period repeats of porcine cardiac Tm. (C) Sequence alignment of α-Tm from porcine and human species. Non-conserved differences are marked by a black rectangle, and conserved substitutions between porcine and human α-Tm are marked with a purple rectangle.

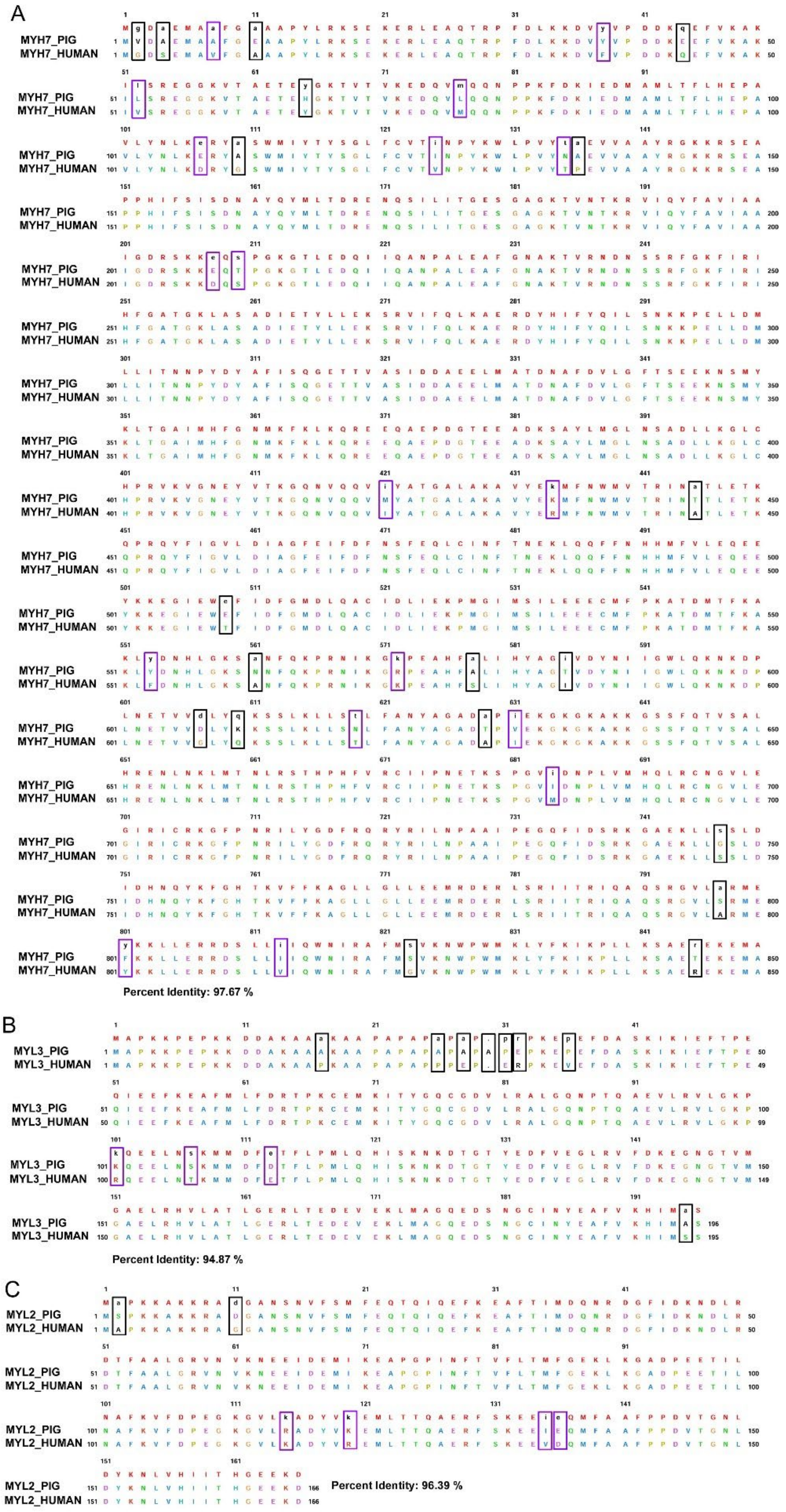
A
MYH7_PIG
MYH7_HUMAN
Percent Identity: 97.67 %
B
MYL3_PIG
MYL3_HUMAN
Percent Identity: 94.87 %
C
MYL2_PIG
MYL2_HUMAN
Percent Identity: 96.39 %

**Figure S3. Amino acid alignment of porcine and human cardiac myosin and essential and regulatory light chains.** (A) Sequence alignment of porcine and human cardiac myosin (residues 1-850). Differences between the porcine and human sequences are marked with black and purple rectangles, respectively, for non-conserved and conserved substitutions. (B) Sequence alignment of porcine and human cardiac ventricular essential light chain. Differences between the porcine and human sequences are marked the same as in (A). (C) Amino acid alignment of porcine and human cardiac ventricular regulatory light chains. Differences between the porcine and human sequences are marked the same as in (A) and (B).

**cryo-EM**

A

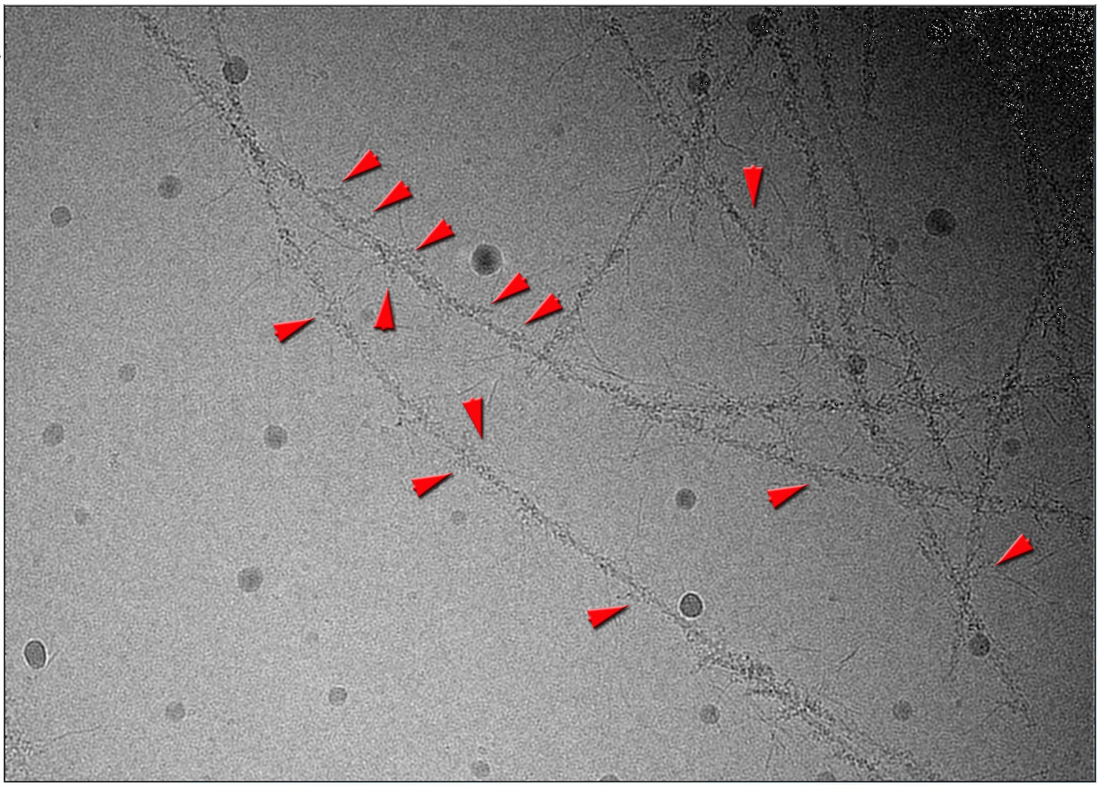

B

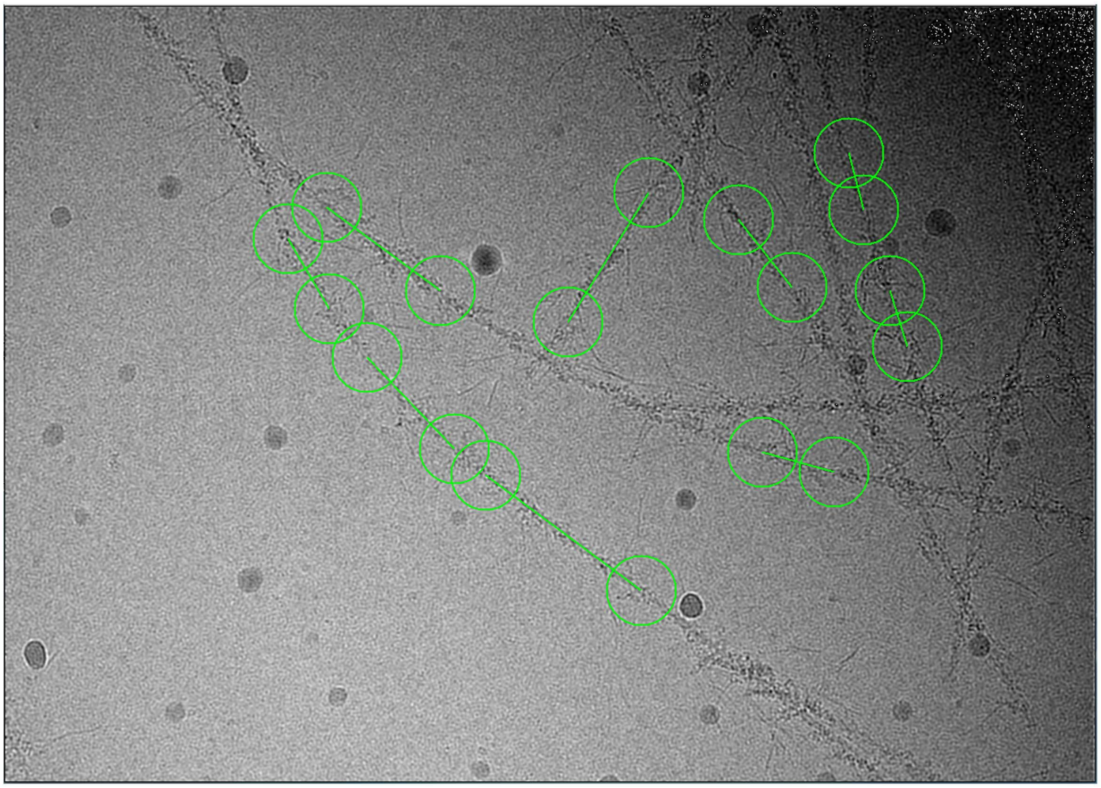

**Figure S4. Cryo-EM of cardiac native TFs decorated with HMM at pCa=4.** (A) Example cryo-EM micrograph of HMM-decorated cTFs. HMM molecules were clearly visible (red arrows), and areas of sparse decoration were used for manual selection. (B) Overlapping HMM-bound cTF segments were manually selected and extracted from the cryo-EM micrographs for 3D classification.

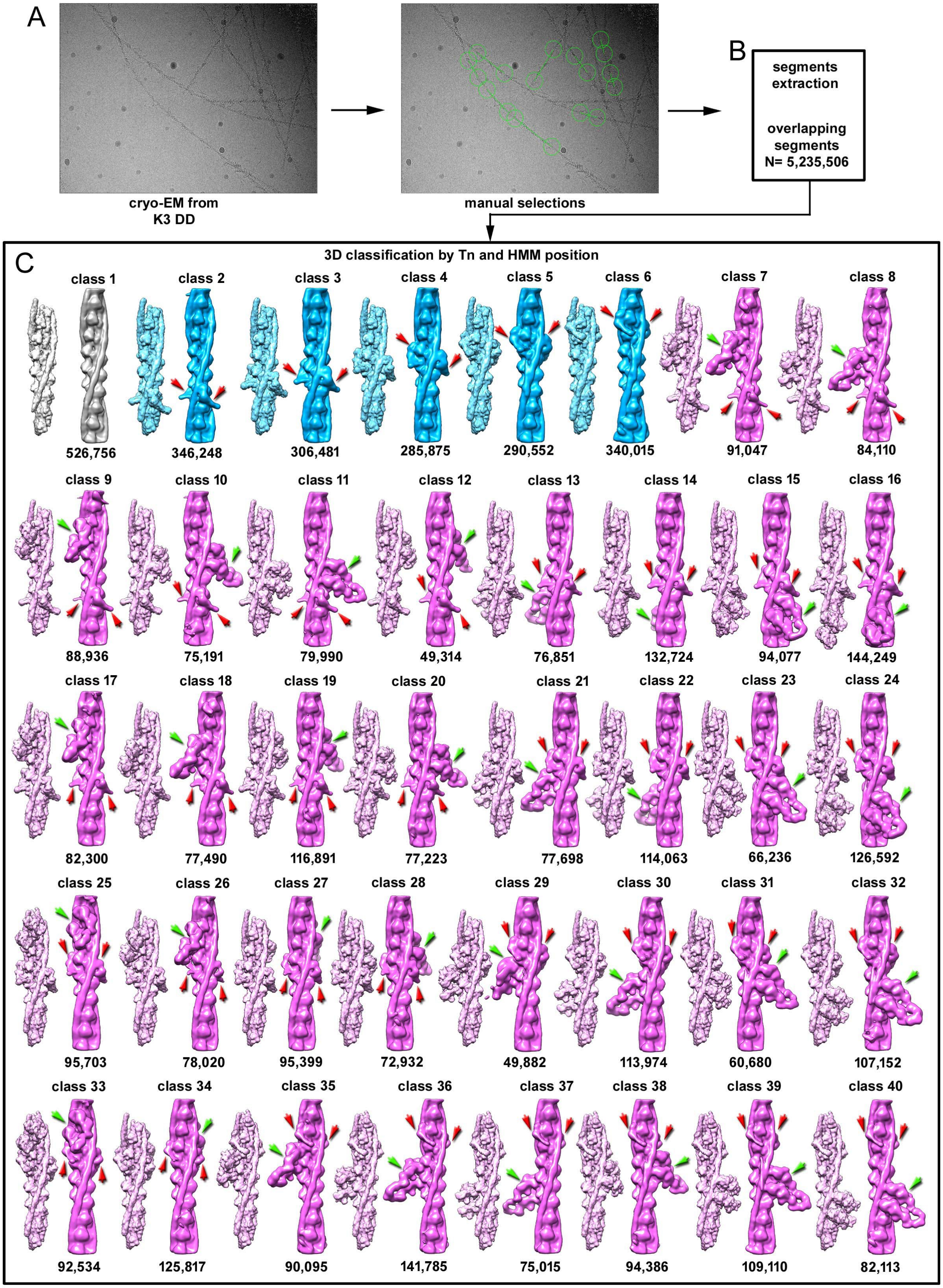
A
cryo-EM from
K3 DD
manual selections
B
segments
extraction
overlapping
segments
N= 5,235,506
C
3D classification by Tn and HMM position
class 1
526,756
class 2
346,248
class 3
306,481
class 4
285,875
class 5
290,552
class 6
340,015
class 7
91,047
class 8
84,110
class 9
88,936
class 10
75,191
class 11
79,990
class 12
49,314
class 13
76,851
class 14
132,724
class 15
94,077
class 16
144,249
class 17
82,300
class 18
77,490
class 19
116,891
class 20
77,223
class 21
77,698
class 22
114,063
class 23
66,236
class 24
126,592
class 25
95,703
class 26
78,020
class 27
95,399
class 28
72,932
class 29
49,882
class 30
113,974
class 31
60,680
class 32
107,152
class 33
92,534
class 34
125,817
class 35
90,095
class 36
141,785
class 37
75,015
class 38
94,386
class 39
109,110
class 40
82,113

**Figure S5. Initial classification of HMM-decorated cTF.** (A-B) Overlapping cTF segments were manually selected and extracted from cryo-EM micrographs for 3D classification (n = 5,235,506). (C) The extracted segments were sorted using a supervised method. Reference surfaces for sorting are shown as short maps in light gray (actin with Tm), light blue (TF without HMM), and light pink (TF with HMM is all possible positions). Of note, references lacked RLCs. The resulting class averages are shown next to the models as long surfaces using the following color codes: actin with Tm only (gray surface), TF without HMM bound (cerulean surfaces, Tn cores are marked with red arrows), TFs with HMM bound to different positions in regard to Tn core (pink surfaces; Tn cores indicated by red arrows, and HMM indicated by green arrows). The goal was to select segments containing the HMM molecule near the particle center (to avoid RLC clipping) that represent all available HMM positions on the RU. Particles from Class 10 (representing HMM bound to actins 1 and 2l Class 1-2), Class 11 (HMM bound to actins 2 and 3; Class 2-3), Class 31 (HMM bound to actins 5 and 6; Class 5-6), Class 39 (HMM bound to actins 6 and 7; Class 6-7), and Class 37 (HMM bound to actins 7 and 1; Class 7-1) were extracted for further processing depicted in Figures S7-S15 and S18.

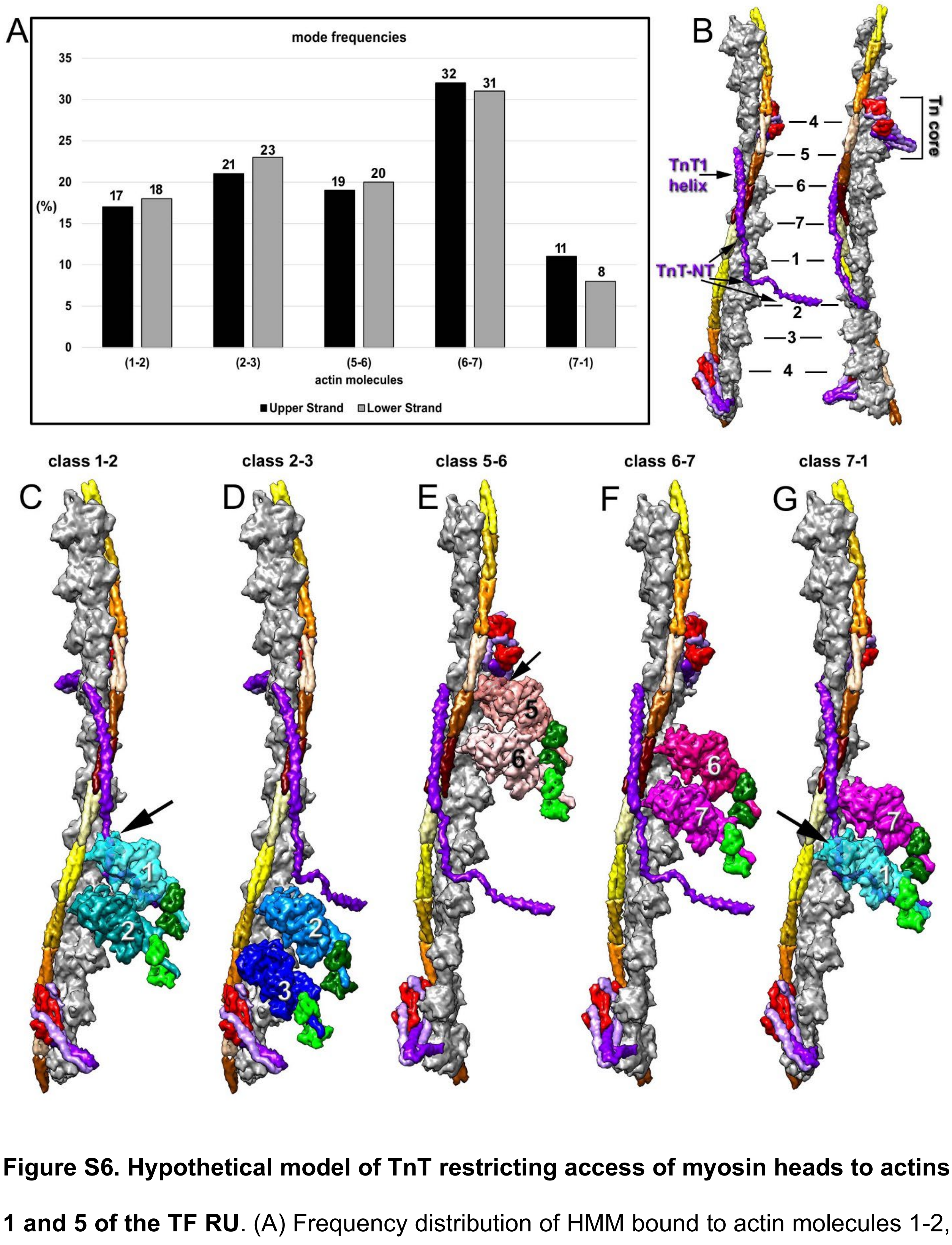


**Figure S6. Hypothetical model of TnT restricting access of myosin heads to actins 1 and 5 of the TF RU**. (A) Frequency distribution of HMM bound to actin molecules 1-2, 2-3, 5-6, 6-7, and 7-1 on the upper and lower strands. There are no significant differences in frequencies between the strands; however, the most populated class is HMM bound to actin molecules 6-7. (B) Model of the cardiac TF using the same color code as Figure 1. Actin molecules are numbered along the RU (black numbers). The full-length TnT is represented as a purple surface; however, the exact location of the N' and C' termini is hypothetical. (C-G) The hypothetical interference of TnT with myosin heads bound to actins 1-2, 2-3, 5-6, 6-7, and 7-1 is denoted with black arrows. (C) Class 1-2 trailing myosin head (cyan surface) is likely to have a major clash with the N-terminus of TnT (purple surface) at actin 1 (large black arrow). (E) Class 5-6 trailing myosin head (salmon surface) may have a minor clash with the Tn-core (small black arrow) at actin 5. (G) Class 7-1 myosin leading head (cyan surface) should have a clash with TnT at actin 1 (large black arrow), consistent with the low frequency of the class 7-1.

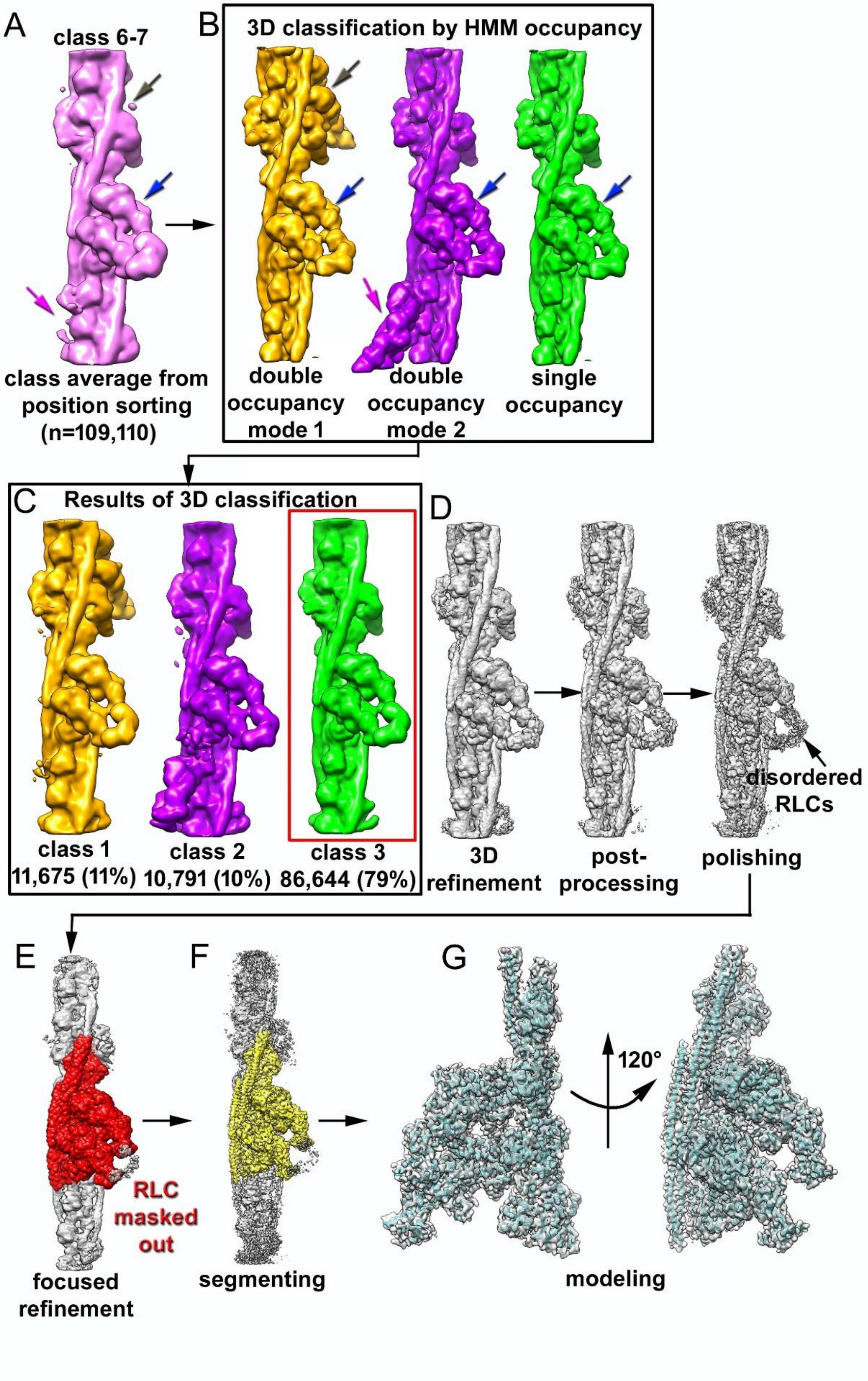

A
class 6-7
class average from position sorting (n=109,110)
B
3D classification by HMM occupancy
double occupancy mode 1
double occupancy mode 2
single occupancy
C
Results of 3D classification
class 1 11,675 (11%)
class 2 10,791 (10%)
class 3 86,644 (79%)
D
3D refinement
post-processing
polishing
disordered RLCs
E
RLC masked out
focused refinement
F
segmenting
G
120°
modeling

**Figure S7. 3D sorting and refinement of Class 6-7.** (A) The class 39 average (Figure S5C), shown as a hot pink surface, guided the construction of two templates that incorporated a second bound HMM molecule, based on HMM density traces in the class 39 average map (blue, magenta, and brown arrows). (B) Three templates for supervised sorting: mode 1 (orange surface, brown and blue arrows), mode 2 (purple surface, magenta and blue arrows), and single-occupancy HMM (green surface, blue arrow). (C) Resultant class averages and corresponding frequencies. Segments containing a single HMM molecule near the filament's center (red box) were used for 3D refinement and Bayesian polishing (D). (E) A focused refinement with a 10 Å mask (red surface) improved resolution in the region where the HMM binds to actin subunits 6 and 7 of TF RU. The HMM region was segmented (F) from the final map for modeling (G).

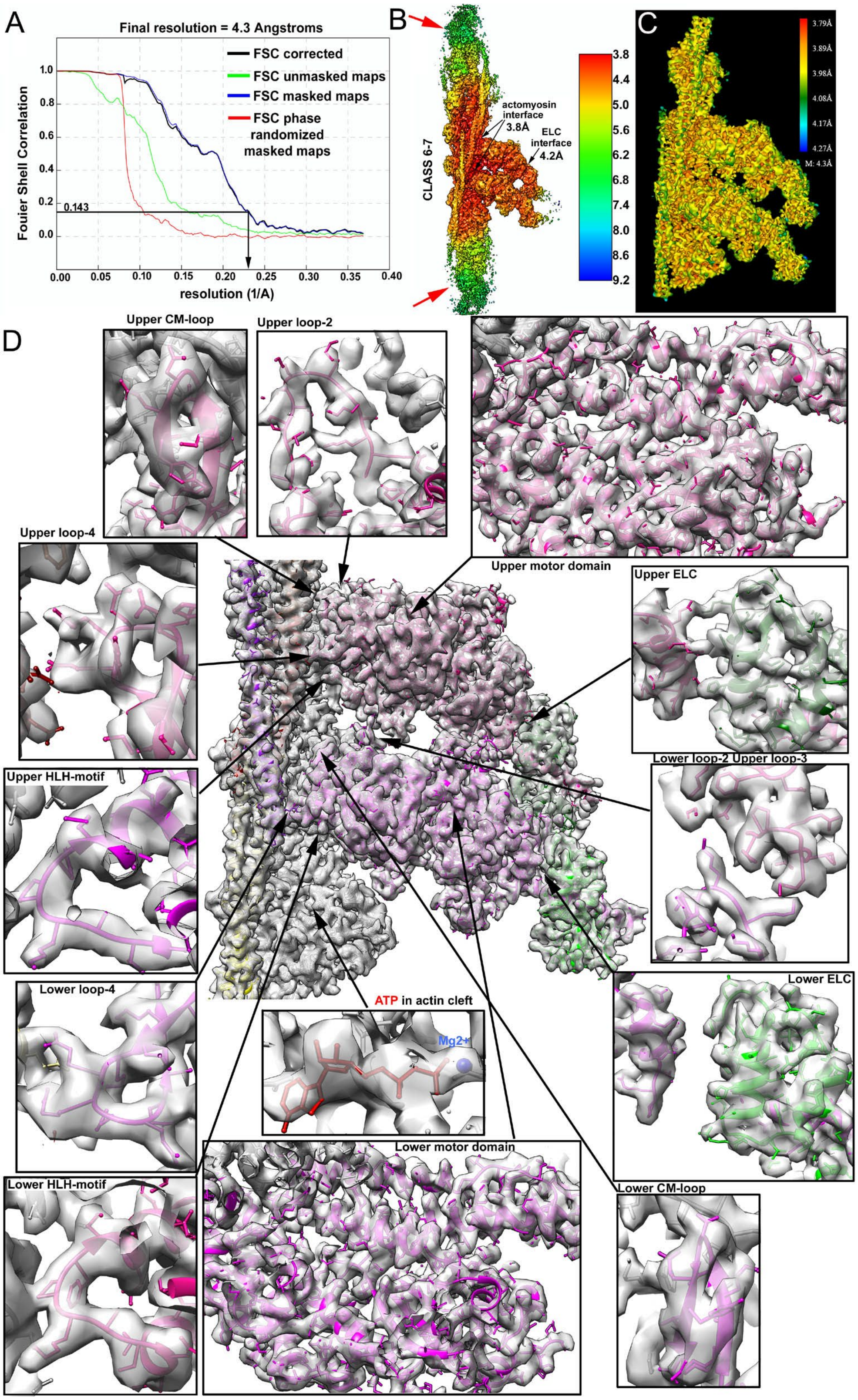
A
Final resolution = 4.3 Angstroms
FSC corrected
FSC unmasked maps
FSC masked maps
FSC phase randomized masked maps
Fouier Shell Correlation
0.143
resolution (1/A)
B
CLASS 6-7
actomyosin interface 3.8Å
ELC interface 4.2Å
C
3.79Å
3.89Å
3.98Å
4.08Å
4.17Å
4.27Å
M: 4.3Å
D
Upper CM-loop
Upper loop-2
Upper motor domain
Upper loop-4
Upper ELC
Upper HLH-motif
Lower loop-2 Upper loop-3
Lower loop-4
ATP in actin cleft
Mg2+
Lower ELC
Lower HLH-motif
Lower motor domain
Lower CM-loop

**Figure S8. Resolution estimation for Class 6-7.** (A) Fourier shell correlation (FSC) plot. The overall map resolution was calculated using the FSC criterion of 0.143. (B) Local resolution map determined by RELION implementation. The resolution is ~3.8 Å at the actomyosin interface and 4.2 Å at the ELC interface, while it drops to ~9.2 Å in the distal regions of the map (red arrows). (C) Local resolution estimation based on the final map using CryoRes. (D) 3D reconstruction of the HMM bound to actin molecules 6 and 7 is shown (grey surface) along with its corresponding model (colored atoms). Actin is shown in gray; period 1 of (Tm) is shown in khaki; period 2 is shown in yellow; period 6 is sienna; and period 7 is dark red. The TnT1 helix is shown in purple. The trailing myosin is deep pink, the trailing ELC is dark green, while the leading head is magenta, and the leading ELC is green. The high-resolution cryo-EM map reveals atomic details of the structural elements of the cardiac HMM involved in interactions with actin, Tm, ELC, and between trailing and leading myosin heads. Detailed views of the various components discussed in the paper are shown alongside the atomic model to illustrate the quality of the data.

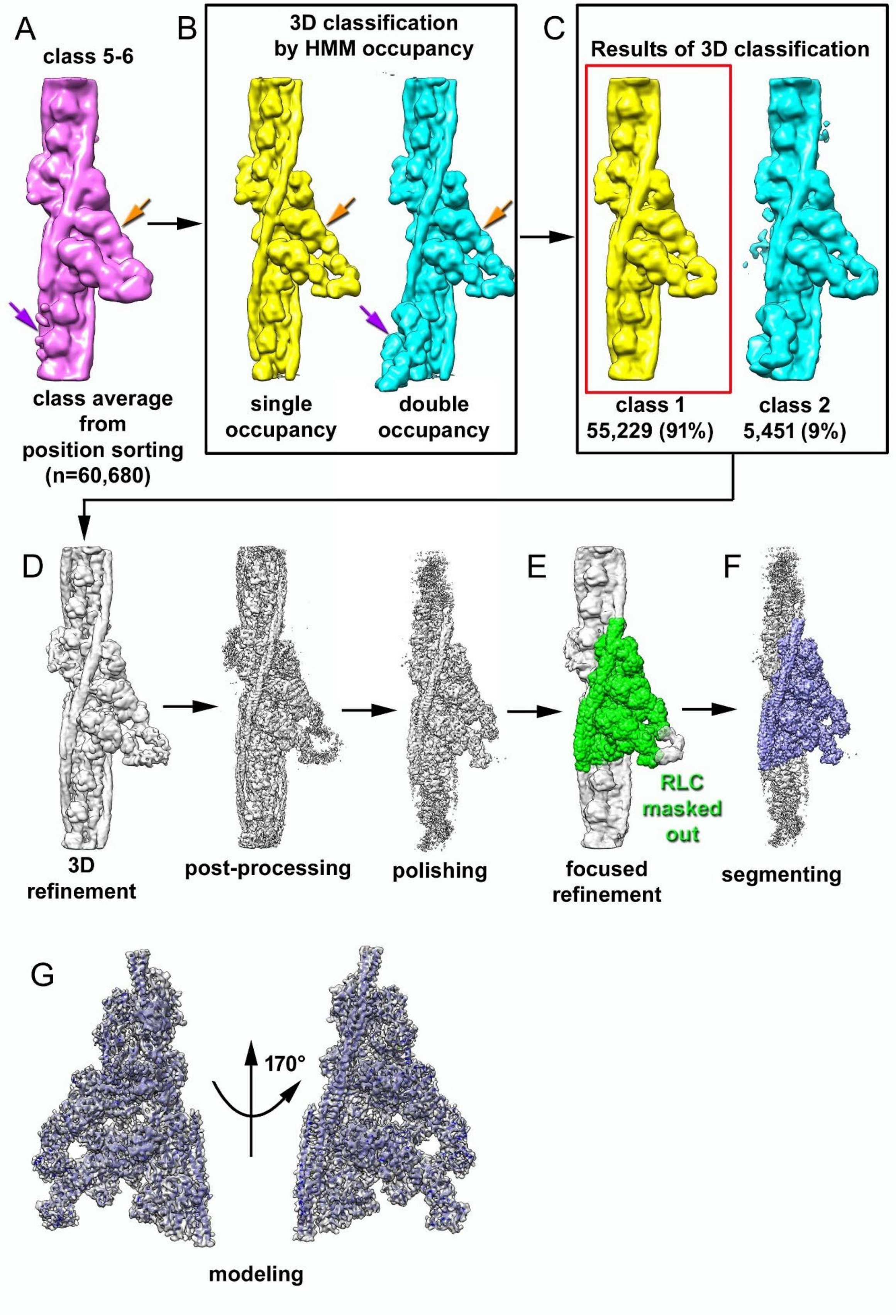

A
class 5-6
class average
from
position sorting
(n=60,680)
B
3D classification
by HMM occupancy
single
occupancy
double
occupancy
C
Results of 3D classification
class 1
55,229 (91%)
class 2
5,451 (9%)
D
3D
refinement
post-processing
polishing
E
RLC
masked
out
focused
refinement
F
segmenting
G
170°
modeling

**Figure S9. 3D sorting and refinement of Class 5-6.** (A and B) The class 31 average (Figure S5C), shown as a hot pink surface, guided the construction of two templates that incorporated either one or two bound HMM molecules, based on HMM density traces in the class 31 average map (purple and orange arrows) that are shown in (B) (yellow and cyan surfaces). (C) Resultant class averages and corresponding frequencies. The segments that possessed one HMM molecule near the center of the filament (yellow surface, red box) were used for 3D refinement and Bayesian polishing (D). (E) A focused refinement with a 10 Å mask (green surface) was used to improve the resolution in the region where HMM was bound to actin molecules 5 and 6. The HMM region was segmented (lavender surface) (F) from the final map for modeling (blue ribbons) (G).

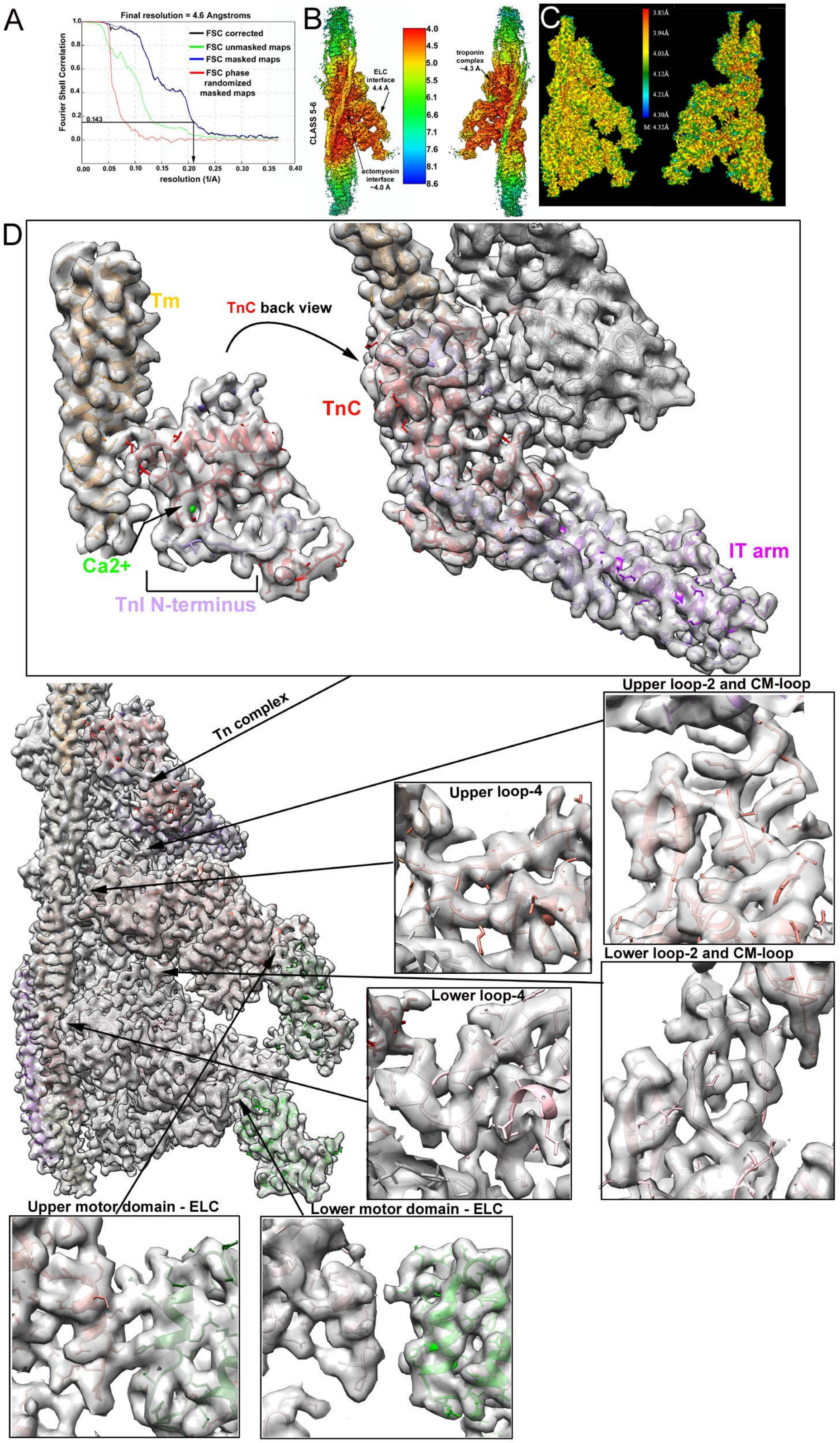

A
Final resolution = 4.6 Angstroms
Fourier Shell Correlation
resolution (1/A)
0.143
FSC corrected
FSC unmasked maps
FSC masked maps
FSC phase randomized masked maps
B
CLASS 5-6
ELC interface 4.4 Å
actomyosin interface ~4.0 Å
troponin complex ~4.3 Å
C
3.85Å
3.94Å
4.03Å
4.12Å
4.21Å
4.30Å
M: 4.32Å
D
Tm
TnC back view
TnC
Ca2+
TnI N-terminus
IT arm
Tn complex
Upper loop-2 and CM-loop
Upper loop-4
Lower loop-2 and CM-loop
Lower loop-4
Upper motor domain - ELC
Lower motor domain - ELC

**Figure S10. Resolution estimation for Class 5-6**. (A) Fourier shell correlation (FSC) plot. The overall resolution of the map was calculated using the FSC criterion of 0.143. (B) Local resolution map determined by the RELION implementation. The resolution is ~4.0 Å at the actomyosin interface, 4.4 Å at the ELC interface, and ~4.3 Å at the Tn core. (C) Local resolution estimated from the final map using CryoRes. (D) 3D reconstruction of the HMM bound to actin molecules 5 and 6 is shown (gray surface) along with its corresponding model (colored atoms). The color code is the same as Figure 1. The high-resolution cryo-EM map reveals atomic details of the structural elements of the cTF (i.e., Tn core, actin, and Tm) and of the cardiac HMM involved in interactions with actin, the Tn core, Tm, ELC, and between trailing and leading myosin heads. Detailed snapshots of the various components discussed in the paper are shown along with the atomic model.

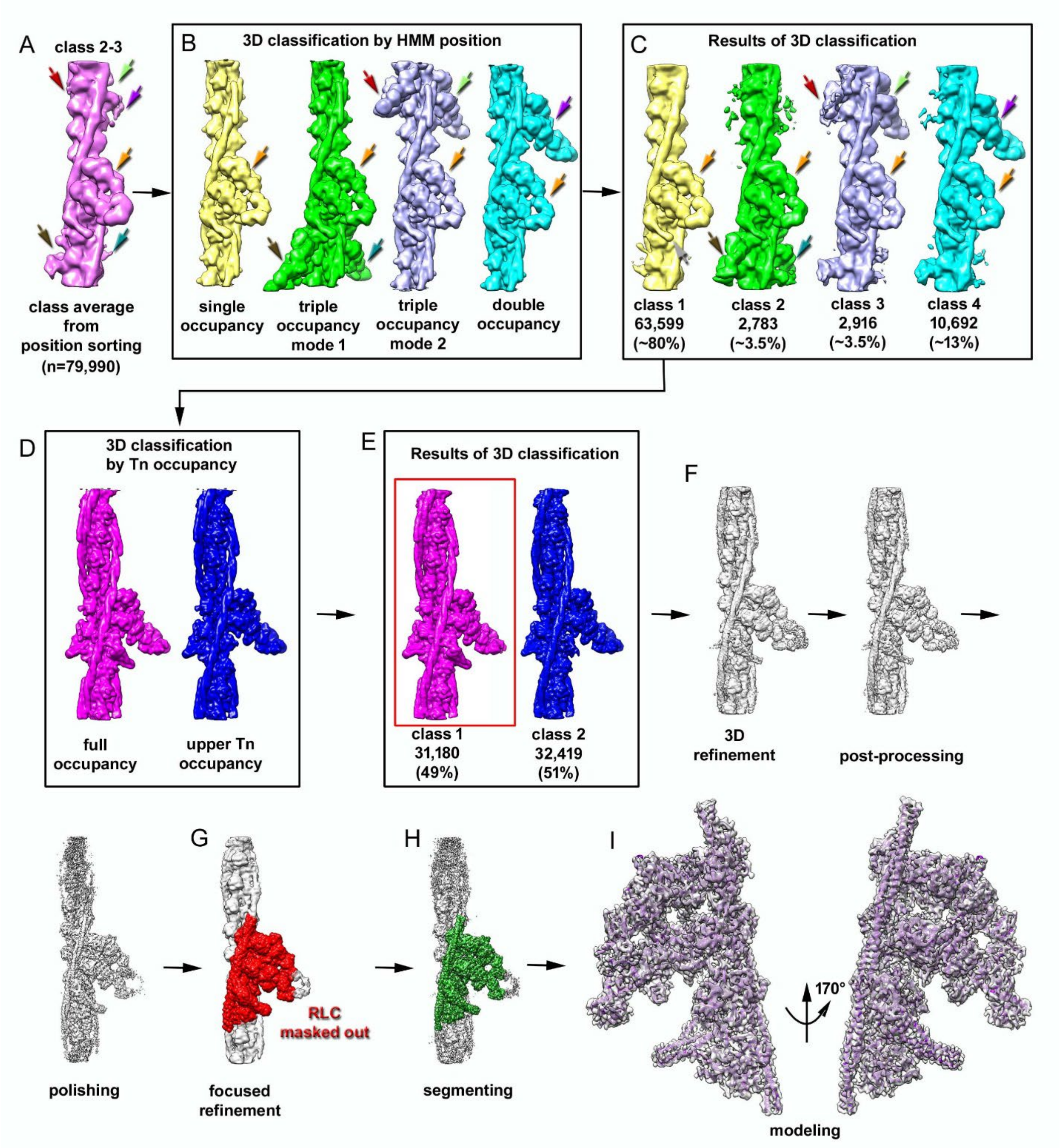


**Figure S11. 3D sorting and refinement of Class 2-3.** (A) The class 11 (Figure S5C) average from the initial position classification (depicted as a hot pink surface) was used as the guide to build templates for supervised sorting based on traces of additional HMM molecules bound (dark red, chartreuse, purple, orange, brown, and teal arrows). (B) The resultant four templates have one (yellow surface), two (cyan surface), or three (green and lavender surfaces) HMM molecules bound. (C) Resultant class averages and corresponding frequencies. The segments that contained one HMM molecule near the center of the filament (yellow surface) had insufficient density for the Tn core (grey arrow) and were sorted based on the integrity of the Tn tandem (D) – one template had both Tn cores (magenta), whereas the other had one Tn complex missing (blue). (E) Resultant class averages where the intact Tn tandem (magenta surface, red box) was re-extracted at a scale of 1.356 Apx and used for 3D refinement and Bayesian polishing (F). (G) A focused refinement with a 10 Å mask (red surface) was used to improve the resolution in the region where HMM was bound to actin molecules 2 and 3. The HMM region (dark green surface) (H) was segmented from the final map for modeling (purple ribbons) (I).

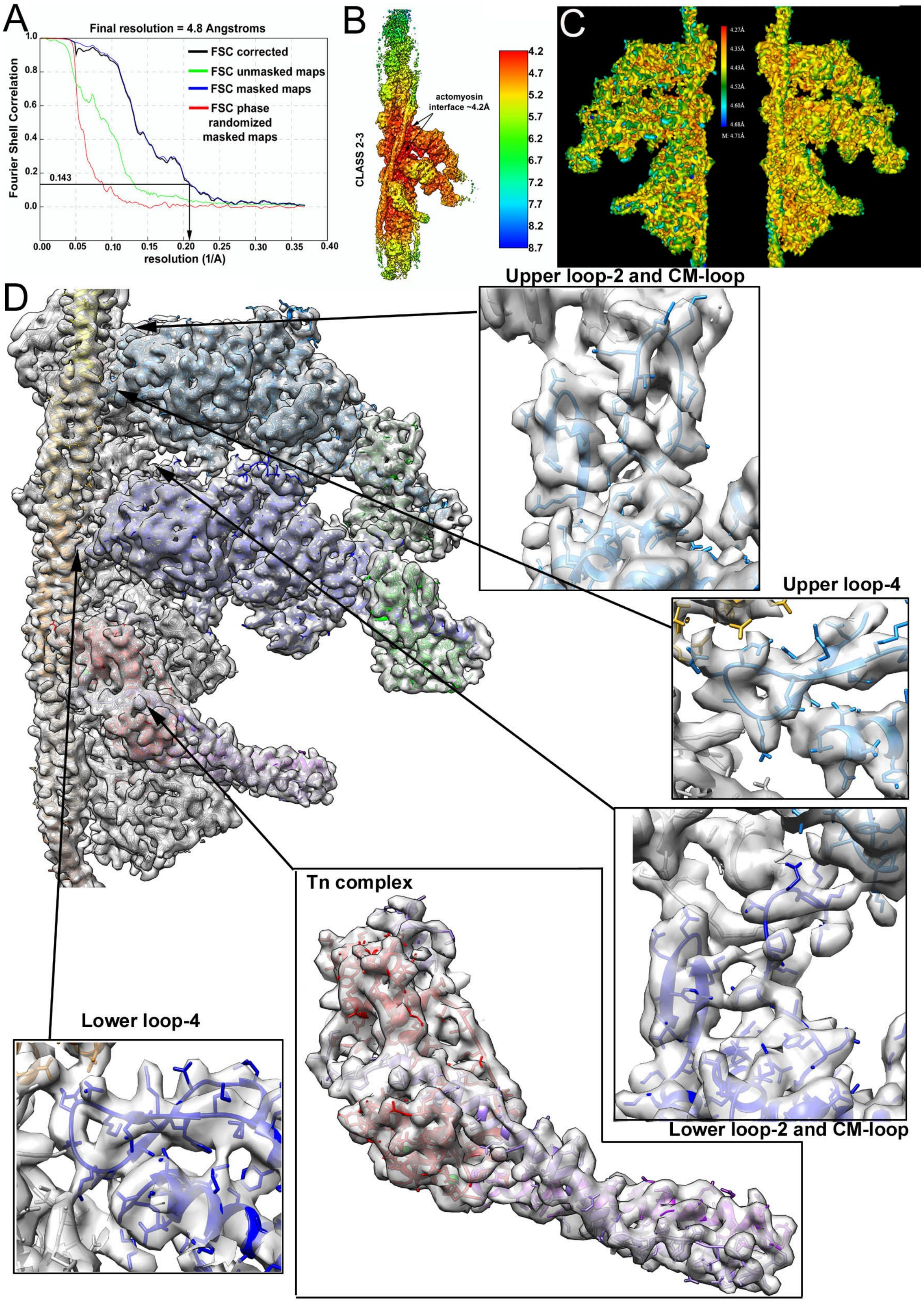

A
Final resolution = 4.8 Angstroms
FSC corrected
FSC unmasked maps
FSC masked maps
FSC phase randomized masked maps
Fourier Shell Correlation
resolution (1/A)
0.143
B
CLASS 2-3
actomyosin interface ~4.2Å
C
D
Upper loop-2 and CM-loop
Upper loop-4
Lower loop-2 and CM-loop
Tn complex
Lower loop-4

**Figure S12. Resolution estimation for Class 2-3.** (A) Fourier shell correlation (FSC) plot. The overall resolution for the map was calculated using the gold standard FSC criterion of 0.143. (B) Local resolution map determined by RELION implementation. The resolution is ~4.2 Å at the actomyosin interface. (C) Local resolution calculated from the final map using CryoRes. (D) 3D reconstruction of the HMM bound to actin molecules 2 and 3 is shown (grey surface) along with its corresponding model (colored atoms). The color code is the same as Figure 1. The cryo-EM map shows the atomic details of the structural elements of the cTF and cardiac HMM involved in interactions with actin, Tn core, Tm, ELC, and interactions between trailing and leading myosin heads. Detailed views of the components discussed in the paper are shown alongside the atomic model.

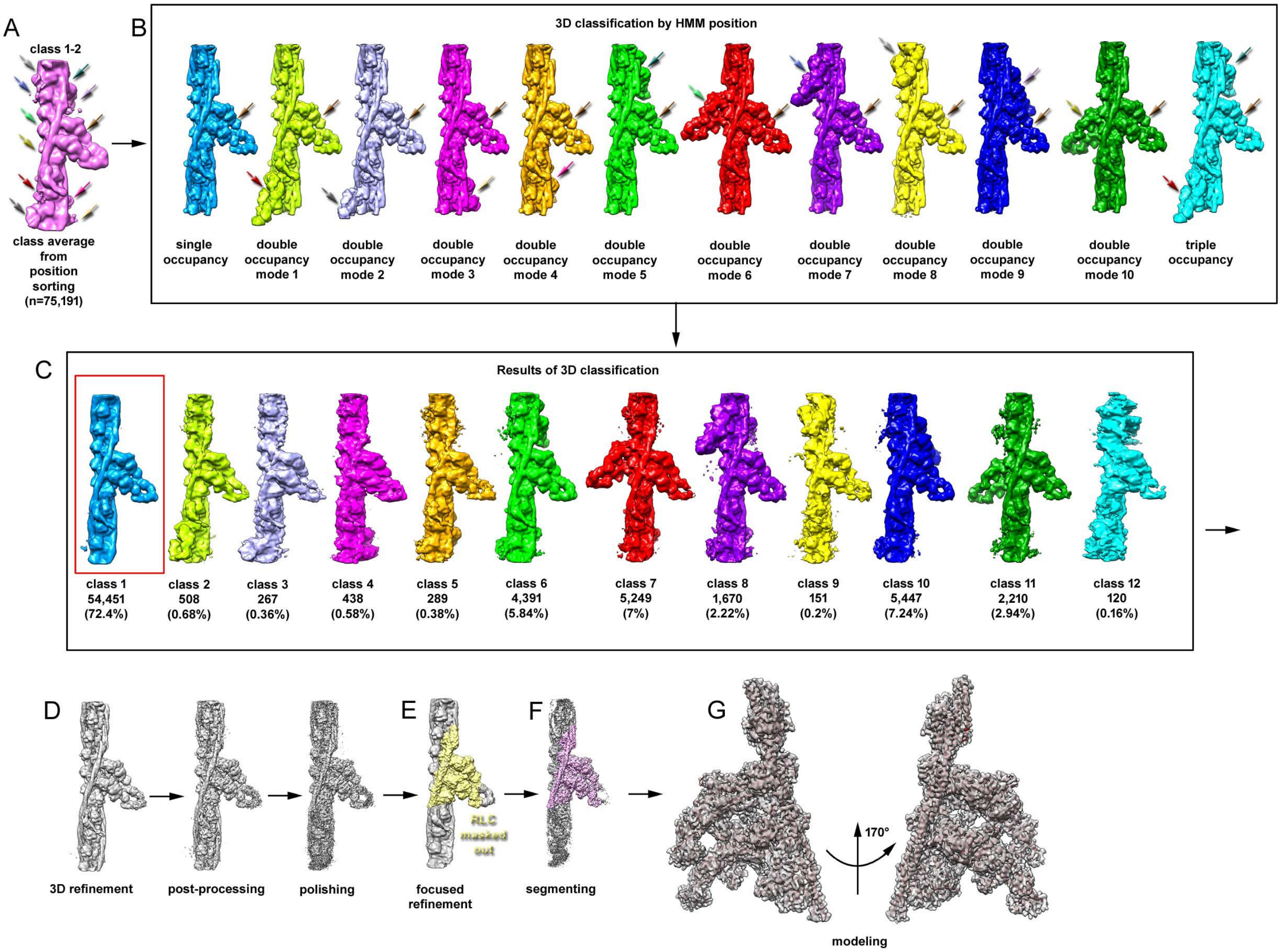


**Figure S13. 3D sorting and refinement of Class 1-2.** (A) The class 10 output map from the initial position classification (Figure S5C), shown as a hot pink surface, served as a guide for building templates with HMM bound to multiple positions, based on HMM traces revealed in the class 10 map (light gray, cornflower, light green, gold, dark red, dark gray, teal, light purple, sienna, deep pink, and khaki arrows). (B) Twelve templates were generated - one single-occupancy HMM (cerulean surface), ten double-occupancy modes (chartreuse, lavender, magenta, orange, green, red, purple, yellow, blue, and dark green surfaces), and one triple-occupancy HMM mode (cyan surface). (C) Resultant class averages and associated frequencies. Segments with one HMM molecule near the center of the filament (cerulean surface, red box) were re-extracted and used for 3D refinement and Bayesian polishing (D). (E) A focused refinement with the 10 Å mask (light yellow surface) was used to improve resolution in the region where HMM was bound to actin molecules 1 and 2. The HMM region was segmented (light pink surface) (F) from the final map for modeling (dark brown ribbons) (G).

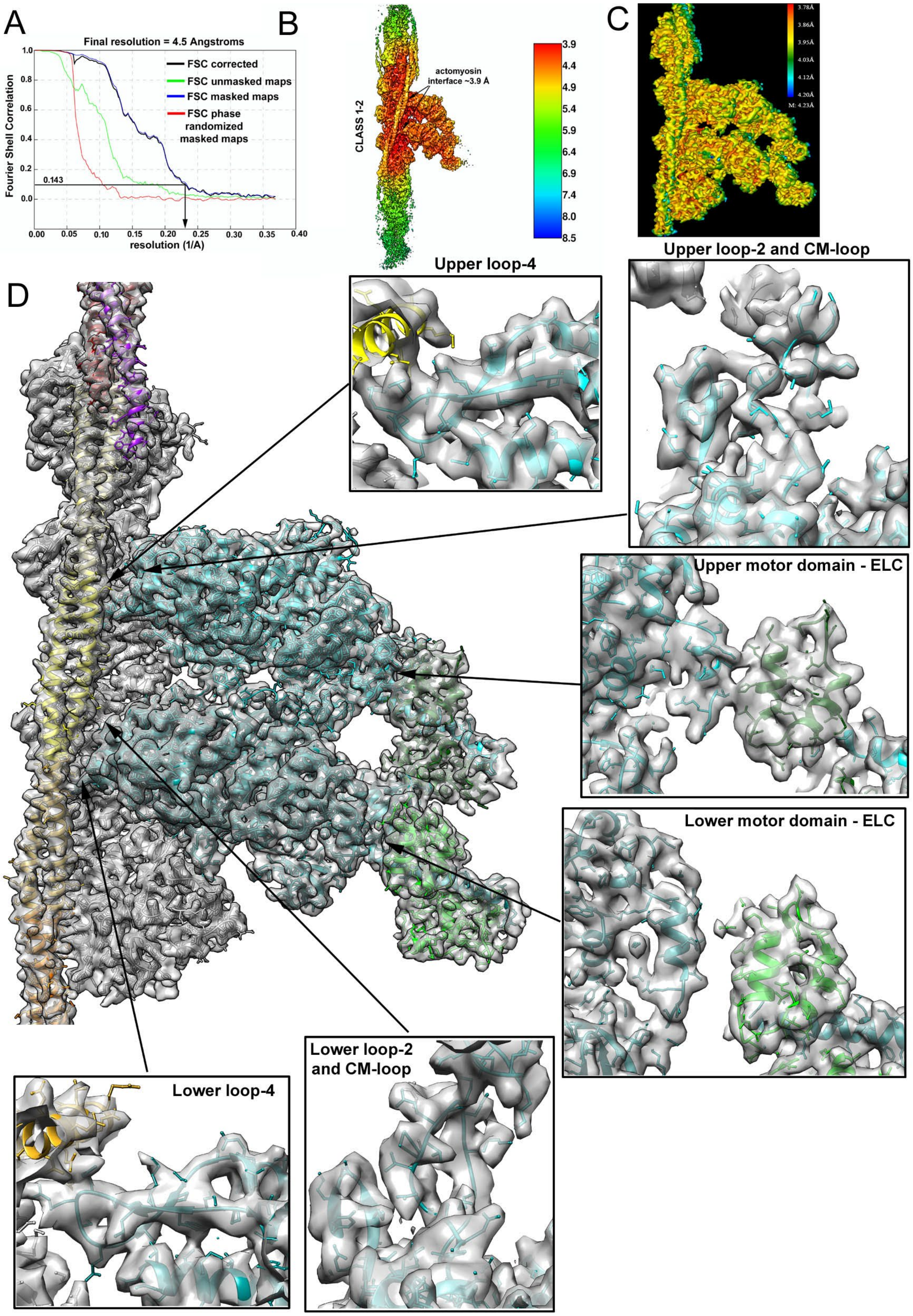
A
Final resolution = 4.5 Angstroms
Fourier Shell Correlation
FSC corrected
FSC unmasked maps
FSC masked maps
FSC phase randomized masked maps
0.143
resolution (1/A)
B
CLASS 1-2
actomyosin interface ~3.9 Å
3.9
4.4
4.9
5.4
5.9
6.4
6.9
7.4
8.0
8.5
C
D
Upper loop-4
Upper loop-2 and CM-loop
Upper motor domain - ELC
Lower motor domain - ELC
Lower loop-2 and CM-loop
Lower loop-4

**Figure S14. Resolution estimation for class 1-2.** (A) Fourier shell correlation (FSC) plot. The global resolution for the map was calculated using an FSC criterion of 0.143. (B) Local resolution map determined by RELION. The resolution is ~3.9 Å at the actomyosin interface. (C) Local resolution estimated from the final map using CryoRes. (D) 3D reconstruction of the HMM bound to actin molecules 1 and 2 is shown (gray surface) along with its corresponding model (colored atoms). The color code is the same as in Figure 1. The high-resolution cryo-EM map reveals atomic details of the structural elements of the cardiac HMM that interact with cTF and ELC, as well as interactions between the trailing and leading myosin heads. Detailed views of the regions discussed in the paper are shown, along with the corresponding atomic model, to demonstrate the quality of the data.

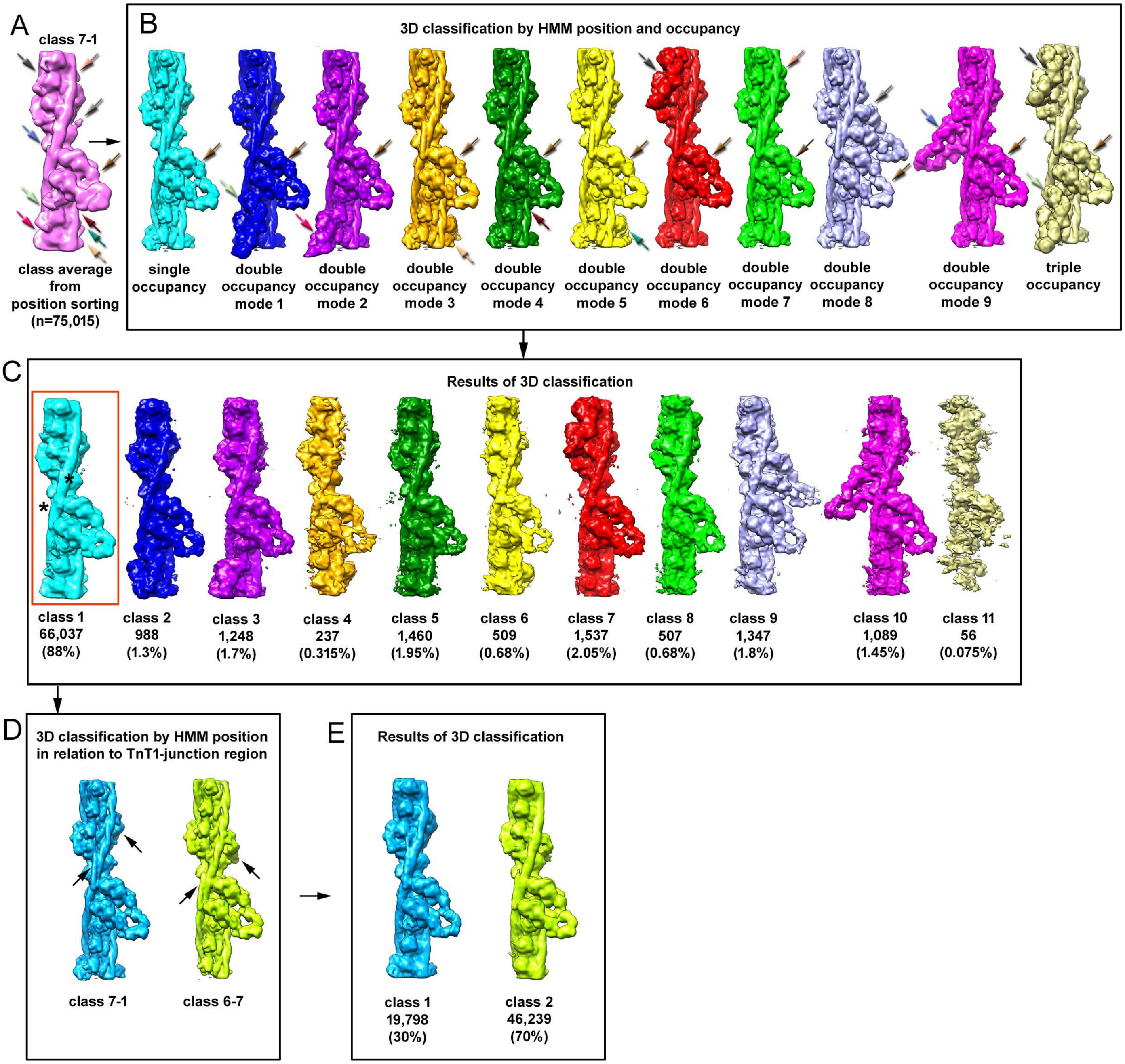


**Figure S15. 3D sorting and refinement of class 7-1.** (A) The class 37 output map from the initial position classification (Figure S5C), shown as a hot pink surface, was used as a guide to build templates with multiple HMM molecules bound, based on traces of additional HMM molecules (dark grey, cornflower, light green, deep pink, peach, light grey, sienna, dark brown, teal, and khaki arrows) revealed in the class 37 class average. (B) The eleven templates used for supervised sorting included one single-occupancy HMM (cyan surface), nine double-occupancy modes (blue, purple, orange, dark green, yellow, red, green, lavender, and magenta surfaces), and one triple-occupancy mode (gold surface). (C) Resultant class averages and corresponding frequencies. Segments with one HMM molecule near the center of the filament (cyan surface, red box) were used for further classification based on the positioning of Tm junctions (black asterisks). Two junction regions indicated a mixture of HMM-bound modes. (D) Supervised sorting based on the position of HMM in relation to the Tm junction region, with HMM bound to actin molecules 7-1 (cerulean surface and black arrow) or 6-7 (chartreuse surface and black arrow). (E) Resultant class averages and corresponding frequencies. Class 1, representing segments with HMM at the position of actin molecule 7 from one RU and actin molecule 1 on the adjacent RU, accounts for only 30% of segments.

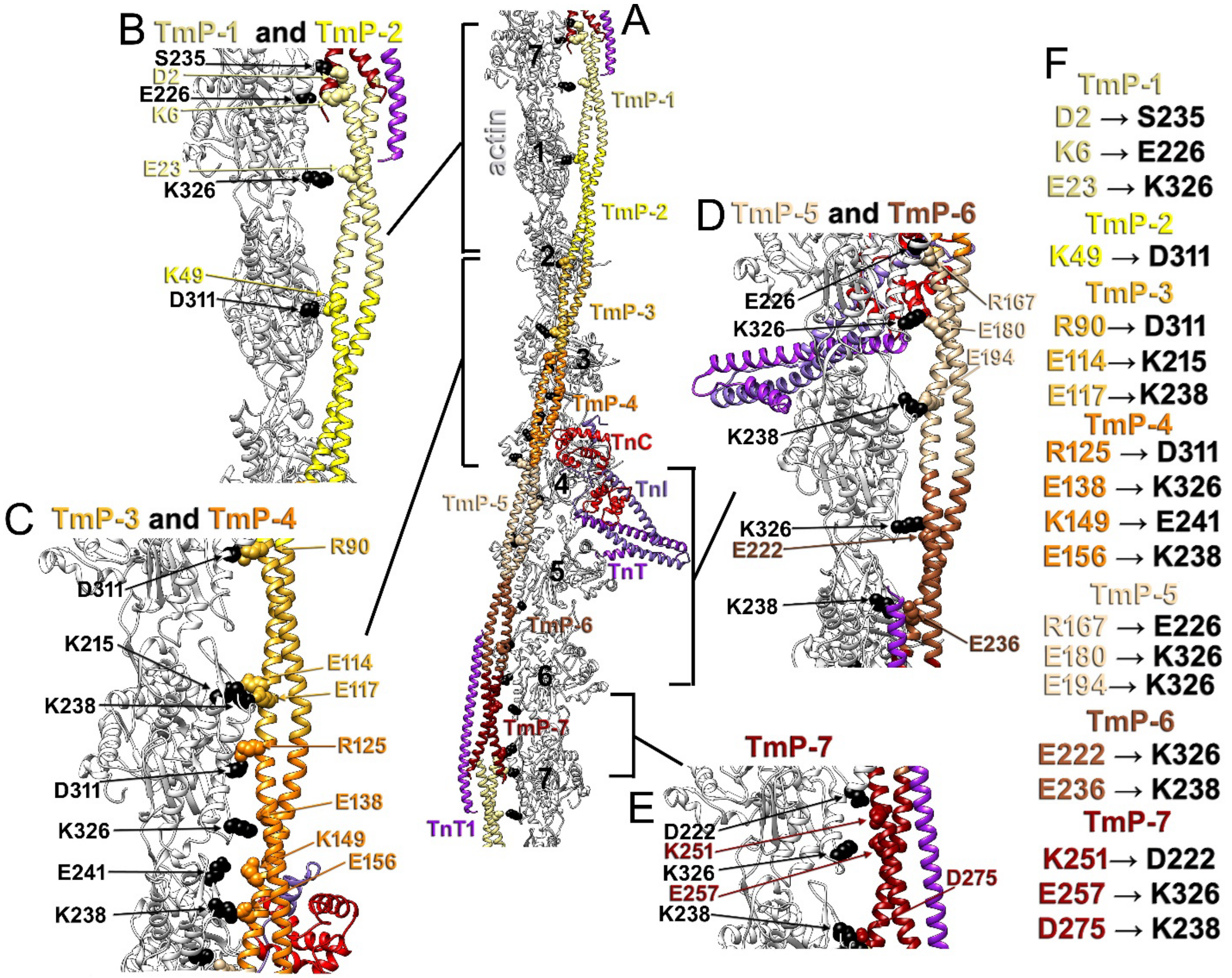


**Figure S16. Interactions between actin and tropomyosin repeats.** (A-E) Model of the cardiac TF using the same color code as in Figure 1. Actin residues that interact with Tm are shown as black spheres, while Tm residues involved in actin binding are shown as spheres colored according to the Tm period. (F) Interactions between actin and Tm periods are arranged in a table.

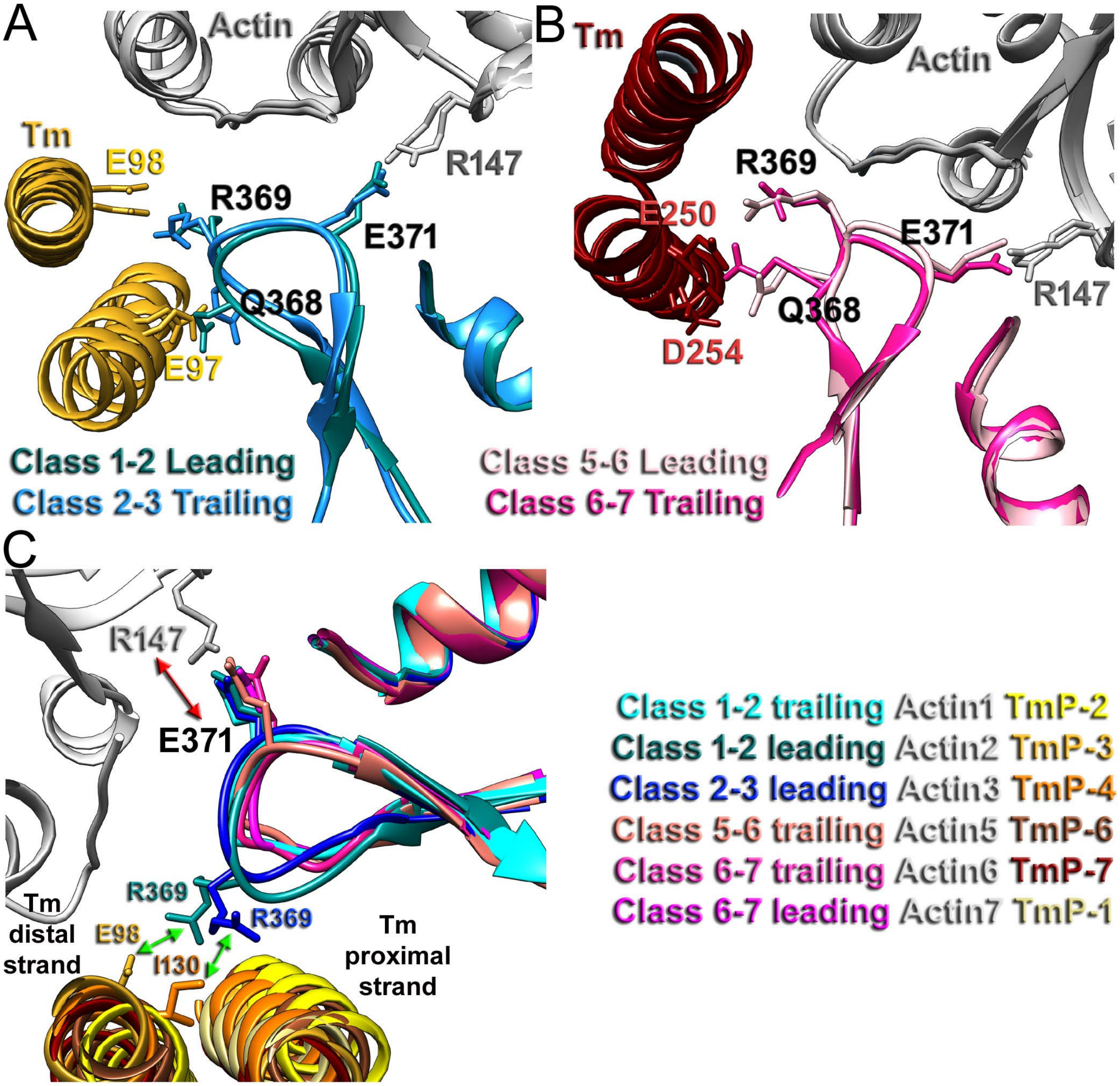


**Figure S17. Interactions of myosin loop-4 with actin and Tm.** (A) Loop-4 of the Class 1-2 leading head and loop-4 of the Class 2-3 trailing head form the same contacts with actin R147 (gray atoms) and Tm E97/E98 (gold atoms). (B) Loop-4 of the Class 5-6 leading head and loop-4 of the Class 6-7 trailing head form the same contacts with actin R147 (gray atoms) and Tm E250/D254 (dark red atoms). (C) The extended conformations of loop-4 from the Class 1-2 leading head and the Class 2-3 leading head correlate with their interactions with the far Tm strand (green arrows).

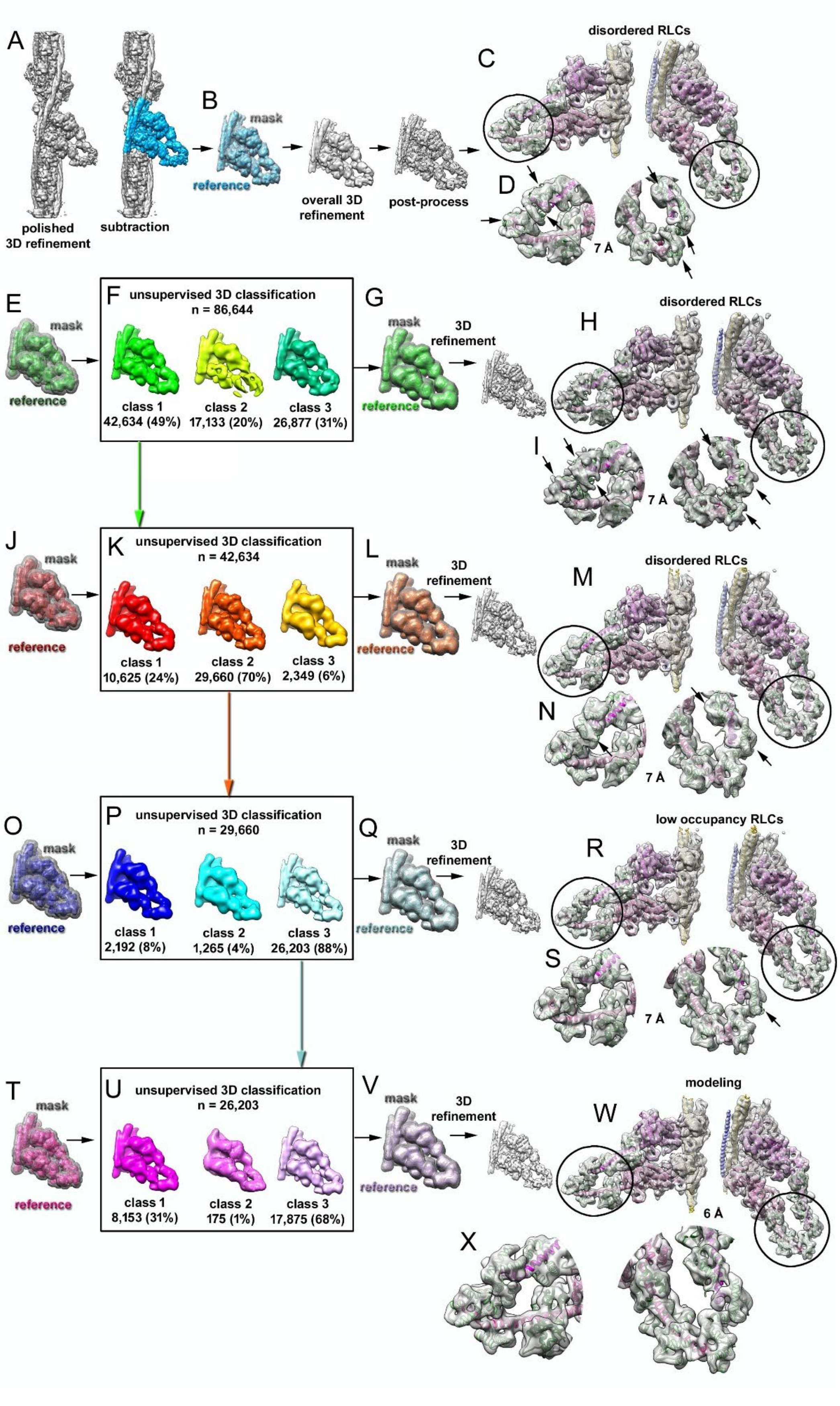

A
polished
3D refinement
subtraction
B
mask
reference
overall 3D
refinement
post-process
disordered RLCs
C
D
7 Å
E
mask
reference
F
unsupervised 3D classification
n = 86,644
class 1
42,634 (49%)
class 2
17,133 (20%)
class 3
26,877 (31%)
G
mask
reference
3D
refinement
disordered RLCs
H
I
7 Å
J
mask
reference
K
unsupervised 3D classification
n = 42,634
class 1
10,625 (24%)
class 2
29,660 (70%)
class 3
2,349 (6%)
L
mask
reference
3D
refinement
disordered RLCs
M
N
7 Å
O
mask
reference
P
unsupervised 3D classification
n = 29,660
class 1
2,192 (8%)
class 2
1,265 (4%)
class 3
26,203 (88%)
Q
mask
reference
3D
refinement
low occupancy RLCs
R
S
7 Å
T
mask
reference
U
unsupervised 3D classification
n = 26,203
class 1
8,153 (31%)
class 2
175 (1%)
class 3
17,875 (68%)
V
mask
reference
3D
refinement
modeling
W
6 Å
X

**Figure S18. 3D classification and reconstruction algorithm for the RLCs of HMM bound to actin molecules 6 and 7.** (A-D) Segmented densities of the HMM and actin molecules 6 and 7 were used for particle subtraction and then served as the reference (cerulean surface) and mask (gray mesh) for 3D refinement of those particles. The RLCs were not completely resolved (C-D, inserts and black arrows). (E-H) The map shown in (C) served as the reference and mask (green surface and gray mesh, respectively) for 3-class unsupervised sorting (F). Class 1 3D class-average (green surface) was used as the initial reference and mask (green surface and gray mesh) for 3D refinement (G), which showed that the RLCs were not completely resolved (H-I; inserts and black arrows). (J-M) The resultant map from (H) served as the reference and mask (J, dark red surface and gray mesh, respectively) to reclassify the selected segments from the first round (n = 42,634) into 3 classes using an unsupervised approach (K). Class 2 3D class average (dark orange surface) was used as the reference and mask (L, dark orange surface and gray mesh) for 3D refinement, which indicated that the RLCs were not completely resolved despite a noticeable improvement (inserts and black arrows). (O-R) The map from (M) served as the reference, and the mask (O, navy blue surface and gray mesh), along with class 2 segments (n = 29,660), were used for the next round of unsupervised sorting (P). From this, the Class 3 3D class average (light blue surface) was used as the initial reference and mask (Q, light blue surface and gray mesh) for 3D refinement. 3D refinement showed that the trailing RLC was still not completely resolved (R-S, inserts and black arrows). (T-X) The map from (R) served as the reference and mask (T, deep pink surface and gray mesh, respectively) for the final round of the unsupervised sorting (U). (V) Class 3 (n=17,895) 3D class average (light pink surface) was used as an initial reference and mask (light pink surface and gray mesh) for 3D refinement. Both RLCs on the trailing and leading heads were resolved and clearly showed their secondary structure (W-X, inserts, and black arrows).

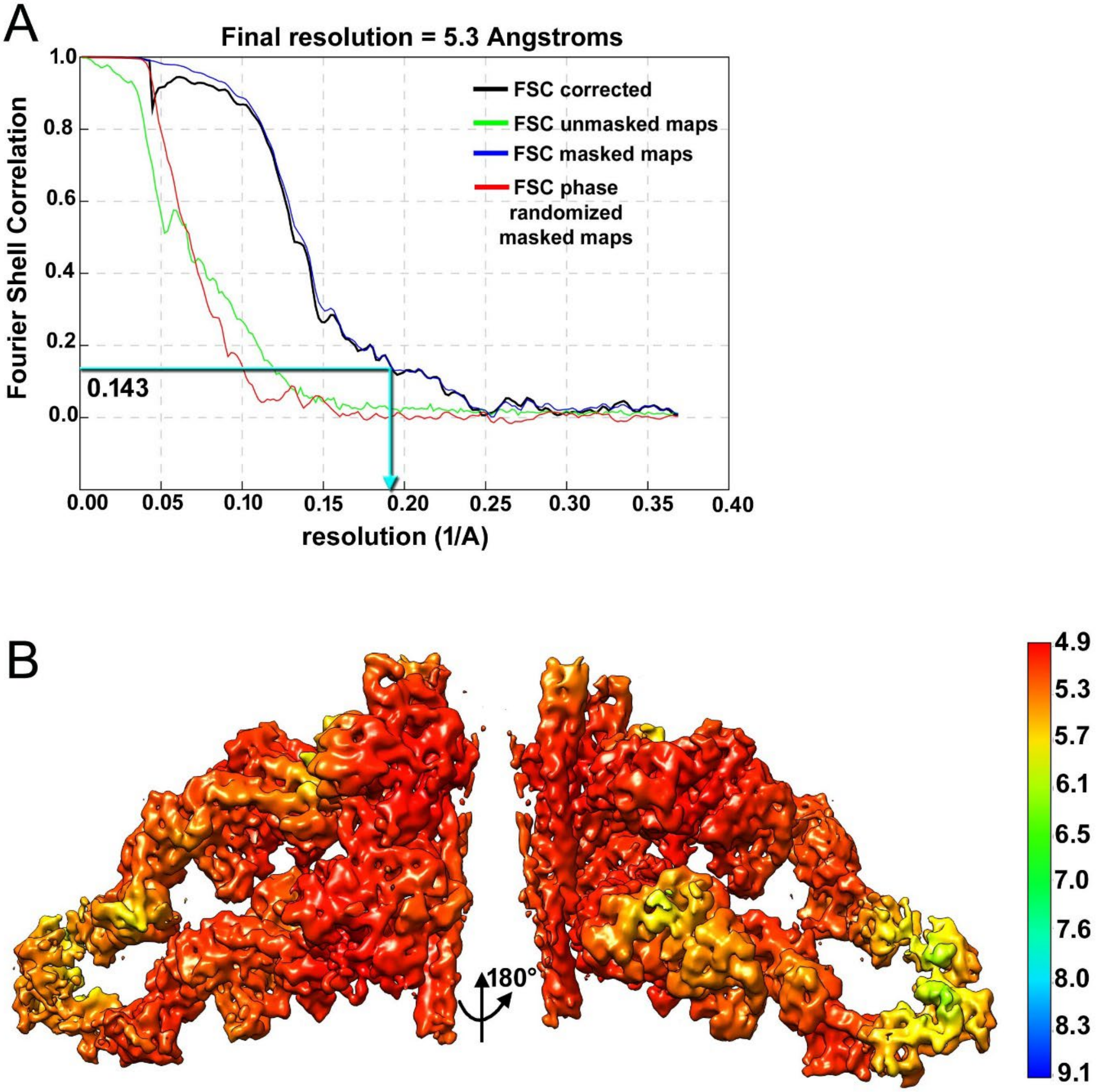


**Figure S19. Resolution estimation for the Class 6-7 map with RLCs.** (A) Fourier shell correlation (FSC) plot. The map's global resolution was calculated using an FSC criterion of 0.143. (B) Local resolution map determined by RELION.

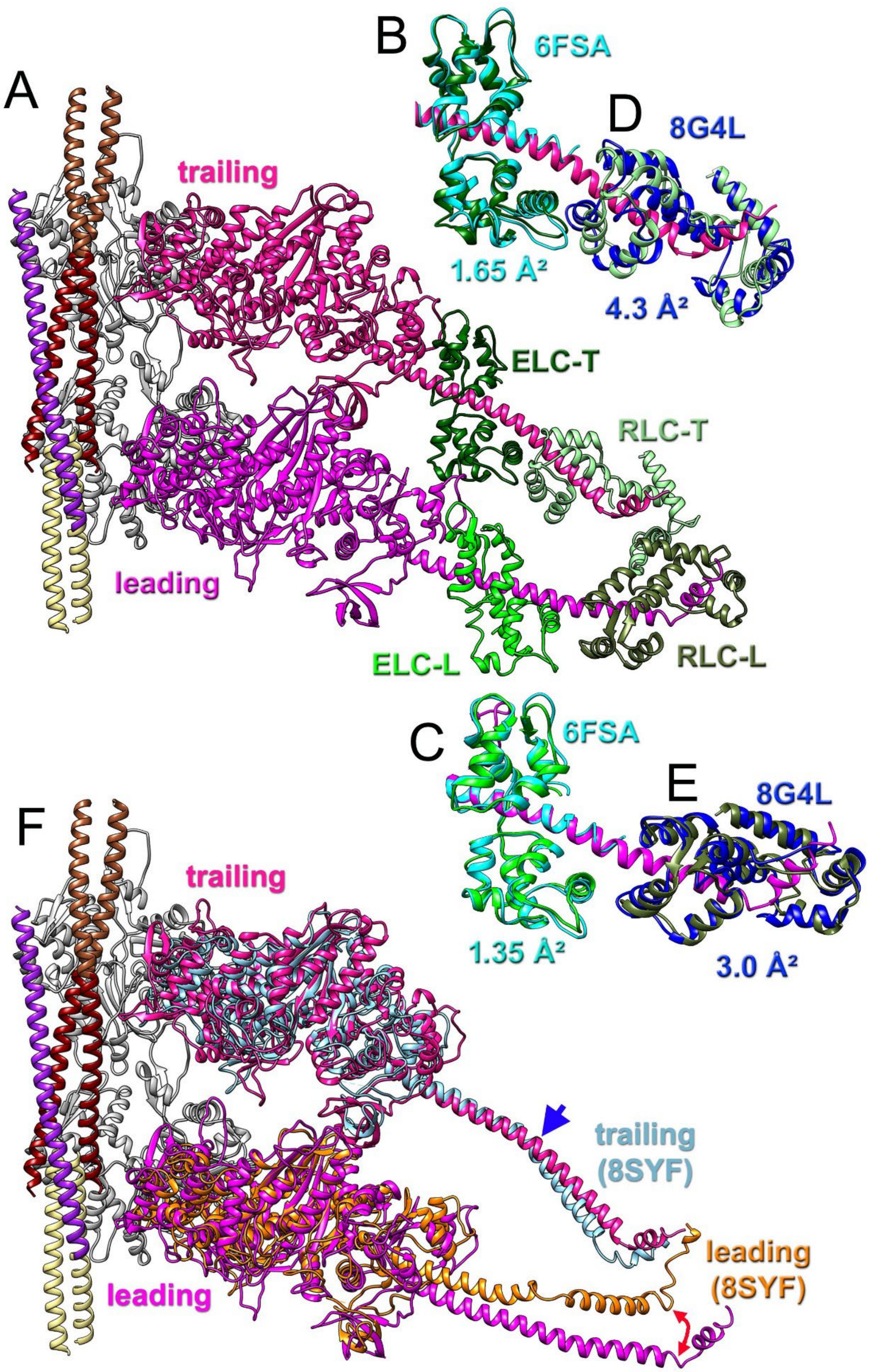

A
B
6FSA
D
8G4L
trailing
1.65 Å²
4.3 Å²
ELC-T
RLC-T
leading
ELC-L
RLC-L
C
6FSA
E
8G4L
F
trailing
1.35 Å²
3.0 Å²
trailing
(8SYF)
leading
(8SYF)
leading

**Figure S20. Comparison of ELC and RLC atomic models and rod-domain structural states for trailing and leading heads obtained in the current study with previously published structures.** (A) An overview of the atomic model of HMM bound to actins 6 and 7 that includes the neck region of myosin. The color codes are the same as in Figure 1. (B-E) Alignment of previously published structures of ELC and RLC with RMSD values provided for every light chain. (B and C) Both ELCs in our atomic model exhibit an excellent agreement with the high-resolution structure of cardiac myosin (PDB 6FSA) [21]. (D and E) Both RLCs are consistent with the structure of the cardiac RLC reported previously by cryo-EM of the inhibited human cardiac myosin thick filament (PDB 8G4L) [22]. (F) Structural alignment of the trailing (light blue) and leading (orange) heads from the cryo-EM structure of the HMM-ADP actomyosin complex (PDB 8SYF) [23] with our Class 6-7 structure, in which the trailing head rod domain is pink, and the leading head rod domain is magenta.

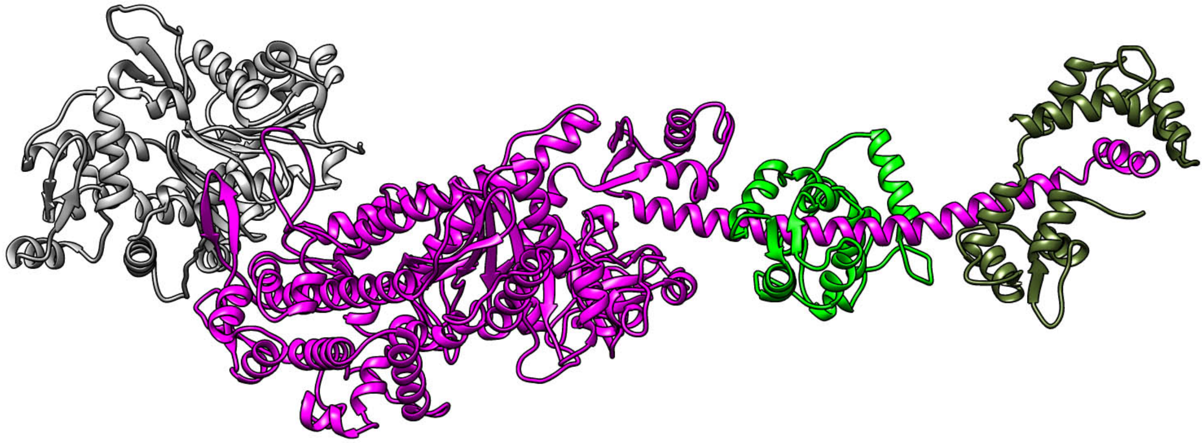


**Movie S1. Transition of the myosin tail between its positions in the leading and trailing heads.** The color code for the myosin molecule is the same as in Figure 1.

**Table S1. Data collection and refinement statistics.**

| **Classes** | HMM bound to actins 6-7 | HMM bound to actins 5-6 | HMM bound to actins 2-3 | HMM bound to actins 1-2 | HMM bound to actins 6-7 with RLC |
|---|---|---|---|---|---|
| **Data collection** | | | | | |
| Magnification | 65,000 | 65,000 | 65,000 | 65,000 | 65,000 |
| Defocus range, µm | 0.5 – 3.5 | 0.5 – 3.5 | 0.5 – 3.5 | 0.5 – 3.5 | 0.5 – 3.5 |
| Voltage, kV | 300 | 300 | 300 | 300 | 300 |
| Microscope | Titan Krios | Titan Krios | Titan Krios | Titan Krios | Titan Krios |
| Camera | K3 (super-resolution mode) | K3 (super-resolution mode) | K3 (super-resolution mode) | K3 (super-resolution mode) | K3 (super-resolution mode) |
| Number of frames | 40 | 40 | 40 | 40 | 40 |
| Total electron dose, $e^-/Å^2$ | 34 | 34 | 34 | 34 | 34 |
| Frames used for final reconstruction and dose used for final reconstruction, $e^-/Å^2$ | Determined by Relion MotionCorr internal implementation | Determined by Relion MotionCorr internal implementation | Determined by Relion MotionCorr internal implementation | Determined by Relion MotionCorr internal implementation | Determined by Relion MotionCorr internal implementation |
| Pixel size, Å/px | 0.678 | 0.678 | 0.678 | 0.678 | 0.678 |
| **Particle statistics** | | | | | |
| Particles | 86,644 | 55,229 | 31,180 | 54,451 | 17,875 |
| Box size, Å | 537 | 537 | 537 | 537 | 537 |
| Pixel size, Å/px | 1.356 | 1.356 | 1.356 | 1.356 | 1.356 |
| **Resolution FSC 0.143, Å** | 4.3 Å | 4.6 Å | 4.8 Å | 4.5 Å | 5.3 Å |

| | | | | | |
|---|---|---|---|---|---|
| **Resolution determined with Cryo-Res and local resolution maps used for modeling, Å** | 3.8 Å | 4.0 Å | 4.2 Å | 3.9 Å | 6 Å |

**Table S2. Atomic models validation.**

| **Classes** | HMM bound to actins 6-7 | HMM bound to actins 5-6 | HMM bound to actins 2-3 | HMM bound to actins 1-2 | HMM bound to actins 6-7 with RLC |
|---|---|---|---|---|---|
| **Atoms** | 30,836 | 33,975 | 33,217 | 30,685 | 26,682 |
| **MolProbity score** | 1.85 | 1.97 | 2.18 | 2.11 | 2.04 |
| **Clash score** | 6 | 6 | 7 | 7 | 8 |
| **Ramachandran plot (%)** | | | | | |
| Outliers | 0.00 | 0.07 | 0.15 | 0.13 | 0.00 |
| Allowed | 3.78 | 3.78 | 4.33 | 4.90 | 4.19 |
| Favored | 96.22 | 96.15 | 95.52 | 94.97 | 95.91 |
| **Rotamer outliers (%)** | 2.38 | 3.04 | 4.31 | 3.22 | 2.79 |
| **Model vs Data correlation** | 0.73 | 0.71 | 0.71 | 0.73 | 0.83 |
| **Deposition ID** | | | | | |
| PDB | 9YA8 | 9YAQ | 9YJP | 9YK9 | 9YKN |
| EMD | 72724 | 72734 | 73030 | 73042 | 73055 |

**Table S3. Missense cardiomyopathy-linked mutations related to the structure of the rigor cross-bridge.**

| *ACTC1* | | | | |
|---|---|---|---|---|
| Residue; porcine sequence number | Mutation; human sequence number | Disease | Role/location | Reference |
| D24 | D24N (D26N) | HCM | TnI binding | [24] |
| A26 | A26V (A28V) | HCM | CM-loop binding | [24] |
| K50 | K50T (K52T) | HCM | TnI binding | clinvar https://www.ncbi.nlm.nih.gov/clinvar/RCV001961500/ |
| Y91 | Y91C/H (Y93C/H) | HCM/LVNC | TnI binding | [25],[26] |
| R95 | R95C (R97C) | HCM | Loop-2 binding | [27] |
| E99 | E99K (E101K) | HCM | TnI binding | [28] |
| D222 | D222Y (D224Y) | LVNC | TmP-7 binding | [29] |
| D311 | D311H (D313H) | RCM | TmP-2, TmP-3, TmP-4 binding | [30] |
| *MYH7* | | | | |
| R369 | R369Q | Non-compaction CM | Tm binding | [31] |
| E374 | E374V | HCM | Tm-P2 binding | [24] |
| E379 | E379K | HCM | TnT binding | [32] |
| V404 | V404L/M | HCM | Actin binding | [33] ,[34] |
| V406 | V406M | HCM | Actin binding | [35] |
| V411 | V411I | HCM | Actin binding | [36] |
| E536 | E536D | HCM | Actin binding | [37] |
| F540 | F540L | DCM | Actin binding | [38] |
| K639 | K639E | LVNC | Actin binding | [39] |
| R723 | R723C/G/H | HCM | ELC binding | [40], [41], https://www.ncbi.nlm.nih.gov/clinvar/RCV000148962/ |
| R787 | R787C/H | HCM | ELC binding | [27], [42] |
| R793 | R793Q | HCM | ELC binding | [43] |

| | | | | |
|---|---|---|---|---|
| R807 | R807G/H | LVNC/HCM | RLC binding | [29], [25] |
| R819 | R819Q/W | HCM/DCM | RLC binding | [32], [24] |
| K835 | K835T | HCM | RLC binding | [44] |
| K837 | K837M/R | | RLC binding | https://www.ncbi.nlm.nih.gov/clinvar/RCV000467403/ https://www.ncbi.nlm.nih.gov/clinvar/RCV001808158/ |
| | | *MYL3* | | |
| E153 | E152K | HCM | Myosin converter interaction | [45] |
| | | *MYL2* | | |
| E22 | E22K | HCM | RLC-RLC interaction | [46] |
| R58 | E58Q/L | HCM | RLC-RLC interaction | [47], https://www.ncbi.nlm.nih.gov/clinvar/RCV000639677/ |
| E163 | E163A | HCM | Myosin rod interaction | https://www.ncbi.nlm.nih.gov/clinvar/RCV000036409/ |
| D166 | D166V | HCM | Myosin rod interaction | [42] |
| | | *TPM1* | | |
| E16 | E16Q | HCM | Actin binding | [24] |
| D20 | D20N | HCM | Actin binding | [24] |
| E23 | E23G | DCM | Actin binding | [48] |
| E54 | E54K | DCM | Loop-4 binding | [49] |
| D55 | D55N/Y | DCM | Loop-4 binding | [50], https://www.ncbi.nlm.nih.gov/clinvar/RCV000852460/ |
| D58 | D58H | HCM | Loop-4 binding | [51] |
| E96 | E96G | HCM | Loop-4 binding | https://www.ncbi.nlm.nih.gov/clinvar/RCV001996289/ |
| E98 | E98K | HCM | Loop-4 binding | [52] |
| E114 | E114G/Q | DCM | Actin binding | [50], [53] |
| I130 | I130T/V | HCM/ congenital heart defect | Loop-4 binding | https://www.ncbi.nlm.nih.gov/clinvar/RCV000549273/, [54] |

| | | | | |
|---|---|---|---|---|
| E180 | E180G/V/K | HCM | Actin binding | [55], [56], https://www.ncbi.nlm.nih.gov/clinvar/RCV001298950/ |
| S215 | S215L | HCM | Loop-4 binding | [27] |
| E250 | E250V | CM | Loop-4 binding | https://www.ncbi.nlm.nih.gov/clinvar/RCV000852462/ |
| E254 | E254G | HCM | Loop-4 binding | [24] |
| *TNNI3* | | | | |
| K37 | K36Q | DCM | TnC interaction | [57] |
| D128 | D127Y | RCM/HCM | Actin binding | [58] |
| R147 | R146S | HCM | Actin binding | https://www.ncbi.nlm.nih.gov/clinvar/RCV001256923/ |
| TNNT2 | | | | |
| R91 | R92L/Q/W | HCM | Loop-4 binding | [59], [55], [60], [61] |
| R202 | R205L/W | DCM | Actin binding | [62], [63] |
| N266 | N269D/K | HCM | Loop-2 binding | [24] [37] |

mutations in the ventricular regulatory myosin light chain gene (MYL2) associated with familial and classical forms of hypertrophic cardiomyopathy. J Mol Med (Berl). **76**, 208-214. https://doi.org/10.1007/s001090050210.

48. Hershberger, R. E., Norton, N., Morales, A., Li, D., Siegfried, J. D. & Gonzalez-Quintana, J. (2010). Coding sequence rare variants identified in MYBPC3, MYH6, TPM1, TNNC1, and TNNI3 from 312 patients with familial or idiopathic dilated cardiomyopathy. Circ Cardiovasc Genet. **3**, 155-161. https://doi.org/10.1161/CIRCGENETICS.109.912345.
49. Olson, T. M., Kishimoto, N. Y., Whitby, F. G. & Michels, V. V. (2001). Mutations that alter the surface charge of alpha-tropomyosin are associated with dilated cardiomyopathy. J Mol Cell Cardiol. **33**, 723-732. https://doi.org/10.1006/jmcc.2000.1339.
50. Pugh, T. J., Kelly, M. A., Gowrisankar, S., Hynes, E., Seidman, M. A., Baxter, S. M., Bowser, M., Harrison, B., Aaron, D., Mahanta, L. M., Lakdawala, N. K., McDermott, G., White, E. T., Rehm, H. L., Lebo, M. & Funke, B. H. (2014). The landscape of genetic variation in dilated cardiomyopathy as surveyed by clinical DNA sequencing. Genet Med. **16**, 601-608. https://doi.org/10.1038/gim.2013.204.
51. Frisso, G., Limongelli, G., Pacileo, G., Del Giudice, A., Forgione, L., Calabro, P., Iacomino, M., Detta, N., Di Fonzo, L. M., Maddaloni, V., Calabro, R. & Salvatore, F. (2009). A child cohort study from southern Italy enlarges the genetic spectrum of hypertrophic cardiomyopathy. Clin Genet. **76**, 91-101. https://doi.org/10.1111/j.1399-0004.2009.01190.x.
52. Matyushenko, A. M., Nefedova, V. V., Kochurova, A. M., Kopylova, G. V., Koubassova, N. A., Shestak, A. G., Yampolskaya, D. S., Shchepkin, D. V., Kleymenov, S. Y., Ryabkova, N. S., Katrukha, I. A., Bershitsky, S. Y., Zaklyazminskaya, E. V., Tsaturyan, A. K. & Levitsky, D. I. (2023). Novel Mutation Glu98Lys in Cardiac Tropomyosin Alters Its Structure and Impairs Myocardial Relaxation. Int J Mol Sci. **24**. https://doi.org/10.3390/ijms241512359.
53. Man, Y., Yi, C., Fan, M., Yang, T., Liu, P., Liu, S. & Wang, G. (2022). Identification of a novel missense mutation in the TPM1 gene via exome sequencing in a Chinese family with dilated cardiomyopathy: A case report and literature review. Medicine (Baltimore). **101**, e28551. https://doi.org/10.1097/MD.0000000000028551.
54. England, J., Granados-Riveron, J., Polo-Parada, L., Kuriakose, D., Moore, C., Brook, J. D., Rutland, C. S., Setchfield, K., Gell, C., Ghosh, T. K., Bu'Lock, F., Thornborough, C., Ehler, E. & Loughna, S. (2017). Tropomyosin 1: Multiple roles in the developing heart and in the formation of congenital heart defects. J Mol Cell Cardiol. **106**, 1-13. https://doi.org/10.1016/j.yjmcc.2017.03.006.
55. Thierfelder, L., Watkins, H., MacRae, C., Lamas, R., McKenna, W., Vosberg, H. P., Seidman, J. G. & Seidman, C. E. (1994). Alpha-tropomyosin and cardiac troponin T mutations cause familial hypertrophic cardiomyopathy: a disease of the sarcomere. Cell. **77**, 701-712. https://doi.org/10.1016/0092-8674(94)90054-x.
56. Regitz-Zagrosek, V., Erdmann, J., Wellnhofer, E., Raible, J. & Fleck, E. (2000). Novel mutation in the alpha-tropomyosin gene and transition from hypertrophic to hypocontractile dilated cardiomyopathy. Circulation. **102**, E112-116. https://doi.org/10.1161/01.cir.102.17.e112.
57. Carballo, S., Robinson, P., Otway, R., Fatkin, D., Jongbloed, J. D., de Jonge, N., Blair, E., van Tintelen, J. P., Redwood, C. & Watkins, H. (2009). Identification and functional characterization of cardiac troponin I as a novel disease gene in autosomal dominant dilated cardiomyopathy. Circ Res. **105**, 375-382. https://doi.org/10.1161/CIRCRESAHA.109.196055.
58. Hassoun, R., Budde, H., Mannherz, H. G., Lodi, M., Fujita-Becker, S., Laser, K. T., Gartner, A., Klingel, K., Mohner, D., Stehle, R., Sultana, I., Schaaf, T., Majchrzak, M., Krause, V., Herrmann, C., Nowaczyk, M. M., Mugge, A., Pfitzer, G., Schroder, R. R., Hamdani, N.,

Milting, H., Jaquet, K. & Cimiotti, D. (2021). De Novo Missense Mutations in TNNC1 and TNNI3 Causing Severe Infantile Cardiomyopathy Affect Myofilament Structure and Function and Are Modulated by Troponin Targeting Agents. Int J Mol Sci. **22**. https://doi.org/10.3390/ijms22179625.
