## Supplementary figures and images for "The structure of the native cardiac crossbridge in the rigor state"

### Supplimental movie

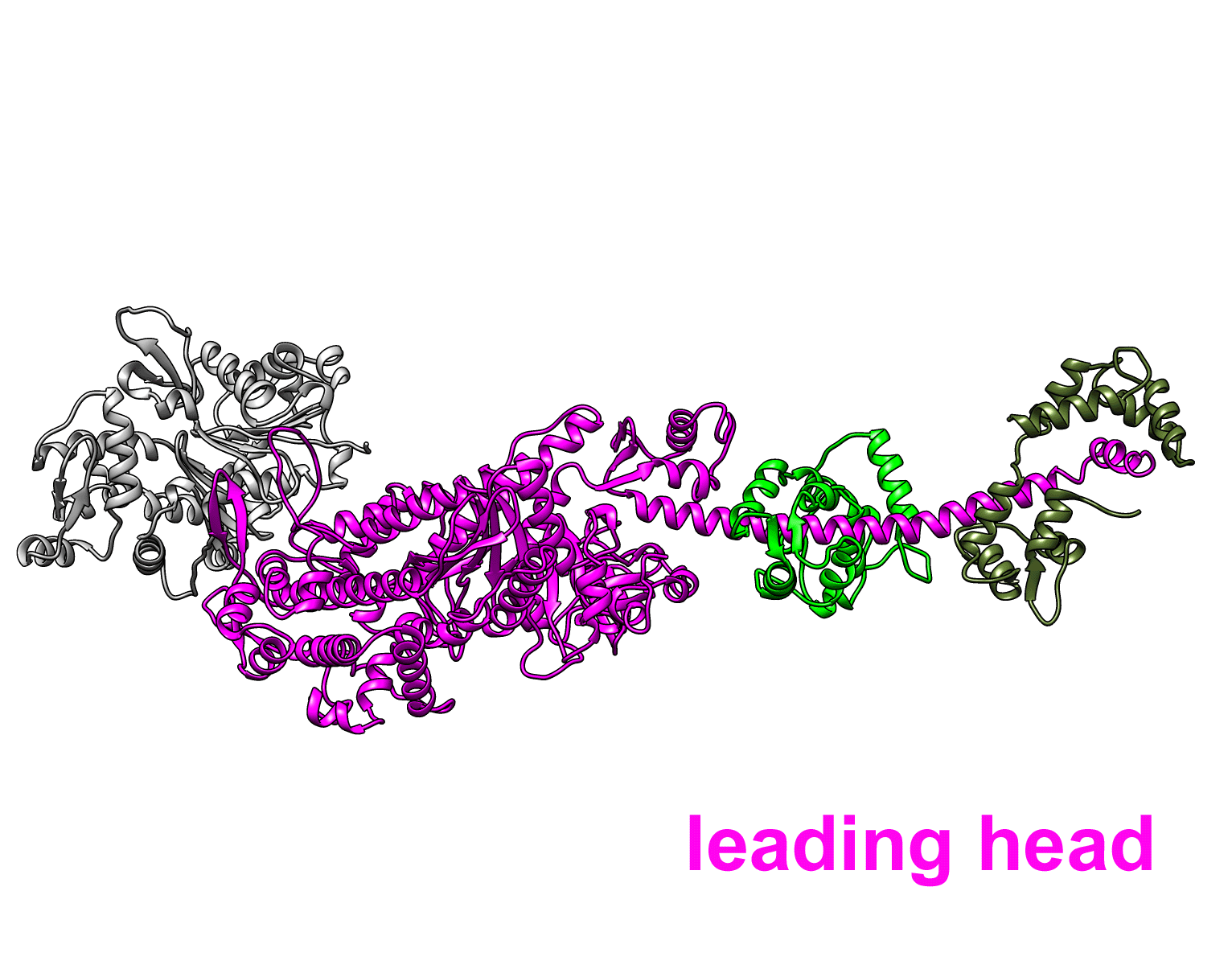
